# Hyperexcitable primary somatosensory cortex in the MitoPark mouse model of Parkinson’s disease

**DOI:** 10.64898/2026.09.29.755142

**Authors:** Ronghao Zhang, Ruorong Qi, Alan J. Emanuel

**Affiliations:** Department of Cell Biology, Emory University School of Medicine, Atlanta, GA 30322, USA; Wallace H Coulter Department of Biomedical Engineering, Georgia Institute of Technology and Emory University, Atlanta, GA 30332, USA

## Abstract

The majority of Parkinson’s disease patients suffer from decreased tactile acuity or neuropathic pain, both of which contribute to low quality of life and likely exacerbate motor dysfunction. How Parkinson’s disease pathology relates to altered somatosensation remains unclear. One potential contributor is the progressive degeneration of dopamine neurons. To assess the extent to which gradual dopamine neuron loss contributes to altered somatosensation, we recorded spontaneous and stimulus-evoked activity in the primary somatosensory cortex of the MitoPark mouse model of Parkinson’s disease. MitoPark mice experience progressive dopamine neuron degeneration that is independent of α-synuclein. We found that MitoPark layer V regular-spiking (largely pyramidal) neurons, but not fast-spiking (largely parvalbumin-expressing inhibitory) neurons, had elevated spontaneous firing rates relative to littermate controls at late stages of symptom progression. Regular-spiking neurons also became hyperresponsive to tactile stimuli – more neurons were recruited by stimuli across a wide range of intensities, and these neurons responded with higher firing rates. Fast-spiking neurons, but not regular-spiking neurons, became desynchronized at late stages of symptom progression. These physiological changes coincided with pronounced degeneration of dopamine neurons and axons, which we measured with tyrosine hydroxylase immunostaining in the midbrain, striatum, and primary somatosensory cortex. Altogether, this characterization revealed cell-type-specific changes in the primary somatosensory cortex that are consistent with both decreased tactile acuity and hyperalgesia in Parkinson’s disease patients. This suggests that the loss of dopamine neurons, potentially including those that innervate primary somatosensory cortex, contributes to somatosensory impairments associated with Parkinson’s disease.

**Significance Statement:** Due to the extensive coordination of sensory inputs and motor controls, impaired somatosensory processing in Parkinson’s Disease (PD) patients likely exacerbates motor dysfunction and affects execution of daily tasks. We report that the primary somatosensory cortex of the MitoPark mouse model of Parkinson’s disease becomes hyperactive and hyperresponsive, which suggests that the loss of dopaminergic signaling directly influences sensory processing. This contributes to the understanding of the mechanisms underlying tactile dysfunction in PD, which should lead to better therapeutics for sensorimotor function in patients, potentially through optimization of non-invasive neuromodulation of the somatosensory system.

## Introduction

Parkinson’s disease (PD) is associated with impaired somatosensation in up to 85% of patients (1). In the touch domain, PD patients exhibit diminished spatial and temporal tactile discrimination, reflecting reduced tactile acuity (2–4) and the gain of pressure perception in PD patients is abnormally inflexible relative to age-matched controls (5). PD is also frequently accompanied by neuropathic pain (6), marked by chronic spontaneous pain as well as heightened sensitivity to mechanical and thermal stimulation (7, 8), alongside nociceptive and nociplastic pain syndromes (9, 10). Given the tight coupling between sensory input and motor output, these somatosensory abnormalities in PD likely compound motor dysfunction and further disrupt the execution of daily tasks (11–13). Yet how PD pathology relates to altered somatosensation is unresolved, and whether tactile dysfunction and pain abnormalities in PD share a common origin remains without consensus.

Proposed mechanisms for these sensory abnormalities include peripheral neuropathy (14), accumulation of α-synuclein in Lewy bodies and neurites (15), medication effects (16, 17), circuit changes within somatosensory and nociceptive pathways (18), or loss of dopamine (DA) neurons (2). Among these, DA degeneration is a strong candidate driver for sensory aberration in PD. Firstly, the progressive loss of midbrain DA neurons, from both substantia nigra pars compacta (SNpc) and ventral tegmental area (VTA), is a key pathological hallmark of PD (19–22). Secondly, PD-related pain tends to emerge during periods of low DA tone (23), and deficits in tactile submodalities can be partially alleviated by DA replacement (24). However, it remains unclear how DA loss shapes the neural representation of tactile signals in PD.

Mouse models enable the dissection of how pathology relates to neural processing. The MitoPark mouse model of PD (*DAT^Cre^;Tfam^flox/flox^*) isolates DA degeneration from other pathological hallmarks through genetically induced mitochondrial dysfunction in DA transporter (DAT)-expressing cells (25, 26). MitoPark mice exhibit relatively stereotyped progressive loss of DA neurons alongside gradual escalation of motor symptoms that parallel the gradual DA loss and symptom progression of human PD patients (27, 28). Parkinsonism in MitoPark mice is similar whether or not α-synuclein is knocked out (25), indicating that DA depletion and compensation thereof are the major phenotypic drivers. Unlike acute toxin-based PD models (29, 30), progressive DA depletion in MitoPark mice allows longitudinal tracking of symptoms across the course of disease. MitoPark mice present a number of non-motor symptoms, including early impairments in olfactory discrimination (31, 32), disrupted circadian rhythms (33), altered cognitive learning and memory (32, 34), and anxiety-like behaviors (32), though measurements of somatosensory capabilities have not been reported. Therefore, we sought to characterize somatosensory signaling in this mouse model across timepoints that span motor symptom progression.

A limitation for the study of somatosensation in PD models is that behavioral tests often conflate sensory and motor functions and expected motor deficits may obscure changes to sensory function. To isolate the effects of DA degeneration on tactile processing, we therefore directly assessed spontaneous and touch-evoked responses in forepaw primary somatosensory cortex (S1), the first cortical processing center important for both discriminative touch and pain. By leveraging precise control over stimulus delivery and high-density recordings of S1 activity (35), we found that hyperexcitability and increased responsiveness is a prominent phenotype in advanced-age MitoPark mice, implicating DA degeneration as a driver of impaired somatosensation in PD.

## Results

### Degeneration of TH^+^ axons in MitoPark S1 occurs concurrently with degeneration in the striatum and midbrain

Degeneration of DA neurons in MitoPark animals varies in timing and extent across neuron populations. Midbrain DA neurons in the substantia nigra pars compacta (SNpc) degenerate faster than those of the ventral tegmental area (VTA), likely due to an especially high calcium load and redox imbalance (25, 36), and midbrain DA neurons in general are more strongly affected than those in the olfactory bulb, which express DA transporter (DAT) at later ages (31). To determine whether the progressive degeneration of DA neurons in MitoPark mice extends to those that innervate S1, we stained brain tissue from control and MitoPark animals at 12 (± 1), 15 (± 1), and 26 (± 3) weeks of age with a tyrosine hydroxylase (TH) antibody that labels catecholaminergic neurons. We quantified TH^+^ fiber density in S1 **(Figure S1)** as well as the intensity of TH staining in the dorsolateral striatum (dlSTR) and midbrain. These timepoints span disease progression from pre-symptomatic stages to the development of overt motor deficits **(Table S1)** (27). TH intensity in both dlSTR and midbrain showed a significant genotype × age interaction (ANOVA). Post-hoc pairwise comparisons between genotypes at each timepoint revealed a significant decrease in midbrain TH intensity in MitoPark mice compared to controls at both 15 and 26 weeks (15 weeks: t = −2.87, p = 0.042; 26 weeks: t = −6.68, p < 0.0001, Bonferroni corrected for multiple comparisons). MitoPark mice exhibited decreased TH intensity in dlSTR compared to controls at 26 weeks (t = −3.52, p = 0.012) **(Fig. 1A, C)**. TH^+^ fibers in S1 were also affected **(Fig. 1B, D)**. We found a significant main effect of genotype in all layers of S1 (ANOVA), with post-hoc comparisons showing a significant deficit specifically at 26 weeks for most S1 layers (layer I: t = −2.50, p < 0.067; layer II/III: t = −3.42, p = 0.00925; layer IV: t = −3.34, p = 0.011; layer V: t = −3.84, p = 0.00361; layer VI: t = −3.69, p = 0.00498). Together, these results indicate that TH^+^ axon loss in S1 in MitoPark mice progresses in parallel with midbrain and striatal TH loss. Because *DAT^Cre^* is not expected to recombine in the locus coeruleus and cortical norepinephrine is at normal levels at this timepoint in MitoPark mice (28), this reduction likely reflects a loss of DA fibers.

**Figure 1.**
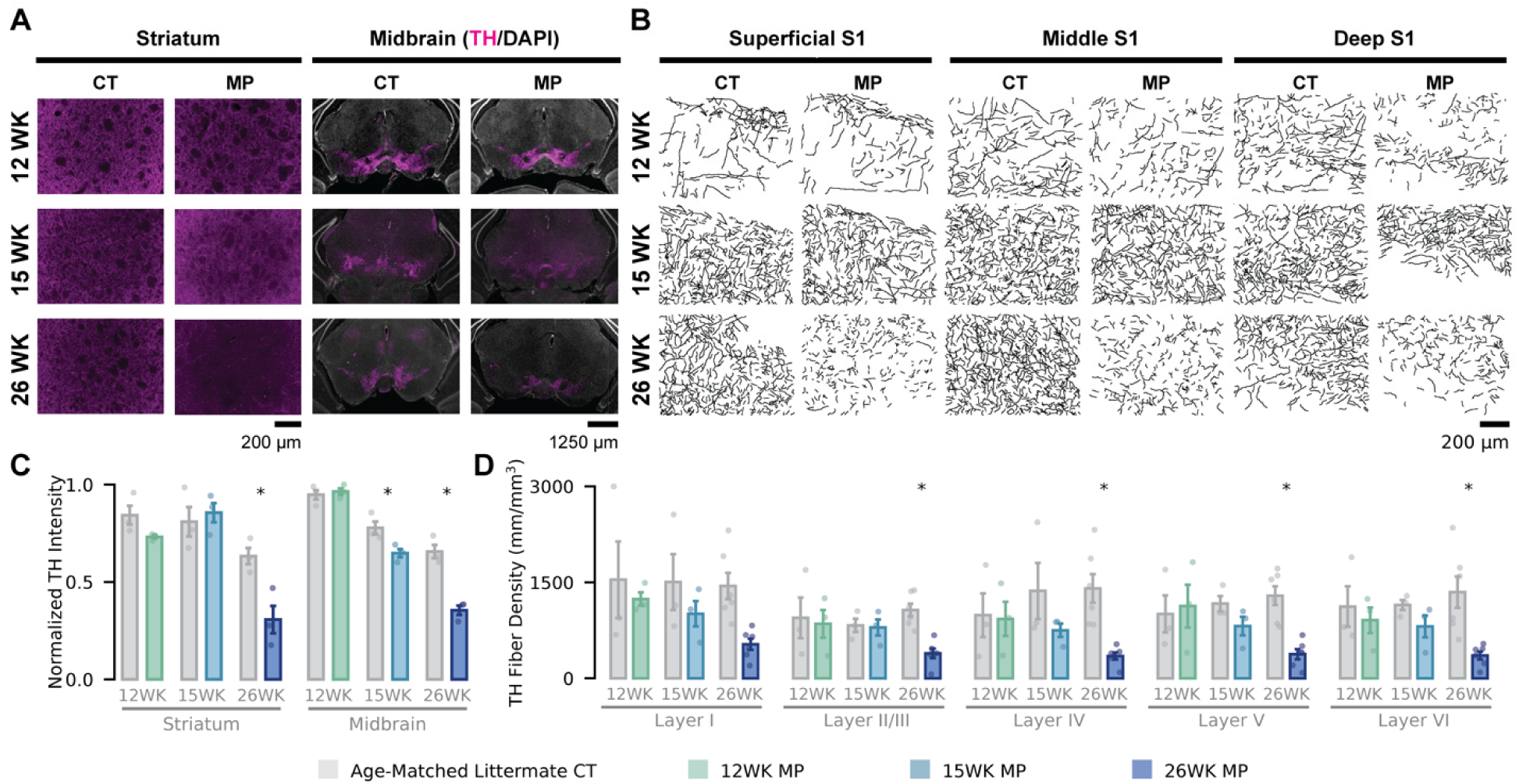
Progressive loss of midbrain TH^+^ neurons and TH^+^ axons in striatum and S1 of MitoPark mice. **(A)** Representative TH immunofluorescence images in dorsolateral striatum (dlSTR) and midbrain of control (CT) and MitoPark (MP) mice at 12, 15, and 26 weeks. WK: weeks. **(B)** Representative reconstructions of TH^+^ axons in superficial, middle, and deep layers of S1 in control and MitoPark mice at each timepoint. **(C)** TH staining intensity (mean ± S.E.M.; n = 3 mice per group) in dlSTR and midbrain of control and MitoPark mice across three time points, normalized to maximum. **(D)** TH^+^ fiber density (mean ± S.E.M.; n = 3 mice per genotype for 12-week and 15-week timepoints, n = 6 for 26-week timepoint) within each cortical layer of S1 across timepoints. *p<0.05 for comparison between age-matched control and MitoPark mice, post hoc simple effect contrasts with Bonferroni correction for multiple comparisons. WK: week; CT: control mice; MP: MitoPark mice.

### Regular-spiking S1 units exhibit increased excitability in advanced-age MitoPark mice

The depletion of TH^+^ axons in cortical and subcortical structures raised the question of whether MitoPark mice exhibit functional changes in S1 activity. We used multielectrode arrays to record from awake, head-fixed mice while stimulating the pedal pads of the plantar surface of the tethered forepaw **(Fig. 2A)**. We targeted the S1 region where neuronal receptive fields of forepaw glabrous skin are concentrated. In total, we recorded 3,906 cortical single units in 84 recording sessions of 24 control and 21 MitoPark mice across three timepoints **(Table S1 Lower Panel)**. Units were classified as fast-spiking (FS, largely parvalbumin-expressing interneurons; n = 1,209) or regular-spiking (RS, largely excitatory; n = 2,697) based on the trough-to-peak interval of spike extracellular waveforms **(Fig. 2B)** (37, 38). Single-unit yield and the fractions of FS units were largely comparable between groups **(Figs. S2A,B)**, though we observed a decreased fraction of sparse-firing RS units in 26-week MitoPark mice relative to age-matched controls (t = −2.553, p = 0.032, post hoc simple effect contrasts between genotypes after linear mixed-effects model [LME] with Bonferroni correction for multiple comparisons) **(Fig. S2C)**. To characterize changes in spiking properties in each layer of the cortex across disease progression, we assigned units to cortical layers using depth measurements calibrated against the current-source density map derived from local field potentials **(Fig. 2C)** (35). We computed baseline firing rates (mean firing rate in a 0.8-s pre-stimulus window, averaged across all trials) and evoked firing rates (peak baseline-subtracted, trial-averaged firing rate across all time bins and stimulus intensities) of each unit **(Fig. 2D)**. Layer V RS units of 15- and 26-week MitoPark mice had elevated baseline firing rates relative to those of age-matched controls, and we did not detect differences in RS units of other layers **(Fig. 2E)**. The baseline firing rates of FS units in MitoPark mice did not differ from those of control mice in any cortical layer **(Fig. 2F)**. We observed a similar pattern in the evoked firing rates. Layer V RS neurons of MitoPark mice had higher evoked firing rates relative to those of control mice at 26 weeks **(Fig. 2G)**, and evoked firing rates did not differ between MitoPark and control FS units in any layer at any timepoint **(Fig. 2H)**. Notably, both baseline and evoked firing rates of RS and FS units from 12-week MitoPark mice were comparable to those of units from age-matched controls. Together, these results reveal a layer- and cell-type-specific hyperexcitability in S1 neurons that emerges around 15 weeks and intensifies by 26 weeks, affecting both spontaneous and sensory-evoked activity.

**Figure 2.**
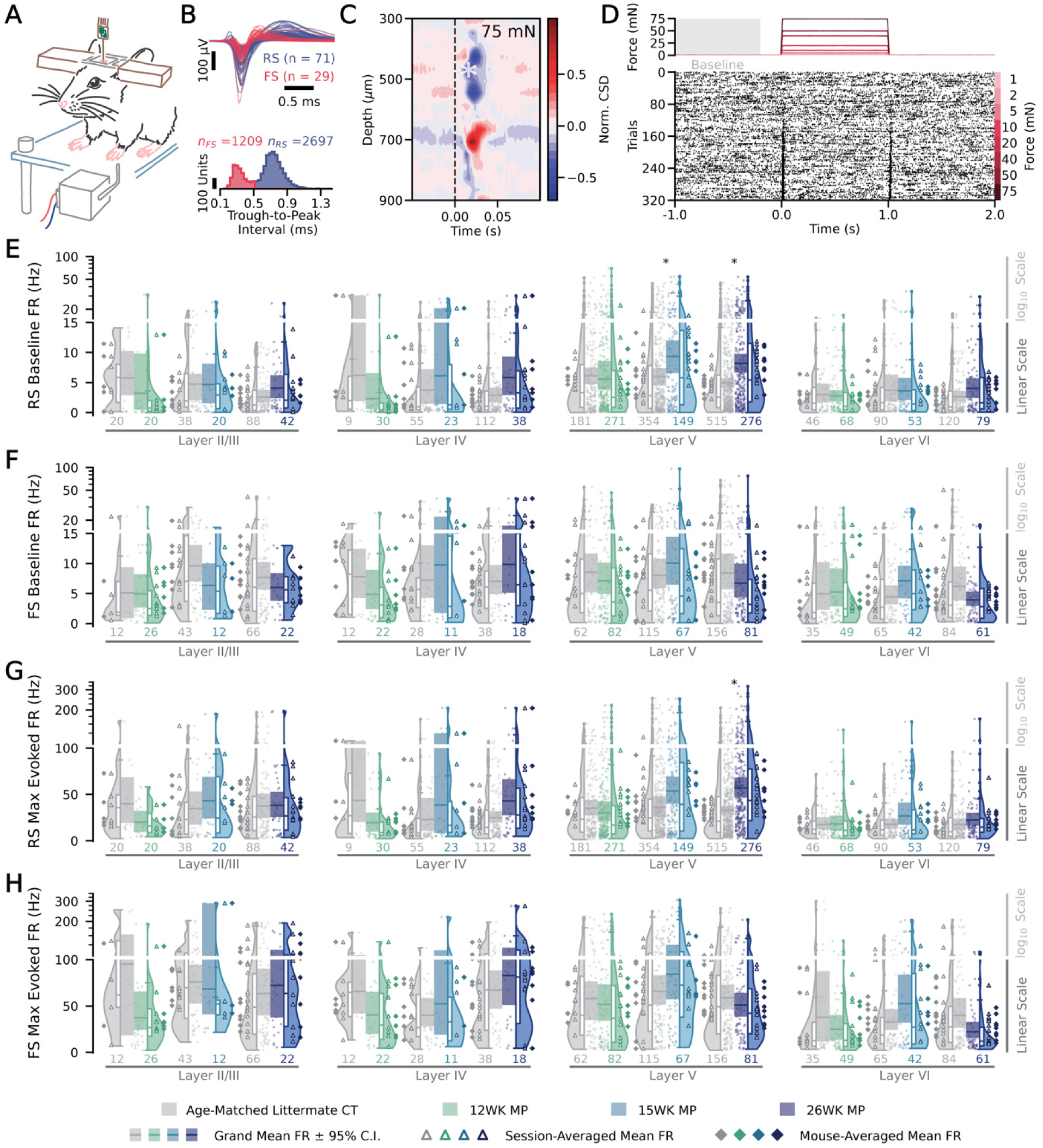
MitoPark S1 layer V regular-spiking units exhibit elevated baseline and sensory-evoked firing rates. **(A)** Schematic of the experimental paradigm. **(B) Top:** mean extracellular waveforms from 100 randomly selected single units recorded with multielectrode arrays. **Bottom:** Distribution of trough-to-peak intervals for all units (n_total_ = 3906, n_FS_ = 1209, n_RS_ = 2697). **(C)** Exemplar current-source density (CSD) map aligned to the onset (black dashed line) of 75 mN stimuli in recording of a control mouse. Depth aligned to the dura. Sources (red) and sinks (blue) appear soon after the onset of the stimuli. White asterisk (*) marks an early sink, corresponding to layer IV (475 µm ventral to the dura). **(D)** Exemplar raster plot of the action potentials aligned to step indentation of an FS unit. Trials sorted by stimulus intensity. Gray window indicates the baseline period. **(E)** Baseline firing rates of RS units, separated by cortical layers, genotypes, and ages. Points denote firing rates for individual units, with solid horizontal bars representing the grand mean firing rate of the group. Colored boxes indicate the 95% C.I. of the grand mean firing rate (hierarchical bootstrap). Violin plots depict the full distribution of data, with overlaid box plots (median and interquartile range) summarizing dispersion. Hollow triangles indicate session-averaged mean firing rates, and solid diamonds indicate animal-averaged mean firing rates. The number of units recorded within each group is shown at the bottom. **(F)** Similar to (E), but for baseline firing rates of FS units. **(G)** Similar to (E), but for baseline-subtracted maximum sensory-evoked firing rates of RS units. **(H)** Similar to (G), but for FS units. *p<0.05, post hoc simple-effects contrast test for genotype at each sampled age with Bonferroni correction for multiple comparisons. RS: regular-spiking units; FS: fast-spiking units; WK: week; CT: control mice; MP: MitoPark mice.

### Regular-spiking S1 units of advanced-age MitoPark mice are hyperresponsive to tactile stimulation

We next asked how MitoPark S1 neurons encode stimulus intensity. To do so, we extracted single-unit responses to each of the eight indentation intensities and computed mean z-scored firing rates **(Fig. S2D)**. As expected (35), most S1 RS units produced transient responses at the onset (ON) and offset (OFF) of step indentations at intensities as low as 5 mN. These responses grew with stronger forces until they saturated. Only a small subset of neurons responded during the sustained phase of high-force indentations **(Figs. 3A-C)**. Comparing across genotypes, we observed a more robust response from 26-week MitoPark RS units during the ON and OFF periods of the 5-mN step compared to control RS units **(Fig. 3C)**.

**Figure 3.**
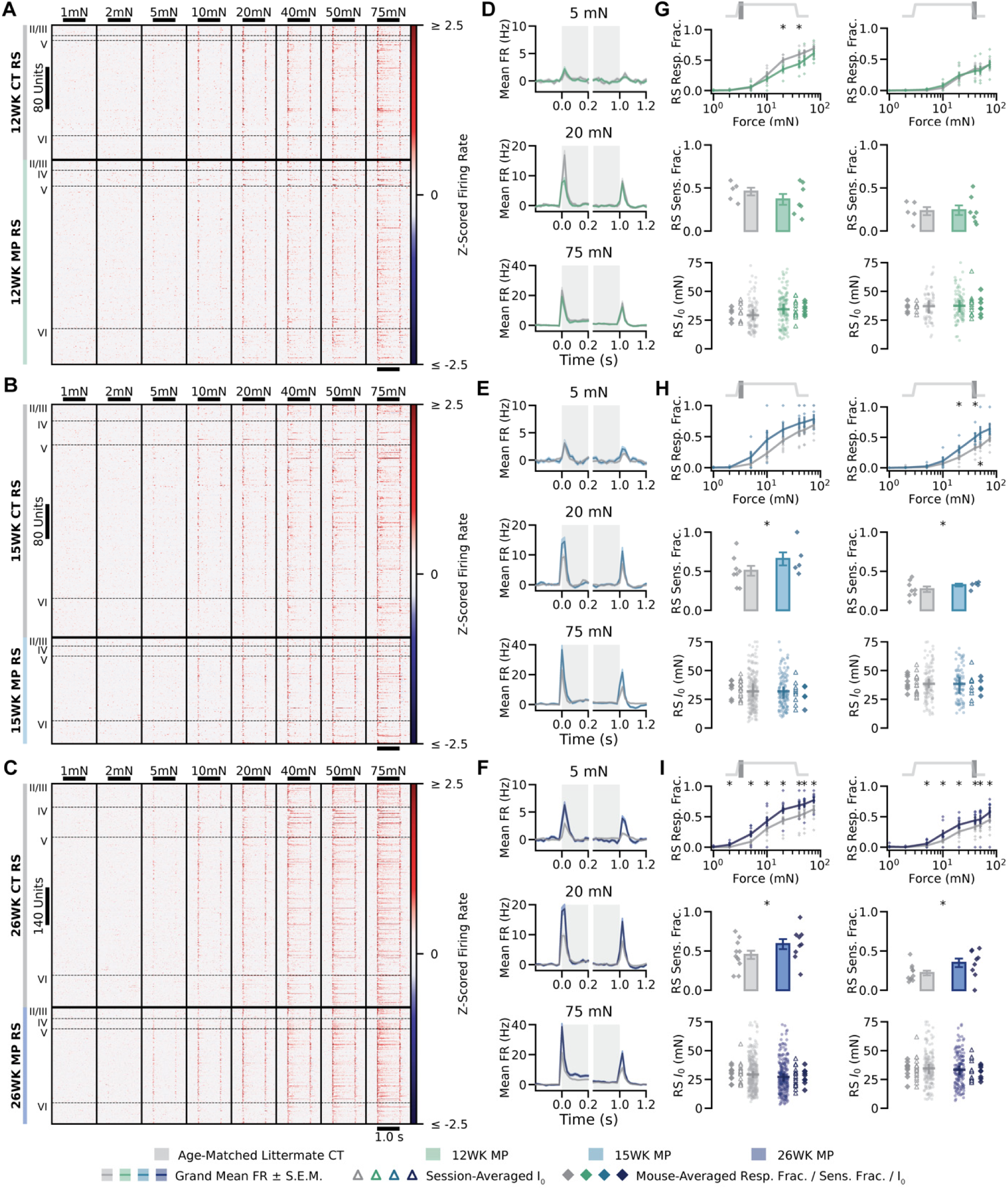
Tactile stimuli recruit increased proportions of S1 regular-spiking units in MitoPark mice at advanced ages. **(A)** Depth-sorted indentation-evoked responses from regular-spiking (RS) S1 units in 12-week control and MitoPark mice, with animal genotypes indicated by the solid black divider and cortical layers by dashed lines. WK; weeks; MP: MitoPark mice; CT: control mice. **(B)** Similar to (A), but for comparisons of 15-week control and MitoPark RS units. **(C)** Similar to (A), but for comparisons of 26-week control and MitoPark RS units. **(D)** Grand-mean (± S.E.M.) evoked firing rates across all cortical RS units of 12-week MitoPark (blue) and age-matched control (gray) in response to 5 mN (upper panel), 20 mN (middle panel), and 75 mN indentation (lower panel). Gray shading indicates stimulus presence. **(E)** Similar to (D), but for comparisons of 15-week control and MitoPark RS units. **(F)** Similar to (D), but for comparisons of 26-week control and MitoPark RS units. **(G) Top:** Fraction (mean ± S.E.M.) of RS neurons responsive to the ON **(left)** and OFF **(right)** period of indentations in 12-week control and 12-week MitoPark mice. **Middle:** Fraction (mean ± S.E.M) of RS neuron ON (left) and OFF (right) responses that were well-fitted by a saturating exponential function, 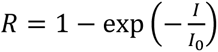, where *R* is the mean firing rates at each intensity, *I* is the intensity of force steps, and *I*_0_ is the fitted parameter indicating the sensitivity threshold of this single unit. Bottom: *I*_0_ distribution (mean ± 95% C.I.; hierarchical bootstrap) for ON **(left)** and OFF **(right)** responses of well-fitted RS neurons recorded from 12-week control and MitoPark mice. Triangles represent the mean *I*_0_ of individual sessions, while diamonds represent the mean *I*_0_ of individual animals. **(H)** Similar to (G), but for comparisons of 15-week control and MitoPark RS units. **(I)** Similar to (G), but for comparisons of 26-week control and MitoPark RS units. *p<0.05 for comparisons of the mean fraction between age-matched control and MitoPark mice, two-proportions z-tests with Bonferroni correction for multiple comparisons. FR: firing rates; RS: regular-spiking units; FS: fast-spiking units; WK: week; CT: control mice; MP: MitoPark mice.

Grand-mean peristimulus time histograms (PSTHs) at three representative forces (5, 20, 75 mN) revealed that 12-week MitoPark RS units tended to have decreased firing rates compared to age-matched controls whereas 15- and 26-week MitoPark RS units tended to have increased firing rates **(Figs. 3D-F)**. In addition to increased single-unit firing rates, a change in the population response could be driven by recruitment of additional S1 neurons. To measure this, we calculated the fraction of RS units responsive to each indentation force during the ON and OFF periods. At 12 weeks, the fraction of responsive RS units in MitoPark mice was slightly higher than in control mice at intermediate intensities during the ON period (20 mN: z = −3.90, p < 0.005; 40 mN: z = −3.70, p = 0.01, two-proportion z-test with Bonferroni correction for multiple comparisons), but no significant difference was observed during the OFF period **(Fig. 3G upper panels)**. However, the trend was reversed at 15 weeks **(Fig. 3H upper panels)** and the difference between genotypes became more pronounced by 26 weeks **(Fig. 3I upper panels)**, when MitoPark mice had significantly elevated response fractions across a broad range of forces during both ON and OFF periods (e.g. 5 mN ON: z = 5.72; 20 mN ON: z = 6.56; 75 mN ON: z = 6.56; 5 mN OFF: z = 4.03; 20 mN OFF: z = 4.58; 75 mN OFF: z = 6.02; p < 0.0001 for all comparisons).

We next asked whether this growing hyperresponsiveness reflected a shift in the intensity-response relationships of individual units. To test this, we fit the ON and OFF intensity-response relationships of each single unit with saturating exponentials to calculate a sensitivity parameter (*I*_0_, where smaller *I*_0_ indicates higher sensitivity) and considered units to be well-fitted if the sum of the residuals was < 1.2 **(Fig. S2E)**. Consistent with the increase in responsive units described above, the fraction of well-fitted RS units was consistently higher in 15- and 26-week MitoPark mice relative to age-matched controls for both ON and OFF periods (15-week ON: z = 2.9, p = 0.02; 15-week OFF: z = 3.47, p = 0.003; 26-week ON: z = 5.87, p < 0.0001; 26-week OFF: z = 5.80, p < 0.0001), while no difference in this fraction was apparent between 12-week MitoPark and control mice **(Figs. 3G-I middle panels)**. However, the *I*_0_ values of well-fitted units were indistinguishable between MitoPark and control mice at each timepoint **(Figs. 3G-I lower panels)**. There were no differences between genotypes in either of these metrics when considering the FS unit population **(Fig. S3)**. Altogether, these data suggest that MitoPark S1 hyperexcitability arises from a growing recruitment of RS units in response to the step indentations.

Concurrent elevation in baseline and evoked firing rates of RS units could enhance or degrade tactile encoding. We computed a variance-based signal-to-noise-ratio (SNR) for each RS unit during the ON and OFF periods of the stimulus to assess whether RS units show improved or compromised stimulus encoding in MitoPark mice. In layers IV and V, MitoPark RS SNR was reduced or unchanged relative to controls at 12 weeks, then increased progressively from 15 weeks to 26 weeks **(Figs. 4A-B)**. At 12 weeks, we observed a reduction in RS SNR only in layer IV during the OFF period (t = −2.59, p < 0.03, post hoc simple effect contrasts between genotypes based on the fitted LME, with Bonferroni correction for multiple comparisons), though the ON period showed a similar trend (t = −1.72, p = 0.26). At 15 weeks, SNR in both layers and periods did not differ between MitoPark and control mice (Layer IV ON: t = 1.23, p = 0.66; Layer IV OFF: t = 1.47, p = 0.43; Layer V ON: t = 0.94, p = 1.0; Layer V OFF: t = 2.01, p = 0.13.). At 26 weeks, SNR was significantly elevated in MitoPark mice relative to controls (Layer IV ON: t = 2.53, p = 0.03; Layer IV OFF: t = 2.41, p < 0.05; Layer V ON: t = 2.17, p = 0.09; Layer V OFF: t = 3.06, p = 0.006). We also analyzed the SNR of the FS population and found no significant difference between MitoPark and control mice at any timepoint **(Figs. S4A-B)**. Together, these results indicate that RS tactile-encoding fidelity in layers IV and V is enhanced at late disease stages, pointing to stage-dependent abnormal signaling in the S1 RS population.

**Figure 4.**
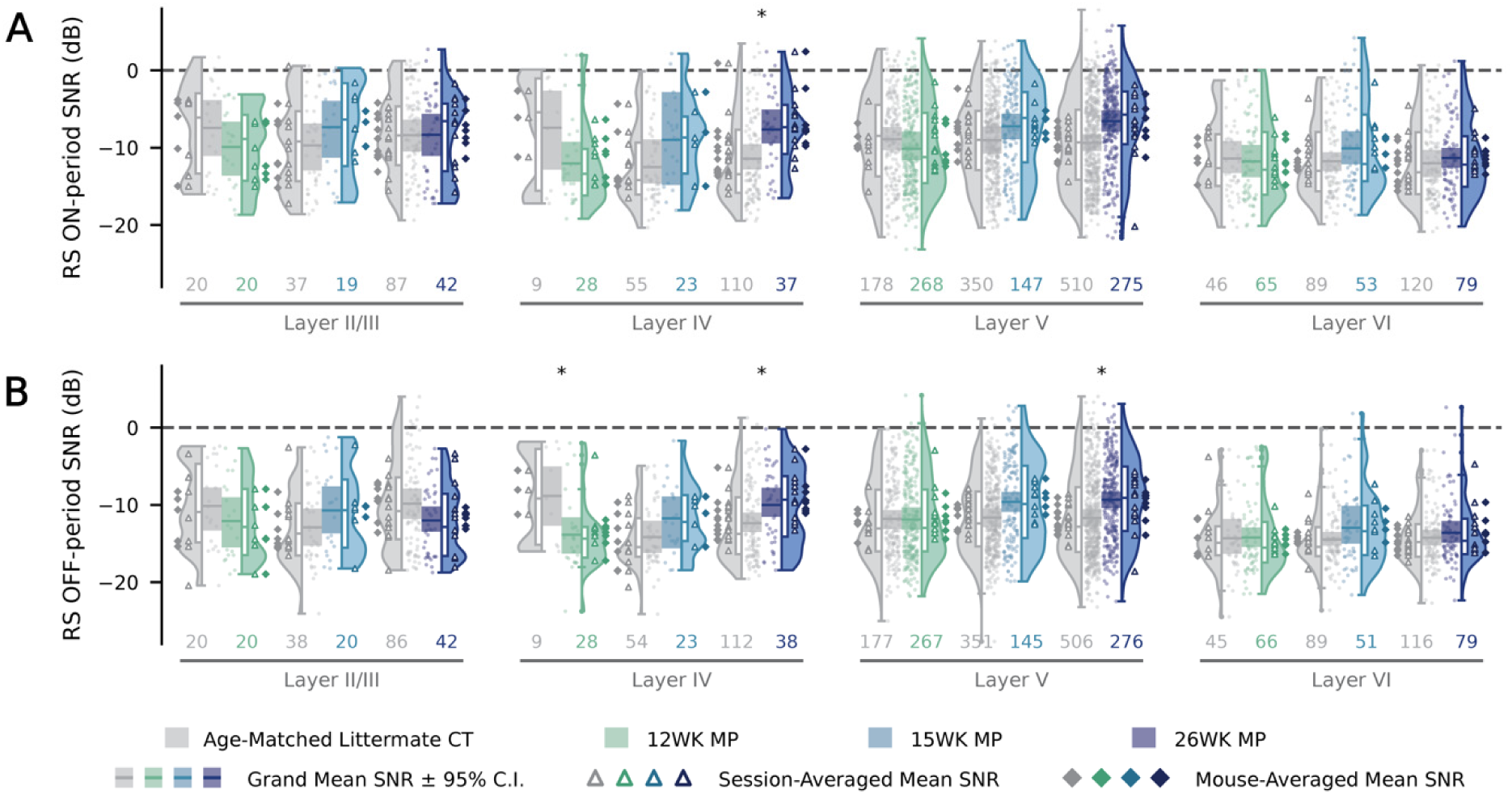
26-week MitoPark S1 Layer IV and V regular-spiking units show increased signal-to-noise ratio relative to controls. **(A)** Signal-to-noise ratio (SNR) of regular-spiking (RS) units measured for ON responses, separated by cortical layers, genotypes, and ages. Points denote firing rates for individual units, with solid horizontal bars representing the grand mean SNR in the group. Colored boxes overlaid on the horizontal bar indicate 95% C.I. of the grand mean SNR (hierarchical bootstrap). Violin plots depict the full distribution of data, with overlaid box plots (median and interquartile range) summarizing dispersion. Triangles indicate session-averaged mean SNRs, and diamonds indicate animal-averaged mean SNRs. The number of units recorded within each group is shown at the bottom. **(B)** Similar to (A), but for OFF responses. *p<0.05, post hoc simple-effects contrast test for genotype at each timepoint with Bonferroni correction for multiple comparisons. FR: firing rates; WK: week; CT: control mice; MP: MitoPark mice.

### Regular-spiking unit pairs in MitoPark S1 exhibit increased signal correlations at 26 weeks

Given the elevated excitability and responsiveness of RS units in MitoPark S1 at late stages of disease progression, we next asked whether these single-unit changes are accompanied by altered coordination of activity across the S1 population. To evaluate the stimulus-evoked tuning between simultaneously recorded S1 unit pairs, we computed signal correlation as the pairwise Pearson correlation between the concatenated, trial-averaged peristimulus firing-rate traces (20- ms bin size) of two simultaneously recorded units across all eight indentation intensities **(Fig. 5A)**. Pooling across all simultaneously recorded S1 pairs regardless of cell type, MitoPark mice showed a trend toward higher signal correlation than control mice at 26 weeks that did not reach statistical significance (t = 2.06, p = 0.119, post hoc simple effect contrasts based on the fitted LME with Bonferroni correction for multiple comparisons across ages) **(Figs. 5B-C)**. Separating the S1 pairs by RS/FS cell type, we found that this trend was driven by RS-RS pairs. Signal correlations between pairs of S1 RS units were significantly elevated in 26-week MitoPark mice relative to age-matched control mice (t = 2.82, p = 0.014), whereas no differences were detected at 12 weeks or 15 weeks **(Figs. 5D-E)**. In contrast, signal correlation did not differ between genotypes at any age for FS-FS pairs or for RS-FS pairs **(Fig. S5A-D)**.

**Figure 5.**
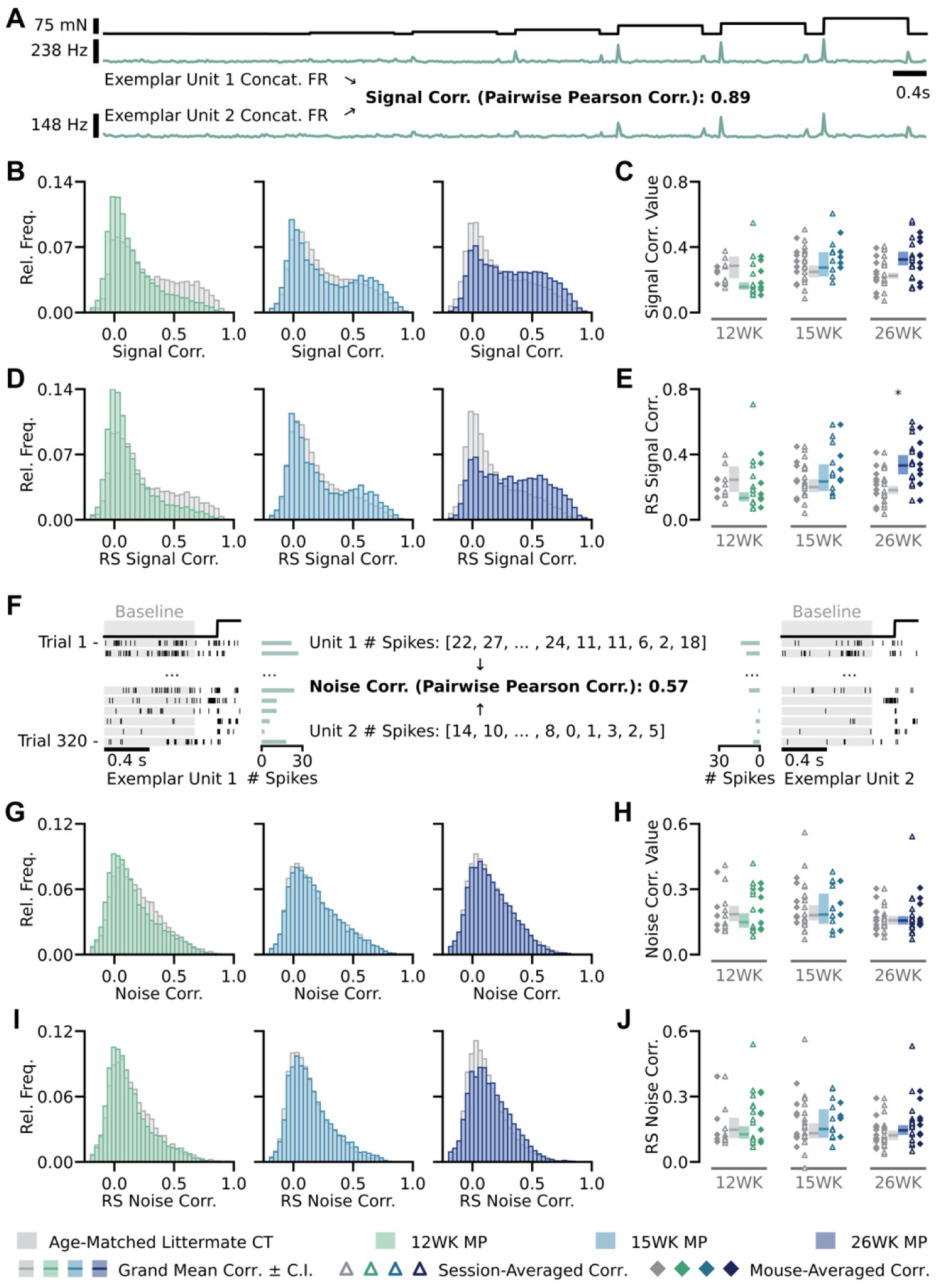
S1 Regular-spiking units of 26-week MitoPark mice exhibit increased signal correlations with no change to noise correlations. **(A)** Illustration of the signal correlation calculation using concatenated trial-averaged firing rates of an example pair of units. Stimulus (black) trace represents the concatenated evoked windows (0 to 1200 ms relative to stimulus onset) of applied force (1 to 75 mN). **(B)** Distribution of signal correlations for pairs of S1 units recorded simultaneously in control and MitoPark mice for each timepoint. **(C)** Mean (± 95% C.I.) signal correlation values for pairs of S1 units in control and MitoPark mice for each timepoint. **(D)** Similar to (B), but for distribution of signal correlations restricted to pairs of regular-spiking (RS) units. **(E)** Similar to (C), but for mean (± 95% C.I.) signal correlation values for pairs of RS units. **(F)** Illustration of the noise correlation calculation using an exemplar simultaneously recorded S1 unit pair. Correlations were calculated on trial-by-trial spike counts from an 800-ms baseline period. **(G)** Distribution of noise correlations for pairs of S1 units in control and MitoPark mice for each timepoint. **(H)** Mean (± 95% C.I.) noise correlation values for pairs of S1 units in control and MitoPark mice for each timepoint. Triangles denote session-averaged, and diamonds denote mouse-averaged noise correlation values. **(I)** Similar to (G), but for distribution of noise correlations restricted to pairs of S1 RS units in control and MitoPark mice. **(J)** Similar to (H), but for pairs of S1 RS units in control and MitoPark mice across. *p<0.05, post hoc simple-effects contrast test for genotype with Bonferroni correction for multiple comparisons. RS: regular-spiking units; FS: fast-spiking units; WK: week; CT: control mice; MP: MitoPark mice.

We next considered long-timescale noise correlations, which we computed as the pairwise Pearson correlation of trial-by-trial spike counts during the pre-stimulus baseline window (−1000 to −200 ms) for each simultaneously recorded pair of units **(Fig. 5F)**. We did not find any difference in noise correlation of S1 unit pairs between MitoPark and control mice at any timepoint, whether considered across all S1 pairs **(Figs. 5G-H)**, RS-RS pairs **(Figs. 5I-J)**, FS-FS pairs **(Figs. S5E-F)**, or RS-FS pairs **(Figs. S5G-H)**. Together, these results indicate that RS units of MitoPark mice, as expected due to their elevated firing rates (39), have enhanced tuning similarity in late stages of disease progression that likely results in amplification of signals from S1 in downstream brain regions.

### Fast-spiking S1 units in 26-week MitoPark mice are asynchronous relative to control

To examine the extent of coordinated population activity at faster timescales, we next asked whether MitoPark S1 also shows changes in millisecond-scale synchronization across the local population. We quantified population coupling (40, 41) as the correlation between the spike train of one unit and the deviance in the population firing rate of all other simultaneously recorded units from baseline (methods; **Fig. 6A)**. Considering all S1 units together, peak population coupling, which invariably occurred at a time lag of 0, did not differ significantly between MitoPark and control mice at any timepoint **(Fig. 6B)**. Populations composed of RS units alone also did not differ in population coupling between genotypes **(Fig. 6C)**. Population coupling between populations composed only of FS units of 26-week MitoPark mice, however, was significantly reduced relative to that of controls (t = −3.04, p = 0.007), though there were no differences between MitoPark and control mice at 12 or 15 weeks **(Fig. 6D)**. To test whether enhanced FS synchronization depended on the presence of the stimulus, we assessed population coupling separately for baseline and stimulus-evoked epochs. The decrease in FS coupling in 26-week MitoPark mice was present in both epochs (baseline: t = −3.07, p = 0.007; evoked: t = −2.78, p = 0.017), whereas RS coupling remained comparable between genotypes in both epochs at all ages **(Fig. S6A-D)**. These observations suggest that DA depletion in MitoPark mice selectively disrupts fine-timescale synchronization among the FS interneuron population in late stages of disease progression, without affecting synchronization among RS units.

**Figure 6.**
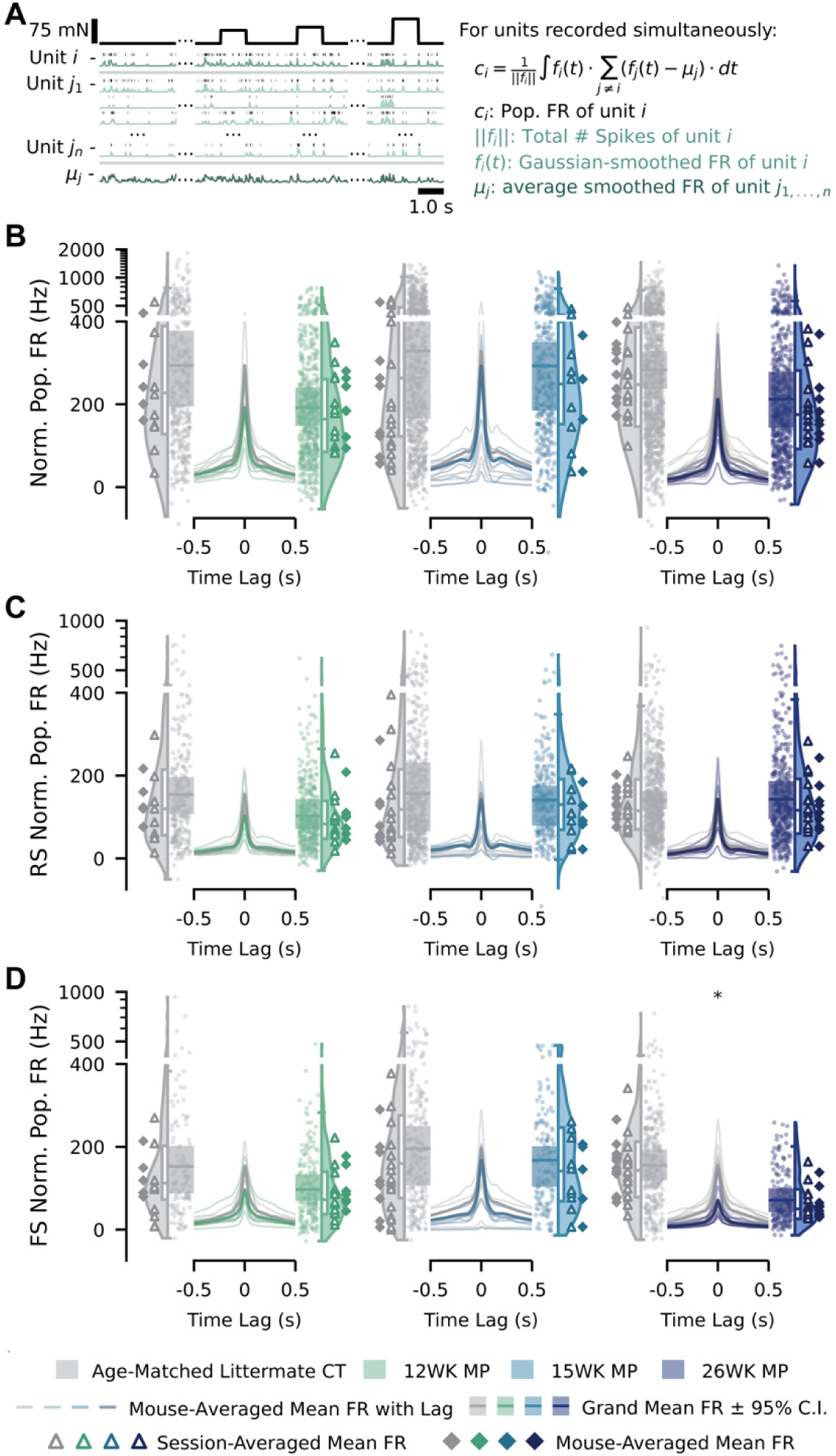
MitoPark fast-spiking populations exhibit decreased millisecond-scale synchrony at 26 weeks. **(A)** Illustration of the population coupling calculation. For each unit *i*, coupling magnitude, *c_i_*, was computed by taking the Gaussian-smoothed firing rate of unit *i*, *f_i_(t)*, and correlating it against the deviation of the average smoothed firing rate of the remaining simultaneously recorded units *j* (*μ_j_*) from baseline, integrated across a sliding time lag and normalized by the total spike count of unit *i* (||*f_i_*||). **(B)** Population coupling for all S1 units recorded in control and MitoPark mice at each timepoint. Left and right raincloud plots show the distribution of peak normalized population firing rates (FR) for control and MitoPark mice, respectively; violin plots depict the full distribution, with overlaid box plots (median and interquartile range) summarizing dispersion, and points denote individual units. The central line plot shows normalized population firing rate as a function of time lag relative to reference-unit spiking, with thin lines representing mouse-averaged mean coupling traces and the shaded band representing the grand mean ± 95% C.I. Triangles denote session-averaged and diamonds denote mouse-averaged peak coupling values. **(C)** Similar to (B), but restricted to RS units. **(D)** Similar to (B), but restricted to FS units. *p<0.05, post hoc simple-effects contrast test for genotype with Bonferroni correction for multiple comparisons. FR: firing rates; RS: regular-spiking units; FS: fast-spiking units; WK: week; CT: control mice; MP: MitoPark mice.

### Unchanged local field potential β and γ band power in MitoPark mice

Decreased population coupling amongst FS neurons may lead to overall changes within the S1 network and drive abnormal cortical gamma rhythms. Additionally, changes to S1 and primary motor cortex (M1) LFPs, such as elevated gamma power and disrupted beta power, have been reported in PD patients (42). To evaluate whether comparable changes are present in S1 of MitoPark mice, we computed power spectral densities (PSDs) of layer IV LFPs from concatenated baseline and stimulus-evoked epochs for each recording session **(Fig. S7A)**.

Because overall power varied considerably across sessions independent of genotype or time point, we normalized each PSD by dividing the area under the curve (AUC) of the baseline PSD before comparing β (10 – 30 Hz) and γ (30 – 100 Hz) band power between MitoPark and control mice. This baseline AUC did not differ by genotype at any age (post hoc simple effect contrasts based on the fitted LME with Bonferroni correction for multiple comparisons) **(Fig. S7B)**.

Normalized β and γ band power during the baseline period were likewise comparable between MitoPark and control mice at all time points (β: all p ≥ 0.086; γ: all p ≥ 0.92) **(Figs. S7C, E)**. As expected, evoked (40 mN) β and γ power was 2- to 5-fold higher than baseline power in both genotypes, but this stimulus-driven increase did not differ between MitoPark and control mice at any timepoint (β: all p ≥ 0.091; γ: all p ≥ 0.69) **(Figs. S7D, F)**. Thus, despite cell-type specific changes to pairwise interactions and population metrics, there was no overt alteration to β or γ oscillatory power in MitoPark mice.

## Discussion

In this study, we found that loss of TH^+^ axons in S1 of MitoPark mice parallels the well-established degeneration of midbrain DA neurons and striatal TH^+^ innervation (25, 43). We observed multiple functional abnormalities that coincided with this anatomical change, including hyperexcitable RS S1 units in layer V of 26-week MitoPark mice, the principal output pathway of the canonical cortical circuit (44). Tactile stimuli recruited a larger fraction of RS units in advanced-age MitoPark S1. This RS-specific hyperexcitability was also associated with increased signal correlations between pairs of RS units. By contrast, the excitability of MitoPark FS units appeared comparable to controls at all ages and the only phenotype observed within FS units was decreased population coupling, reflecting asynchronous recruitment of this cell type. Overall, the progressive loss of DA neurons in MitoPark mice was associated with the emergence of a hyperexcitable S1.

One central question raised by our results is what drives hyperexcitability and hyperresponsiveness specifically in RS units of advanced-age MitoPark S1. Since population coupling reflects how synchronized a unit’s activity is with the rest of the population (40), the FS desynchronization in 26-week MitoPark S1 indicates that FS neurons, despite maintaining a comparable excitability with controls, may provide less temporally coordinated and, therefore, less effective inhibition to S1 RS neurons (45). Reduced inhibitory efficacy offers a plausible circuit-level mechanism for RS-specific hyperexcitability in advanced-age MitoPark S1.

The loss of DA fiber innervation of S1 may directly contribute to this phenotype. DA axons innervating healthy S1 may modulate cortical processing of tactile signals via D1-like (D1, D5) and D2-like (D2, D3, D4) receptors (46–48), which activate and inhibit cAMP signaling respectively (49, 50). In the cerebral cortex, D1 receptors are the most abundant followed by D2 (49). Spatial transcriptomics data indicate that D1 and D2 receptors are expressed in minimally overlapping S1 cell populations, with D1 receptor expression notably enriched in parvalbumin-expressing interneurons (51). This raises the possibility that DA acts directly on FS neurons to regulate their synchrony, such that absence of DA in advanced-stage MitoPark S1 destabilizes FS population coupling independent of any change in FS excitability contributing to S1 hyperexcitability. Such a local role for DA signaling is additionally supported by the observation that acute pharmacological blockade of D1 or D2 receptors in the rat sensorimotor cortex increased somatosensory evoked potential amplitudes (52), and infusion of DA reversibly inhibits layer V pyramidal cells in rat medial prefrontal cortex (53). However, since MitoPark mice experience systemic degeneration of DAT-expressing cells, pathways beyond local S1 DA innervation may also shape the phenotype and regulate sensory processing. The prefrontal cortex and dorsolateral striatum receive dense DA innervation and are connected to S1 through corticocortical and corticostriatal-thalamocortical circuits, respectively (54, 55). Diminished mesocortical and nigrostriatal pathways may indirectly modulate S1 activity through these pathways, independent of or in combination with local DA within S1. Notably, lesions of the medial forebrain bundle with 6-OHDA produce pain-like behavioral phenotypes alongside altered bursting in wide-dynamic-range neurons of the spinal cord dorsal horn (56, 57), raising the possibility that descending DAnergic or DA-dependent serotonergic spinal projections contribute to hyperexcitability of S1 in MitoPark mice (15). Disentangling the relative contributions of local S1 DA innervation and other DA pathways to S1 hyperexcitability will require pathway-specific manipulations in future work.

Another outstanding question is whether hyperexcitable S1 in MitoPark mice relates to the tactile aberration experienced by PD patients. Our observations provide a potential explanation for both hyperalgesia and decreased tactile acuity in PD patients. S1 is involved in the perception of both innocuous and noxious somatosensory signals (58–60). Therefore, elevated excitability of pyramidal neurons, especially in layer V, could explain PD-related neuropathic pain or hyperalgesia (61, 62). Meanwhile, the expanded recruitment of pyramidal neurons by tactile stimuli likely reflects increased receptive field sizes (35). If so, the spatial discriminative capacity of the S1 population is likely reduced in MitoPark mice.

Other PD-related pathologies, such as the accumulation of α-synuclein or peripheral neuropathy, may also contribute to tactile aberration (14–18). Given the independence of α-synuclein pathology in MitoPark mice (25), our results indicate the loss of DA alone is sufficient for S1 hyperexcitability. However, this does not exclude other contributors that interact with DA loss in human patients to ultimately shape sensory phenotypes. For instance, α-synuclein accumulation in central pain processing regions, including the spinal cord dorsal horn, can result in an augmented nociceptive responses (63), while peripheral neuropathy or malformed peripheral end organs of sensory neurons may also contribute to tactile dysfunction and pain (14, 64). Determining how these factors combine, and whether their relative contributions vary across patients, will be important both for interpreting heterogeneity in PD sensory symptoms and for identifying which patients might benefit most from DA-targeted versus circuit-targeted interventions.

Notably, we did not detect changes in β or γ band power in S1 LFPs of MitoPark mice at any age, in contrast to the altered β and γ dynamics reported in S1 and M1 of human PD patients (42). This dissociation suggests that alteration in PD patiently likely arise from a non-DA-dependent factors or a combination of pathologies that are not fully recapitulated by the MitoPark model.

Our results provide a systematic characterization of single-unit activity and population dynamics of MitoPark S1 across disease progression. Looking forward, dissecting other non-DA-dependent pathologies in PD-related somatosensory deficits and the circuits that caused the hyperexcitability in advanced-age MitoPark S1 will be necessary to build a complete mechanistic picture of somatosensory dysfunction in PD. Understanding the mechanism of PD-related pain and tactile aberrations should translate to better therapeutics for improving the sensorimotor functions and development of non-invasive neuromodulation of the somatosensory system.

## Data, Materials, and Software Availability

Data and analysis code are available upon reasonable request to the corresponding author.

## Acknowledgements

We thank Katherine Greenhut and Yuna Lee for assistance with genotyping and mouse husbandry; Christopher Rodgers and Dieter Jaeger for providing the MitoPark breeders; and Christopher Rodgers and members of the Emanuel lab for discussions.

## Author Contributions

R.Z., R.Q., and A.J.E. conceptualized the project. R.Z. and R.Q. collected and analyzed data with guidance from A.J.E. R.Z., R.Q., and A.J.E. wrote the manuscript, contributed to editing and approved the final version.

## Materials and Methods

### Animals

Animal husbandry and experimental procedures were performed in accordance with protocol PROTO202200102 approved by the Emory University School of Medicine Institutional Animal Care and Use Committee (IACUC). Mice were group-housed under a 12-hour light-dark cycle with *ad libitum* access to standard food and water in a temperature- and humidity-controlled rodent facility in Emory University’s Whitehead Biomedical Research Building. All animals had access to running wheels within their cages.

MitoPark mice (25) were generated by crossing *DAT^IREScre^* mice (RRID:IMSR_JAX:006660), which express Cre from the *Slc6a3* locus (65), with *Tfam^loxP^* mice (RRID:IMSR_JAX:026123), in which *loxP* sites flank critical exons of the mitochondrial transcription factor A (*Tfam*) gene (66). The resulting *DAT^Cre/+^;Tfam^fl/+^* male mice were crossed with *Tfam^fl/fl^* female mice to produce MitoPark (*DAT^Cre/+^;Tfam^fl/fl^*) and control (*+/+;Tfam^fl/fl^*, *+/+;Tfam^fl/+^*, and *DAT^Cre/+^;Tfam^fl/+^*) mice. Initial breeders were generously provided by Drs. Christopher Rodgers and Dieter Jaeger. All animals were maintained on the C57BL/6J background. Consistent with previous reports (27), 26-week-old MitoPark mice moved noticeably less in their home cages than controls and differences were not readily apparent at 12 or 15 weeks. Experimenter blinding to genotype was maintained through data acquisition and preprocessing.

### Immunohistochemistry

We deeply anesthetized mice with >3% inhaled isoflurane (Piramal Critical Care, Inc., Bethlehem, PA, USA) and transcardially perfused them with 15 mL of iced phosphate-buffered saline (PBS), followed by 15 mL of iced 4% paraformaldehyde (PFA) (Electron Microscopy Science, Hatfield, PA, USA) in PBS. We extracted brains and post-fixed them in 4% PFA solution for 1 hr. After fixation, we rinsed brains three times in PBS for 15 min each and cryoprotected them in 30% sucrose in PBS at 4 °C for at least 24 hrs. We then embedded brains in O.C.T. compound (Fisher HealthCare, Houston, TX, USA) and froze them at −80 °C. We sliced brains into 50-μm sections using a cryostat (Leica, Wetzlar, Germany) and directly mounted them onto glass slides (Superfrost Plus; Thermo Fisher Scientific, Waltham, MA, USA). Brain sections were first dried overnight in the dark at room temperature and rehydrated in PBS before staining.

We immunostained sections for tyrosine hydroxylase (TH) by washing twice for 5 min each with 1× PBS containing 0.1% Triton X-100 (0.1% PBST), then incubating with blocking solution [5% normal donkey serum (017-000-121; Jackson ImmunoResearch Labs, West Grove, PA, USA) in 0.1% PBST] for 2 hrs at room temperature and then incubating with primary antibodies (rabbit anti-TH; AB152; Millipore Sigma, Burlington, MA, USA; RRID: AB_390204; dilution 1:500) in blocking solutions at 4°C for 24 hrs. We then washed the tissue three times (10 min) with 0.1% PBST and subsequently incubated with secondary antibodies (donkey anti-rabbit conjugated to Alexa 546; A10040; ThermoFisher, Waltham, MA, USA; RRID: AB_2534016; dilution: 1:750) in blocking solution for 2 hrs at room temperature. After incubation, we washed three times (10 mins) with 0.1% PBST and applied an additional wash (5 mins) with PBS including 4′,6-diamidino-2-phenylindole (DAPI) (5 µg mL^−1^; AAT Bioquest, Pleasanton, CA, USA). Finally, we washed with PBS for 5 min and mounted with Fluoromount-G (Southern Biotech, Birmingham, AL, USA). We imaged tissue with a BZX-7000 automated fluorescence microscope (Keyence, Itasca, IL, USA). To quantify the intensity of TH staining of dlSTR and midbrain in MitoPark mice and control mice, we imaged tissues with fixed exposure time across all image conditions and calculated the mean pixel intensity within the region of interest using Fiji (67), which we normalized by dividing by the maximum value of TH intensity across ages and genotypes. We traced TH^+^ fibers in one section of forelimb S1 per animal using the SNT v4.2.1 plugin in FIJI (68). Regions of interest (ROIs) for each layer were drawn using the DAPI image. For each layer, we skeletonized the tracing paths within the ROI and used the analyze skeleton tool to compute the total path length in mm. Fiber density was calculated by dividing this length by the total volume (in mm^3^; taking into account the 0.05-mm section thickness) of the ROI.

### Surgery & Craniotomy

All experimental animals received an intraperitoneal (i.p.) injection of dexamethasone (2.0 mg kg^-1^; NDC63323-165-30, Fresenius Kabi, Lake Zurich, IL, USA) 4-12 hrs before surgery to minimize postoperative swelling. We used dual-mode analgesia to minimize pain and discomfort associated with surgery. Immediately before the procedure, we administered sustained-release meloxicam (3.0 mg kg^−1^ s.c.; NDC79926-070-17, Wedgewood Pharmacy, Swedesboro, NJ, USA). During surgery, mice were anaesthetized with isoflurane (2.5% induction; 1.5% maintenance) and locally at the incision with lidocaine (6.0 mg kg^-1^; NDC63323-201-03, Fresenius Kabi, Lake Zurich, IL, USA). After fur removal and sterile skin preparation, we incised the scalp and periosteum, dried the skull, and mounted a titanium headplate to the skull with dental acrylic (Metabond, Parkell, Edgewood, NY, USA). We made a circular craniotomy (∼1 mm in diameter) above forepaw S1 (right hemisphere: 2.10 mm lateral, 0.00 mm posterior to bregma). We sealed the craniotomy with Kwik-Sil (WPI, Sarasota, FL, USA) and cemented a 3D-printed resin ring onto the headplate to provide a well for a recording bath solution. Extended-release buprenorphine (1.0 mg kg^−1^ s.c.; Wedgewood Pharmacy, Swedesboro, NJ, USA) was administered post-operatively for analgesia. We closely monitored the post-operative condition of the mice until awake and allowed mice to recover overnight before the first recording session.

### In vivo Cortex Electrophysiology

All recordings were performed in forepaw S1 (right hemisphere: 2.05 to 2.35 mm lateral, 0.25 anterior to 0.15 mm posterior to bregma) of awake, head-fixed mice. Prior to the first recording, mice underwent at least three handling sessions (5-10 mins each) and were habituated to head restraint (10-15 mins). Then, we removed the Kwik-Sil sealant and submerged the craniotomy in a HEPES-buffered saline solution (150 mM NaCl, 2.5 mM KCl and 10 mM HEPES, 5mM NaOH; pH 7.4). We inserted a Neuropixels 1.0 probe into the forepaw S1 of the right hemisphere and advanced the probe tip to ∼1600 µm below the dura. To verify S1 placement, we gently brushed the glabrous skin on the plantar side of the contralateral (left) forepaw with a fine paintbrush. We monitored brush-evoked spikes across multiple cortical recording channels that span the depth of the cortex to approximate the receptive field location. If the receptive field was not on the forepaw pedal pads, we removed the probe from the brain and reinserted it into a new location. A maximum of four insertions were allowed per mouse per day. In rare cases, mice requiring more than four attempts to locate the target area were excluded from that day’s experiments to allow for recovery. Once we were satisfied with the insertion location, we restrained the left forepaw with vinyl tape in a natural position over a rectangular aperture (approx. 7 × 4 mm in size) cut into an acrylic platform supporting the mouse.

During each recording session, we used a cylindrical indenting probe (0.5 mm in diameter; Teflon-tipped) controlled by a dual-mode force controller (300C-I; Aurora Scientific, Aurora, ON, Canada) to apply stimuli to the paw through the aperture. The indenter was aligned beneath the pad that exhibited the strongest response during manual receptive field mapping and a minimal force was applied to keep the indenter tip in contact with the skin. This holding force is not expected to evoke a response from most sensory neurons (35). The position, force, and displacement of the indenter were commanded via custom Python scripts interfacing with NIDAQ board (PXIe-6341; National Instruments, Austin, TX, USA), connected through an BNC-2090A breakout box (National Instruments, Austin, TX, USA). After the placement of the indenter, the bath solution was replaced with HEPES-buffered saline solution with 1% agarose to stabilize the shank. Recording sessions were restricted to a maximum duration of 45 minutes.

We acquired data using the Neuropixels PXIe-1071 control system (IMEC, Leuven, Belgium) and Open Ephys GUI software (69). The ground electrode was attached via an alligator clip to the edge of the headplate well, with its tip submerged in the bathing solution. Raw signals were collected, digitized (30 kHz), amplified (500×), bandpass filtered (0.5-10,000 Hz), and saved.

### Spike Sorting

We concatenated all binary files from the same recording session for spike sorting with Kilosort 4 (70). The resulting spike clusters were manually curated using Phy2 (https://github.com/cortex-lab/phy) and classified as single units, multi-units, or noise. Only clusters classified as single units were included in subsequent analysis. The following criteria were used to classify single units: (1) a clear inter-spike interval (> 2 ms) evident in the auto-correlogram; (2) waveforms of action potentials had large spike amplitudes relative to baseline noise; (3) waveforms were distinguishable from nearby clusters; and (4) the overall firing rate was > 0.15 Hz.

### Laminar and Cell-Type Identification

We determined the laminar locations of individual S1 units and LFP channels based on the spatial distribution of waveforms along the probe, which we registered to physiological indicators of cortical layers, as described and validated previously (35, 71). Briefly, we calculated the depth of each channel using the depth of shank tip recorded directly from the probe manipulator. To ensure accurate laminar localization, depth values were corrected for the probe’s insertion angle and further refined by identifying the center of layer IV through current source density (CSD) analysis. To do so, we low-pass filtered voltage waveforms from the LFP band (0.5–500 Hz) at 250 Hz with a third-order Butterworth filter and then calculated the second derivative of this signal across laminar locations on the probe. We rigidly adjusted the depth of S1 units and LFP channels so that the center of layer IV identified in the CSD plot was assigned to 476 µm below the cortical surface. S1 units and LFP channels were classified into cortical layers according to the following layer depths: layer II/III: 119–416.5 µm; layer IV: 416.5–535.5 µm; layer V: 535.5–952 µm; and layer VI: 952–1200 µm. We classified S1 units as regular-spiking (RS) or fast-spiking (FS) based on the trough-to-peak times of the spike waveform on the channel with the largest amplitude. We defined FS units as those with trough-to-peak times ≤0.5 ms and RS units as those with trough-to-peak times >0.5 ms.

### Extraction of Baseline and Maximum-Evoked Firing Rates

During each recording session, force-controlled indentations were delivered at 8 different intensities (1, 2, 5, 10, 20, 40, 50, and 75 mN) above the minimal force required to keep the indenter tip in contact with the skin, with 40 repetitions per force level. Each trial lasted 3 seconds and consisted of a 1-second pre-stimulus baseline, a 1-s sustained force step of randomized intensity, and a 1-second post-stimulus period. Stimuli were filtered with a 30-ms boxcar filter, generating a 30-ms ramp from baseline to full intensity at the onset and offset of the stimulus presentation. Firing rate of each unit (in Hz) was calculated by counting spikes within non-overlapping 20 ms time bins (150 bins in total for 3-sec trials). The baseline firing rate was defined as the mean firing rate in a 0.8-s pre-stimulus window (−1000 to −200 ms), averaged across all trials. The maximum evoked firing rate was defined as the maximum value of baseline-subtracted and trial-averaged firing rate across all time bins and stimulus intensities.

### Analysis of Intensity-Response Relationships

Response magnitude and unit sensitivity were quantified to assess intensity-response relationships. Firstly, the fraction of units responsive to different phases of the step indentation— ON (0–40 ms after stimulus onset), OFF (0–40 ms after stimulus offset—was calculated separately. A unit was classified as responsive if it met the following criteria: (1) its mean z-scored firing rate within the designated response window exceeded the 95% C.I. of its baseline firing rate; (2) it had a monotonic intensity-response relationship across force steps with different intensities. Secondly, the intensity-response relationship during the onset and offset period of each unit was fit with a saturating exponential, 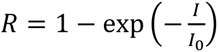, where *R* is the mean firing rates at each intensity, *I* is the intensity of force steps, and *I*_0_ is the fitted parameter indicating the sensitivity threshold of this single unit. We calculated the fraction of well-fitted units (∑|residuals| < 1.2) and analyzed the distribution of *I*_0_ for all well-fitted units.

### Computation of Signal-to-Noise Ratio

A variance-based SNR was computed for each cortical unit to quantify how reliably a unit’s firing rate distinguishes stimulus intensities relative to its trial-to-trial variability. ON and OFF responses were analyzed independently. The mean baseline-subtracted firing rate was computed per trial by averaging across the two 20-ms PSTH bins spanning the window. SNR was then defined as:

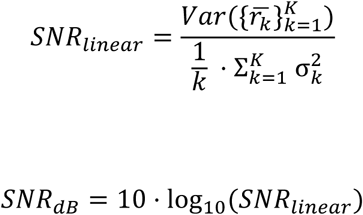

where *r̅_k_* is the trial-averaged delta firing rate at *k* intensity (*K*= 8 intensities: 1, 2, 5, 10, 20, 40, 50, and 75 mN), and 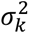 is the within-intensity trial variance. The numerator captures how much the mean response varies across stimulus intensities (signal), and the denominator is the average trial-to-trial variability pooled across intensities (noise). We excluded units with zero variance or undefined signal variance (i.e. not responsive to stimuli). We converted linear SNR to decibels to facilitate comparison across layers and cell types.

### Noise Correlation and Signal Correlation

We computed signal and noise correlation between all pairs of cortical units that were recorded simultaneously and visualized the correlation values for interactions between different cell types: between RS unit pairs only, between FS unit pairs only, and between RS-FS unit pairs. **Noise correlation:** For each unit, spikes occurring within −1000 to −200 ms pre-stimulus window (time aligned to the onset of stimulus) were counted per trial to form a sweep-by-sweep spike-count vector. Pairwise Pearson correlations were computed between all unit pairs within each recording, yielding a symmetric n × n matrix with units ordered by recording depth. Unique pair values were extracted from the strict lower triangle of each matrix. **Signal correlation:** PSTHs were constructed by binning spike times into 20-ms bins and converting bin counts to firing rate (Hz). For each unit, the 40 trials per stimulus were averaged to obtain a trial-averaged PSTH, then the stimulus-driven response was isolated by extracting the bins spanning 0 to 1200 ms (time aligned to the onset of stimulus) for each of the 8 stimulus intensities. Pairwise Pearson correlations between signal vectors were computed for all unit pairs within each recording.

### Population Coupling

To assess population coupling (40, 41), the normalized population firing rate of each unit (*c_i_*) was estimated by the formula 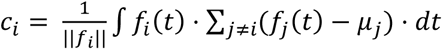, where ||*f_i_*|| is the total number of spikes generated by unit *i*, *f_i_* represents its Gaussian-smoothed firing rate (binsize: 1 ms, half width of 12/√2 ms), and *μ*_j_ is the mean firing rate of unit *j* across the entire recording. The integral was evaluated over the full recording duration, yielding a scalar coupling index for each unit that reflects how strongly its firing covaries with the summed activity of all simultaneously recorded neurons. To further characterize the time course of population coupling, the cross-correlation between *f_i_* and the summed population activity ∑*_j_*_≠_*_i_*(*f*_j_(*t*) − *μ*_j_) was computed across time lags spanning ±500 ms; the value at lag = 0 served as the coupling index for statistical comparisons. Analyses were performed separately for all cortical units combined, within RS units only, and within FS units only. Additionally, population coupling was computed for RS and FS units during baseline (−1000 to −200 ms) and evoked (0 to 1200 ms) windows separately.

### LFP Power

For each recording session, LFP signals from Layer IV channels, identified by current source density (CSD) analysis, were used for spectral analysis. The LFP was low-pass filtered at 250 Hz with a zero-phase, 8th-order Butterworth filter. Baseline epochs (−1000 to −200 ms relative to stimulus onset) were concatenated across all trials, regardless of stimulus intensity, while evoked epochs (0 to 1200 ms relative to stimulus onset) were concatenated across trials of the 40 mN intensity only. Power spectral densities (PSDs) were computed for each concatenated LFP across the 1–100 Hz frequency range. We normalized PSDs by dividing them the area under the curve (AUC) of their respective Layer IV baseline PSD, integrated across the full 1–100 Hz range. Normalized β (10–30 Hz) and γ (30–100 Hz) band power was then calculated as the integral of the normalized PSD within each respective frequency band, computed separately for baseline and evoked (40 mN) epochs.

