## Supplemental Information for "Hyperexcitable primary somatosensory cortex in the MitoPark mouse model of Parkinson’s disease"

**This PDF file includes:**

Figures S1 to S7

Table S1

Statistical Tables Appendix

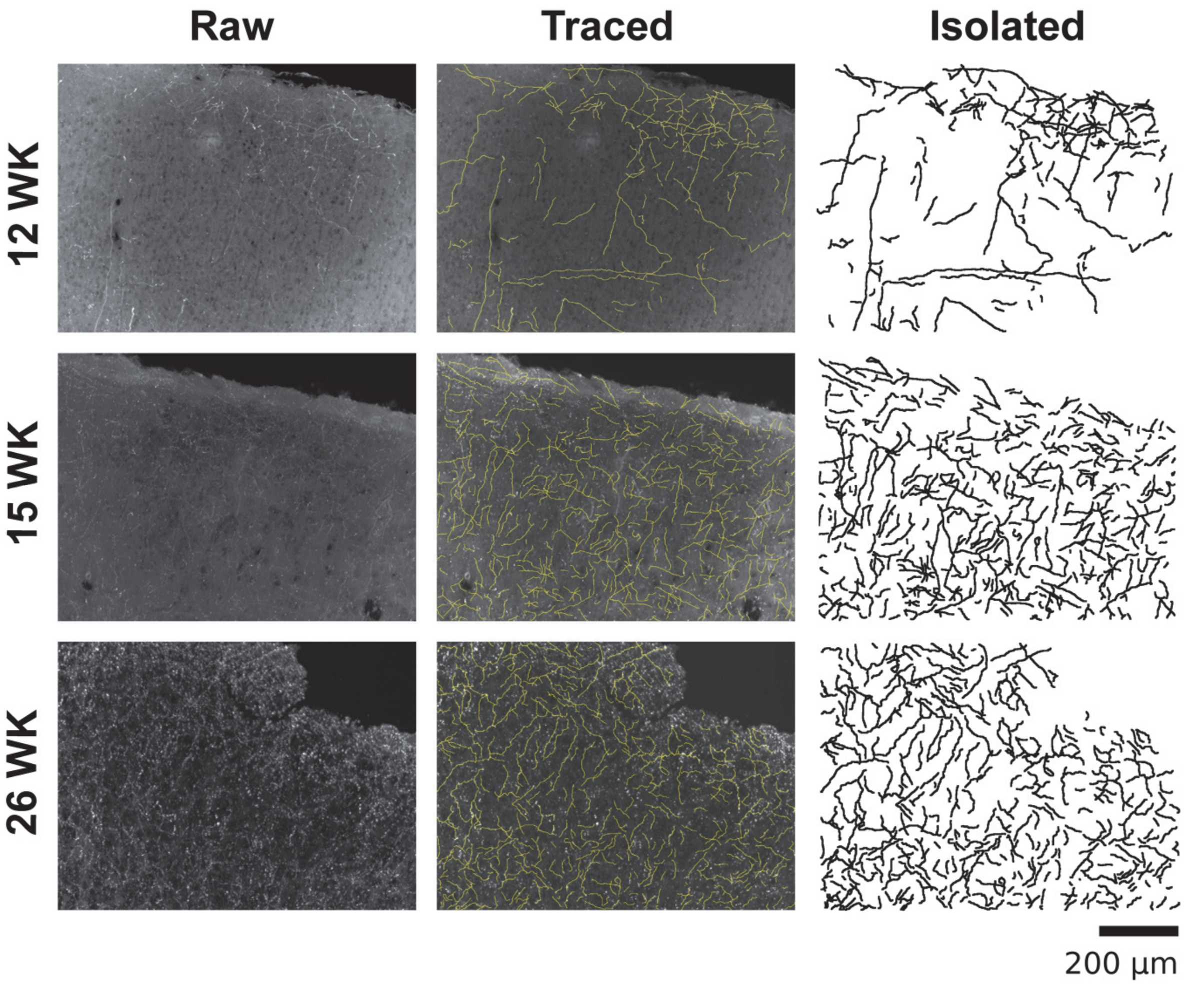


**Figure S1. Representative TH immunofluorescence and fiber tracing in superficial S1 of control mice. Left:** Raw TH immunofluorescence images from three control mice, one per time point. **Middle:** Overlay showing colocalization of immunofluorescence signal with traced fibers. **Right:** Isolated $TH^{+}$ fibers. WK: weeks.

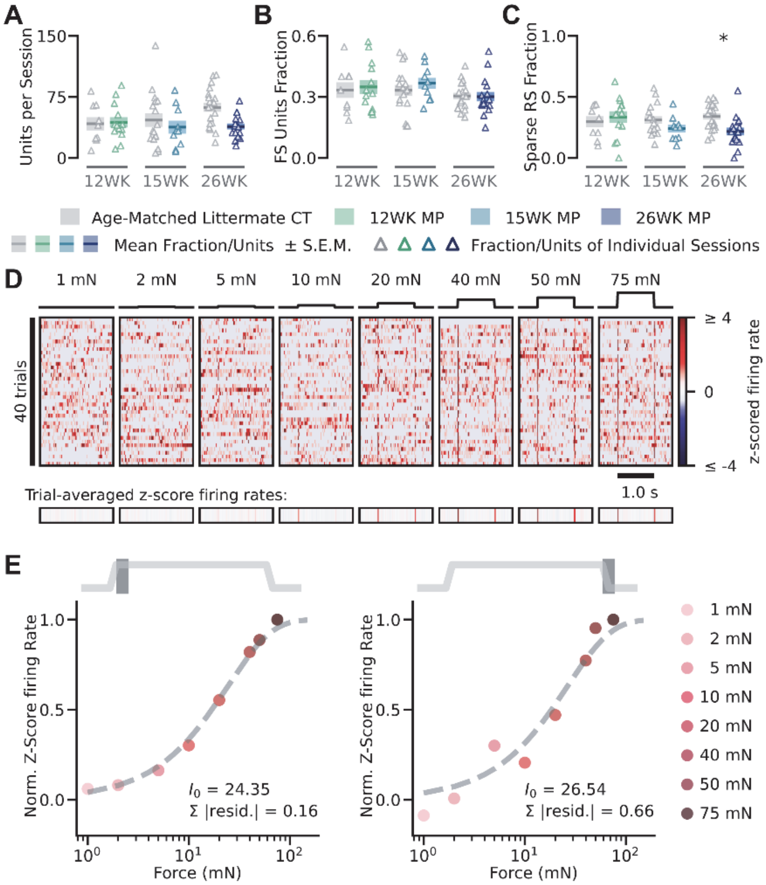


**Figure S2. Calculation of mean z-scored firing rates and the sensitivity parameter, $I_0$. (A)** Single-unit yield for each session. Hollow triangles represent number of units obtained from each individual recording session; horizontal bar is the mean value across all recordings; and the colored box indicates the S.E.M. of the mean value. **(B)** Fraction of fast-spiking (FS) units recorded per recording session. **(C)** Fraction of sparse-firing regular-spiking units (baseline firing rates < 1 Hz) for each session. **(D) Top:** Z-scored firing rate for each individual trial of the examplar unit shown in Fig. 2D. Order of stimulus presentation was randomized. Here, the trials are grouped by stimulus intensity (40 trials per intensity) and sorted according to chronological order of presentation. **Bottom:** Trial-averaged z-scored firing rates in response to each intensity. **(E)** Normalized trial-averaged z-scored firing rates of the ON response (left) and the OFF response (right) from the same unit and the best fit to a saturating exponential function (gray dashed line). $I_0$: fitted single-unit sensitivity parameter; Σ |resid.|: the sum of the absolute values of the fit residuals; WK: week; CT: control mice; MP: MitoPark mice.

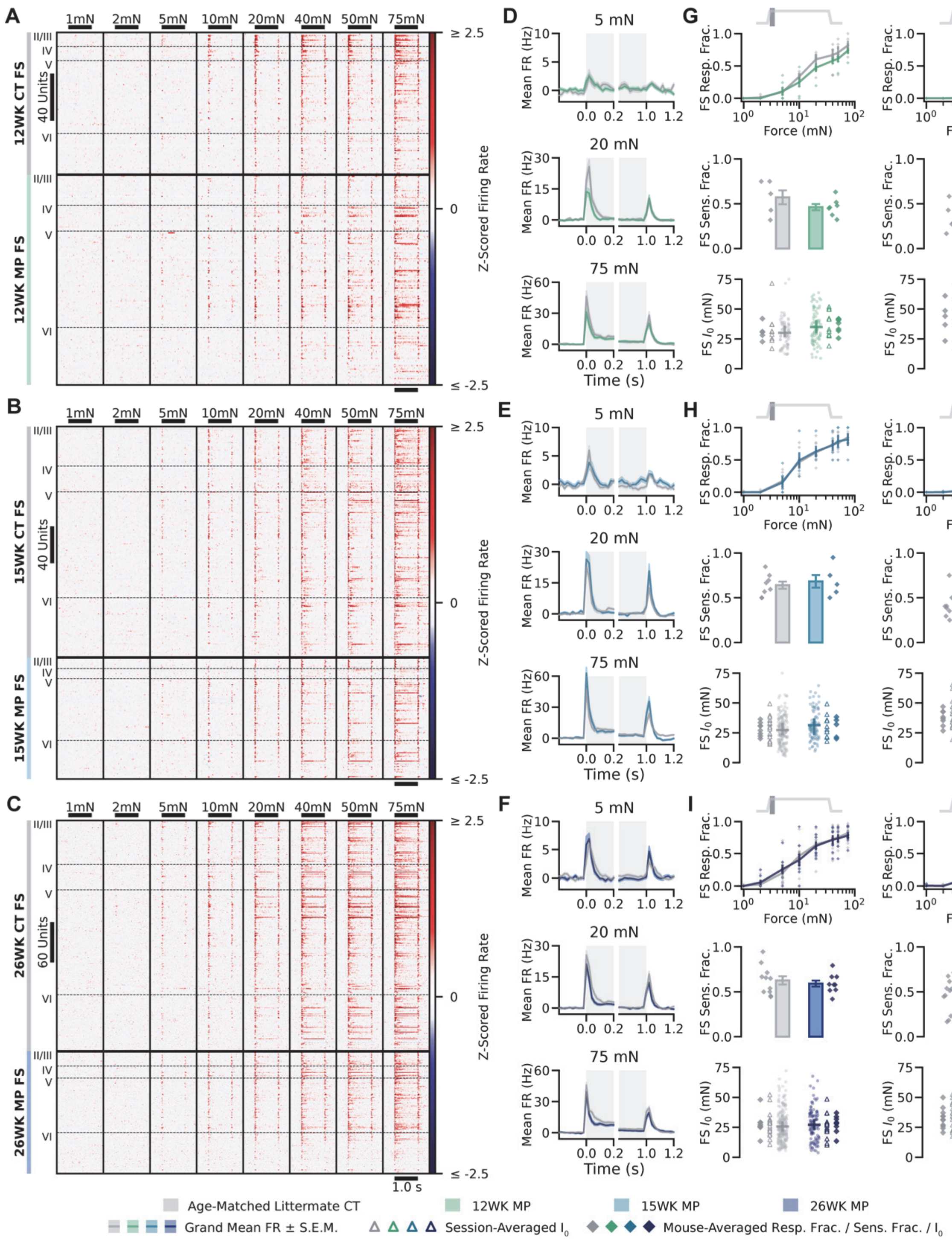


34

35 **Figure S3. Responsiveness and sensitivity of MP S1 fast-spiking units are similar to**
36 **controls at all ages. (A-I)** Similar to Fig. 3A-I, but for fast-spiking (FS) units.

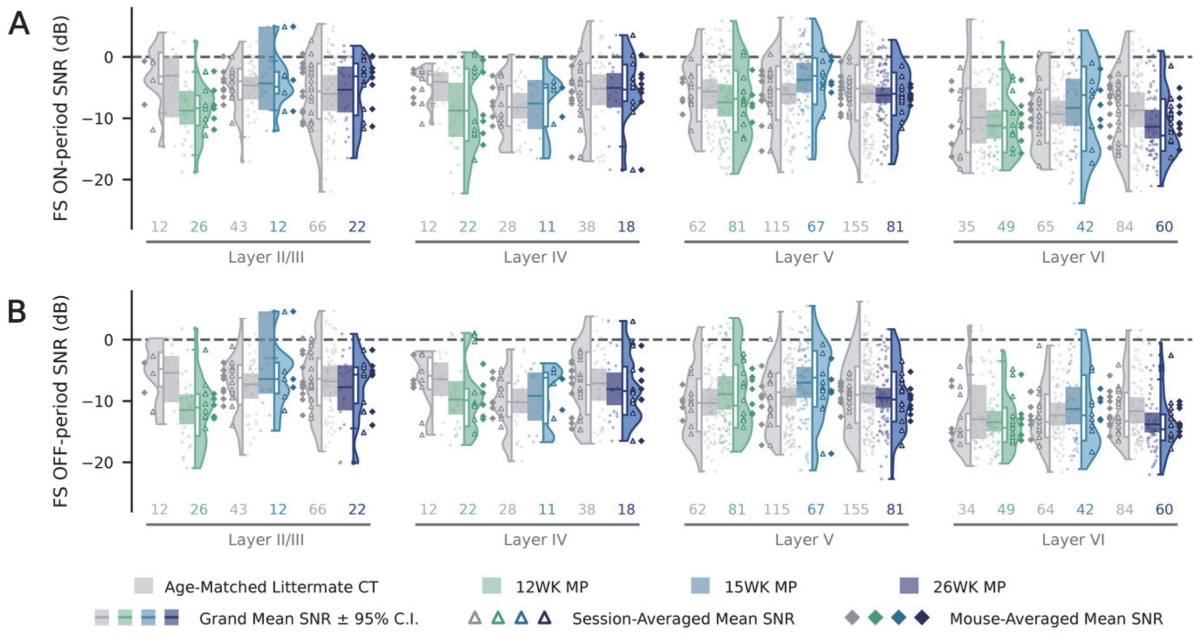


**Figure S4. No difference in SNR between S1 FS units of MitoPark and control mice. (A)** SNR of fast-spiking (FS) units for ON responses, separated by cortical layers, genotypes, and ages. Points denote firing rates for individual units, with solid horizontal bars representing the grand mean SNR in the group. Colored boxes overlaid on the horizontal bar indicate 95% C.I. of the grand mean SNR (hierarchical bootstrap). Violin plots depict the full distribution of data, with overlaid box plots (median and interquartile range) summarizing dispersion. Triangles indicate session-averaged mean SNRs and diamonds indicate animal-averaged mean SNRs. The number of units recorded within each group is shown at the bottom. **(B)** Similar to (A), but for SNRs of FS units for OFF responses. *$p<0.05$, post hoc simple-effects contrast test for genotype at each sampled age with Bonferroni correction for multiple comparisons. FR: firing rates; WK: week; CT: control mice; MP: MitoPark mice.

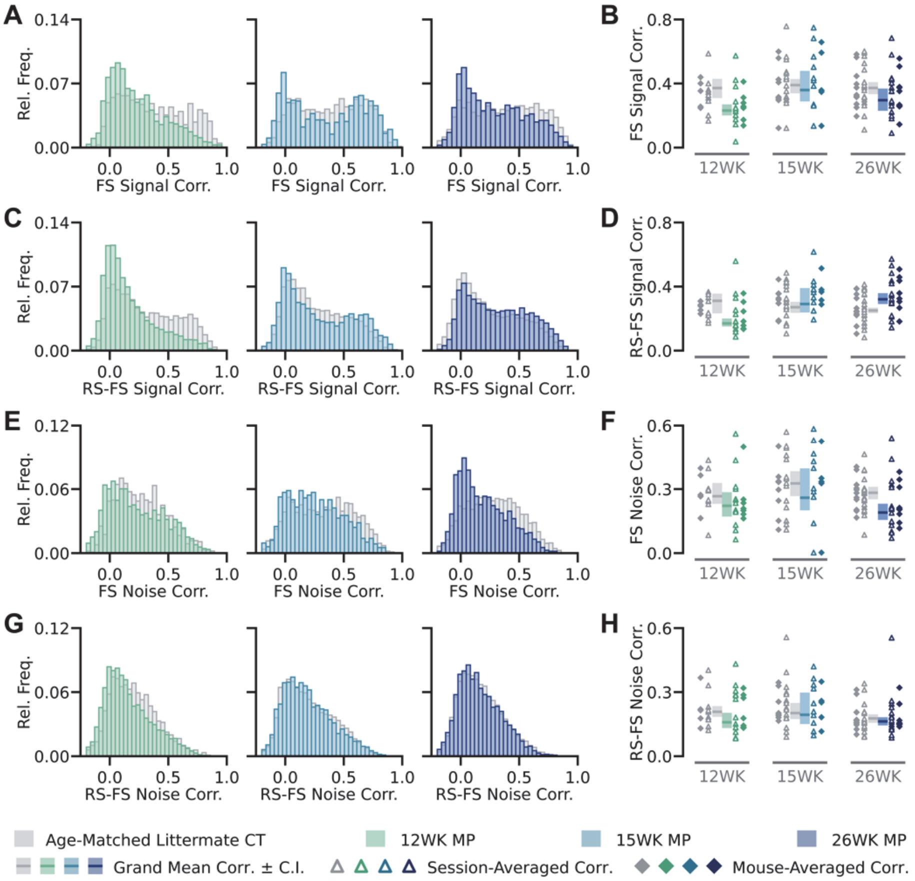


**Figure S5. No difference in noise or signal correlation between control and MitoPark S1 fast-spiking (FS) or regular-spiking (RS)-FS unit pairs. (A)** Distribution of signal correlations for pairs of S1 FS units in control and MitoPark mice for each timepoint. **(B)** Mean (± 95% C.I.) signal correlation values for pairs of S1 FS units in control and MitoPark mice for each timepoint. Triangles denote session-averaged, and diamonds denote mouse-averaged noise correlation values. **(C)** Distribution of signal correlations for RS-FS unit pairs in control and MitoPark mice for each timepoint. **(D)** Mean (± 95% C.I.) signal correlation values for RS-FS unit pairs in control and MitoPark mice for each timepoint. **(E)** Distribution of noise correlations for pairs of S1 FS units recorded simultaneously in control and MitoPark mice at each timepoint. **(F)** Mean (± 95% C.I.) noise correlation values for pairs of S1 FS units in control and MitoPark mice for each timepoint. **(G)** Distribution of noise correlations for RS-FS unit pairs (one RS unit and one FS unit recorded simultaneously) in control and MitoPark mice for each timepoint. **(H)** Mean (± 95% C.I.) noise correlation values for RS-FS unit pairs in control and MitoPark mice for each timepoint. RS: regular-spiking units; FS: fast-spiking units; WK: week; CT: control mice; MP: MitoPark mice.

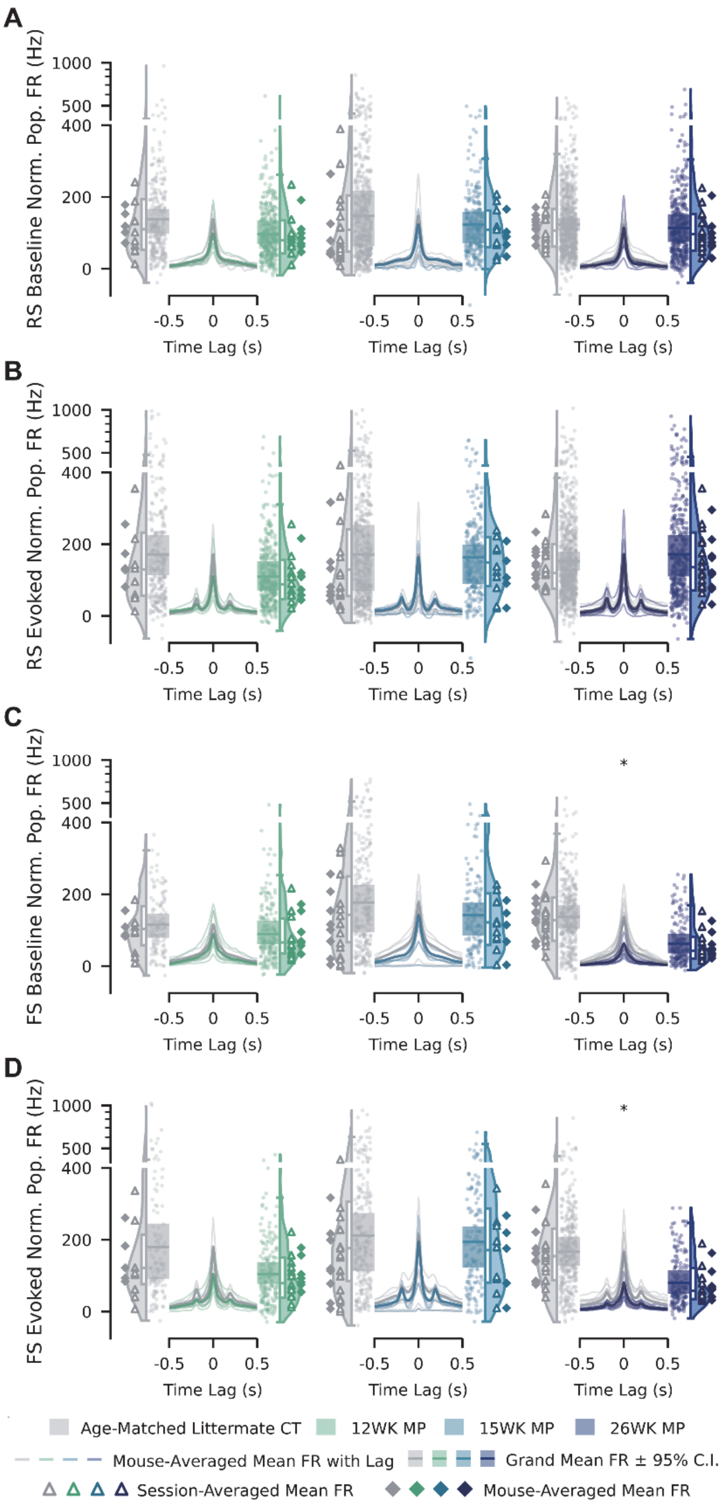


**Figure S6. MitoPark S1 fast-spiking units show decreased population coupling during both baseline and evoked windows at 26 weeks. (A)** Population coupling (calculated as in Figure 6A) for S1 regular-spiking (RS) units during the pre-stimulus baseline window, in control and MitoPark mice at 12 weeks, 15 weeks, and 26 weeks. Left and right raincloud plots show the distribution of peak normalized population firing rates for control and MitoPark, respectively;

violin plots depict the full distribution, with overlaid box plots (median and interquartile range) summarizing dispersion, and points denote individual units. Central line plot shows normalized population firing rate as a function of time lag relative to reference-unit spiking, with thin lines representing mouse-averaged mean coupling traces and the shaded band representing the grand mean ± 95% C.I. Triangles denote session-averaged, and diamonds denote mouse-averaged peak coupling values. **(B)** Similar to (A), but for S1 RS units during the stimulus-evoked window (0 to 1200 ms). **(C)** Similar to (A), but for S1 fast-spiking (FS) units during the pre-stimulus baseline window. **(D)** Similar to (B), but for S1 FS units during the stimulus-evoked window. *$p<0.05$, post hoc simple-effects contrast test for genotype with Bonferroni correction for multiple comparisons. FR: firing rates; WK: week; CT: control mice; MP: MitoPark mice.

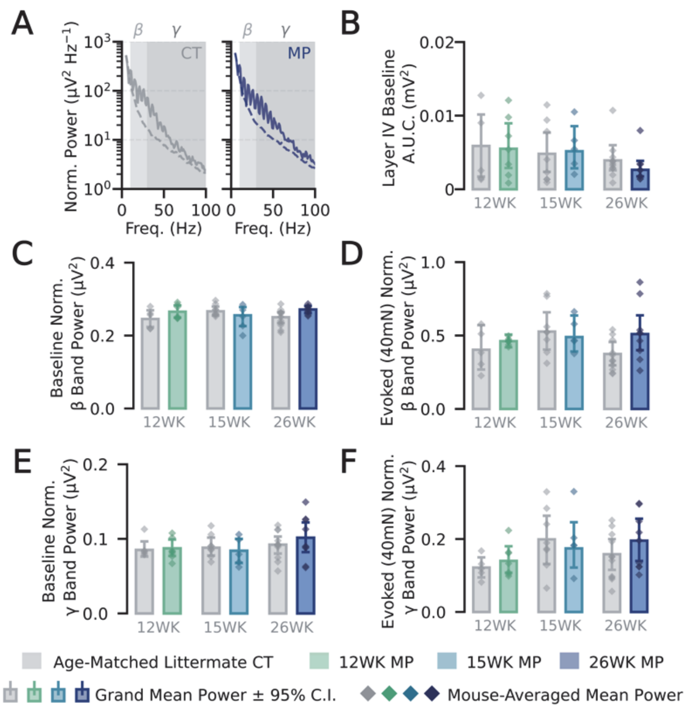


**Figure S7. Intracortical local field potential β and γ band powers are indistinguishable between control and MitoPark mice. (A)** Representative power spectral densities (PSD) of layer IV local field potentials from a 26-week control (left) and 26-week MitoPark (right) mouse. PSDs measured during the pre-stimulus baseline window (dashed) or the stimulus-evoked window (solid, 40 mN). β and γ frequency bands indicated by shading. **(B)** Area under the curve (A.U.C.) of the unnormalized baseline PSD, in control and MitoPark mice at 12 weeks, 15 weeks, and 26 weeks. Bars represent grand mean power ± 95% C.I.; diamonds denote mouse averages. **(C)** Baseline β band power in control and MitoPark for the three timepoints. **(D)** Stimulus-evoked (40 mN) β band power in control and MitoPark mice for each timepoint. **(E)** Similar to (C), but for baseline γ band power. **(F)** Similar to (D), but for evoked γ band power. WK: week; CT: control mice; MP: MitoPark mice.

| 6WK | → | 12WK – 14WK | → | 20WK | → | 30WK |
|---|---|---|---|---|---|---|
| ↓ Mitochondrial DNA expression | | Fewer midbrain TH$^+$ Neurons | | Overt Motor Deficits | | Severe motor deficits |
| Normal Number of TH$^+$ DA Neurons | | Reduced TH$^+$ Striatal Fibers | | L-DOPA ameliorates motor phenotype | | L-DOPA results in peak of activity |
| | | Modest Decrease in Locomotion | | Fewer TH$^+$ cells and fibers | | Fewer TH$^+$ cells and fibers |

| Group | 12WK CT | 12WK MP | 15WK CT | 15WK MP | 26WK CT | 26WK MP | TOTAL |
|---|---|---|---|---|---|---|---|
| # Mice | 5 (2, 3) | 7 (2, 5) | 8 (5, 3) | 5 (5, 0) | 11 (1, 10) | 9 (1, 8) | 45 (16, 29) |
| # Records | 9 | 13 | 17 | 10 | 19 | 16 | 84 |
| # neurons | 377 | 568 | 788 | 377 | 1179 | 617 | 3906 |

**Table S1. Disease progression and recording summary across timepoints in MitoPark mice. Upper:** Development of motor symptoms and pathological hallmarks in MitoPark mice, based on reference (27). **Lower:** Number of mice (male, female), electrophysiological recordings, and well-isolated neurons included in this dataset. WK: week; CT: control mice; MP: MitoPark mice.

**Figure 1 (STR & Midbrain TH Density ):** Two-way ANOVA (genotype × age) with post-hoc simple-effects contrasts on the fitted model (Bonferroni correction).

**Two-way ANOVA (genotype × age) — Striatum**

| Location | Source | SS | df | MS | F | p | sig |
|---|---|---|---|---|---|---|---|
| Striatum | Genotype | 0.075945016 | 1 | 0.075945016 | 5.9127607 | **0.03163891** | * |
| Striatum | C(age, Treatment(ref=12)) | 0.468488154 | 2 | 0.234244077 | 18.2372621 | **0.00023015** | * |
| Striatum | C(genotype_1, Treatment(ref='CT')):C(age, Treatment(ref=12)) | 0.104525167 | 2 | 0.052262584 | 4.06894574 | **0.04477169** | * |
| Striatum | Residual | 0.154131081 | 12 | 0.012844257 | | | ns |

**Post-Hoc simple-effects contracts (CT vs MP) — Striatum**

| Location | Age | A | B | t_stat | p_unc | p_corrected | sig |
|---|---|---|---|---|---|---|---|
| Striatum | 12 | CT | MP | -1.19846003 | 0.25387134 | 0.76161403 | ns |
| Striatum | 15 | CT | MP | 0.502689685 | 0.62428057 | 1 | ns |
| Striatum | 24 | CT | MP | -3.51591365 | 0.00425532 | **0.01276596** | * |

**Two-way ANOVA (genotype × age) — Midbrain**

| Location | Source | SS | df | MS | F | p | sig |
|---|---|---|---|---|---|---|---|
| Midbrain | Genotype | 0.085933272 | 1 | 0.085933272 | 28.0459732 | **0.00018962** | * |
| Midbrain | C(age, Treatment(ref=12)) | 0.609782228 | 2 | 0.304891114 | 99.5070686 | **3.3823E-08** | * |
| Midbrain | C(genotype_1, Treatment(ref='CT')):C(age, Treatment(ref=12)) | 0.076255388 | 2 | 0.038127694 | 12.4437049 | **0.00118527** | * |
| Midbrain | Residual | 0.036768176 | 12 | 0.003064015 | | | ns |

**Post-Hoc simple-effects contracts (CT vs MP) — Midbrain**

| Location | Age | A | B | t_stat | p_unc | p_corrected | sig |
|---|---|---|---|---|---|---|---|
| Midbrain | 12 | CT | MP | 0.371495965 | 0.71674454 | 1 | ns |
| Midbrain | 15 | CT | MP | -2.86816738 | 0.01413783 | **0.04241348** | * |
| Midbrain | 24 | CT | MP | -6.67600103 | 2.2734E-05 | **6.8201E-05** | * |

**Figure 1 (S1 TH Fiber Density [mm/mm^3]):** Two-way ANOVA (genotype × age) with post-hoc simple-effects contrasts on fitted model (Bonferroni correction).

**Two-way ANOVA (genotype × age) — Layer I**

| Location | Source | SS | df | MS | F | p | sig |
|---|---|---|---|---|---|---|---|
| Layer I | Genotype | 2578590.664 | 1 | 2578590.664 | 6.46058991 | **0.0204485** | * |
| Layer I | C(age, Treatment(ref=12)) | 730740.1063 | 2 | 365370.0531 | 0.91542489 | 0.4181967 | ns |
| Layer I | C(genotype_1, Treatment(ref='CT')):C(age, Treatment(ref=12)) | 421296.7834 | 2 | 210648.3917 | 0.52777391 | 0.5987694 | ns |
| Layer I | Residual | 7184271.493 | 18 | 399126.194 | | | ns |

**Post-Hoc simple-effects contracts (CT vs MP) — Layer I**

| Location | Age | A | B | t_stat | p_unc | p_corrected | sig |
|---|---|---|---|---|---|---|---|
| Layer I | 12 | CT | MP | -0.58788654 | 0.56391203 | 1 | ns |
| Layer I | 15 | CT | MP | -0.9609223 | 0.34932277 | 1 | ns |
| Layer I | 26 | CT | MP | -2.49943103 | 0.02233442 | 0.0670033 | ns |

**Two-way ANOVA (genotype × age) — Layer II/III**

| Location | Source | SS | df | MS | F | p | sig |
|---|---|---|---|---|---|---|---|
| Layer II/III | Genotype | 819928.2512 | 1 | 819928.2512 | 6.97646197 | **0.016591** | * |
| Layer II/III | C(age, Treatment(ref=12)) | 118095.8596 | 2 | 59047.92982 | 0.50241669 | 0.6133042 | ns |
| Layer II/III | C(genotype_1, Treatment(ref='CT')):C(age, Treatment(ref=12)) | 565702.0175 | 2 | 282851.0087 | 2.40667315 | 0.1185091 | ns |
| Layer II/III | Residual | 2115500.462 | 18 | 117527.8035 | | | ns |

**Post-Hoc simple-effects contracts (CT vs MP) — Layer II/III**

| Location | Age | A | B | t_stat | p_unc | p_corrected | sig |
|---|---|---|---|---|---|---|---|
| Layer II/III | 12 | CT | MP | -0.33153522 | 0.7440691 | 1 | ns |
| Layer II/III | 15 | CT | MP | -0.12089199 | 0.90511587 | 1 | ns |
| Layer II/III | 26 | CT | MP | -3.41544694 | 0.00308415 | **0.0092525** | * |

**Two-way ANOVA (genotype × age) — Layer IV**

| Location | Source | SS | df | MS | F | p | sig |
|---|---|---|---|---|---|---|---|
| Layer IV | Genotype | 2907714.428 | 1 | 2907714.428 | 9.66600329 | **0.0060586** | * |
| Layer IV | C(age, Treatment(ref=12)) | 132780.9082 | 2 | 66390.45411 | 0.22069923 | 0.804096 | ns |
| Layer IV | C(genotype_1, Treatment(ref='CT')):C(age, Treatment(ref=12)) | 1012951.705 | 2 | 506475.8525 | 1.68365821 | 0.2136499 | ns |
| Layer IV | Residual | 5414736.382 | 18 | 300818.6879 | | | ns |

**Post-Hoc simple-effects contracts (CT vs MP) — Layer IV**

| Location | Age | A | B | t_stat | p_unc | p_corrected | sig |
|---|---|---|---|---|---|---|---|
| Layer IV | 12 | CT | MP | -0.12717202 | 0.90021416 | 1 | ns |
| Layer IV | 15 | CT | MP | -1.37154007 | 0.1870628 | 0.5611884 | ns |
| Layer IV | 26 | CT | MP | -3.3370683 | 0.00366831 | **0.0110049** | * |

**Two-way ANOVA (genotype × age) — Layer V**

| Location | Source | SS | df | MS | F | p | sig |
|---|---|---|---|---|---|---|---|
| Layer V | Genotype | 1589856.283 | 1 | 1589856.283 | 9.33279977 | **0.0068182** | * |
| Layer V | C(age, Treatment(ref=12)) | 245757.7835 | 2 | 122878.8917 | 0.72132564 | 0.4996354 | ns |
| Layer V | C(genotype_1, Treatment(ref='CT')):C(age, Treatment(ref=12)) | 1127424.499 | 2 | 563712.2494 | 3.30911266 | 0.059723 | ns |
| Layer V | Residual | 3066326.696 | 18 | 170351.4831 | | | ns |

**Post-Hoc simple-effects contracts (CT vs MP) — Layer V**

| Location | Age | A | B | t_stat | p_unc | p_corrected | sig |
|---|---|---|---|---|---|---|---|
| Layer V | 12 | CT | MP | 0.359942001 | 0.72307913 | 1 | ns |
| Layer V | 15 | CT | MP | -1.04055774 | 0.31185672 | 0.9355702 | ns |
| Layer V | 26 | CT | MP | -3.83910229 | 0.00120195 | **0.0036059** | * |

**Two-way ANOVA (genotype × age) — Layer VI**

| Location | Source | SS | df | MS | F | p | sig |
|---|---|---|---|---|---|---|---|
| Layer VI | Genotype | 2402032.785 | 1 | 2402032.785 | 11.1119806 | **0.0036977** | * |
| Layer VI | C(age, Treatment(ref=12)) | 130400.1866 | 2 | 65200.09328 | 0.30162043 | 0.7432845 | ns |
| Layer VI | C(genotype_1, Treatment(ref='CT')):C(age, Treatment(ref=12)) | 783944.1795 | 2 | 391972.0898 | 1.81329176 | 0.1916732 | ns |
| Layer VI | Residual | 3890988.625 | 18 | 216166.0347 | | | ns |

**Post-Hoc simple-effects contracts (CT vs MP) — Layer VI**

| Location | Age | A | B | t_stat | p_unc | p_corrected | sig |
|---|---|---|---|---|---|---|---|
| Layer VI | 12 | CT | MP | -0.56422103 | 0.57956701 | 1 | ns |
| Layer VI | 15 | CT | MP | -0.87783977 | 0.39159761 | 1 | ns |
| Layer VI | 26 | CT | MP | -3.69453869 | 0.00165871 | **0.0049761** | * |

**Figure 2 (Baseline and Evoked Firing Rates):** LME Models: One table per cell type x firing-rate condition (baseline/evoked) x cortical layer (16 models total).

**LME — RS | Baseline | Layer II/III [N obs = 228, N groups = 37, Log-Likelihood = -665.206, Scale(sigma^2) = 18.44732, Method = REML, Converged = Yes]**

| Term | Beta (Coef.) | SE | z | p | 95% CI Low | 95% CI High | sig |
|---|---|---|---|---|---|---|---|
| Intercept (CT, 12W, reference sex) | 5.7838 | 1.4919 | 3.877 | 0.0001 | 2.8597 | 8.7079 | |
| Age: 15W (vs 12W, reference) | -0.6783 | 2.2018 | -0.308 | 0.758 | -4.9937 | 3.6371 | ns |
| Age: 26W (vs 12W, reference) | -2.8218 | 1.7061 | -1.654 | 0.0981 | -6.1658 | 0.5222 | ns |
| Group: MP (vs CT, reference) | -2.0052 | 2.1055 | -0.952 | 0.3409 | -6.1318 | 2.1214 | ns |
| Sex: Male (vs Female, reference) | -1.1196 | 1.7921 | -0.625 | 0.5322 | -4.632 | 2.3929 | ns |
| Interaction: Age 15W x Group MP | 3.1615 | 2.732 | 1.157 | 0.2472 | -2.1931 | 8.5161 | ns |
| Interaction: Age 26W x Group MP | 3.3915 | 2.4718 | 1.372 | 0.17 | -1.4531 | 8.2361 | ns |
| Random Effect: Mouse Intercept Variance | 0 | 0.1715 | 0 | 1 | -0.3362 | 0.3362 | |
| Random Effect: Session Variance Component (nested within Mouse) | 0.3158 | 0.1838 | 1.718 | 0.0859 | -0.0446 | 0.6761 | |

**LME — RS | Baseline | Layer IV [N obs = 267, N groups = 39, Log-Likelihood = -823.334, Scale(sigma^2) = 23.15011, Method = REML, Converged = Yes]**

| Term | Beta (Coef.) | SE | z | p | 95% CI Low | 95% CI High | sig |
|---|---|---|---|---|---|---|---|
| Intercept (CT, 12W, reference sex) | 9.3603 | 3.3678 | 2.779 | 0.0054 | 2.7596 | 15.961 | |
| Age: 15W (vs 12W, reference) | -3.0634 | 4.1012 | -0.747 | 0.4551 | -11.1016 | 4.9748 | ns |
| Age: 26W (vs 12W, reference) | -6.5054 | 3.5945 | -1.81 | 0.0703 | -13.5504 | 0.5397 | ns |
| Group: MP (vs CT, reference) | -5.8465 | 3.9024 | -1.498 | 0.1341 | -13.4951 | 1.8021 | ns |
| Sex: Male (vs Female, reference) | -2.5323 | 2.3675 | -1.07 | 0.2848 | -7.1725 | 2.1079 | ns |
| Interaction: Age 15W x Group MP | 10.123 | 5.0728 | 1.996 | **0.046** | 0.1805 | 20.0654 | * |
| Interaction: Age 26W x Group MP | 9.6584 | 4.6833 | 2.062 | **0.0392** | 0.4794 | 18.8375 | * |
| Random Effect: Mouse Intercept Variance | 0 | 1.484 | 0 | 1 | -2.9086 | 2.9086 | |
| Random Effect: Session Variance Component (nested within Mouse) | 0.8171 | 0.8849 | 0.923 | 0.3558 | -0.9173 | 2.5515 | |

**LME — RS | Baseline | Layer V [N obs = 1746, N groups = 45, Log-Likelihood = -6081.854, Scale(sigma^2) = 60.14072, Method = REML, Converged = Yes]**

| Term | Beta (Coef.) | SE | z | p | 95% CI Low | 95% CI High | sig |
|---|---|---|---|---|---|---|---|
| Intercept (CT, 12W, reference sex) | 6.716 | 1.0636 | 6.314 | 0 | 4.6313 | 8.8007 | |
| Age: 15W (vs 12W, reference) | 0.6538 | 1.2828 | 0.51 | 0.6103 | -1.8604 | 3.1679 | ns |
| Age: 26W (vs 12W, reference) | -1.6092 | 1.2238 | -1.315 | 0.1885 | -4.0078 | 0.7894 | ns |
| Group: MP (vs CT, reference) | 0.0612 | 1.2878 | 0.048 | 0.9621 | -2.4628 | 2.5852 | ns |
| Sex: Male (vs Female, reference) | -1.5125 | 0.956 | -1.582 | 0.1136 | -3.3862 | 0.3611 | ns |
| Interaction: Age 15W x Group MP | 3.9367 | 1.8231 | 2.159 | **0.0308** | 0.3635 | 7.51 | * |
| Interaction: Age 26W x Group MP | 3.097 | 1.6195 | 1.912 | 0.0558 | -0.0772 | 6.2711 | ns |
| Random Effect: Mouse Intercept Variance | 0.0075 | 0.0217 | 0.348 | 0.7281 | -0.0349 | 0.05 | |
| Random Effect: Session Variance Component (nested within Mouse) | 0.068 | 0.0285 | 2.391 | 0.0168 | 0.0123 | 0.1238 | |

**LME — RS | Baseline | Layer VI [N obs = 456, N groups = 40, Log-Likelihood = -1354.063, Scale(sigma^2) = 21.65094, Method = REML, Converged = Yes]**

| Term | Beta (Coef.) | SE | z | p | 95% CI Low | 95% CI High | sig |
|---|---|---|---|---|---|---|---|
| Intercept (CT, 12W, reference sex) | 3.0307 | 1.0036 | 3.02 | 0.0025 | 1.0637 | 4.9978 | |
| Age: 15W (vs 12W, reference) | 0.6591 | 1.303 | 0.506 | 0.613 | -1.8947 | 3.2129 | ns |
| Age: 26W (vs 12W, reference) | -0.4946 | 1.1734 | -0.422 | 0.6734 | -2.7945 | 1.8053 | ns |
| Group: MP (vs CT, reference) | -0.2936 | 1.2376 | -0.237 | 0.8125 | -2.7193 | 2.1321 | ns |
| Sex: Male (vs Female, reference) | 0.1284 | 1.1135 | 0.115 | 0.9082 | -2.0541 | 2.3109 | ns |
| Interaction: Age 15W x Group MP | 0.2185 | 1.7395 | 0.126 | 0.9 | -3.1907 | 3.6278 | ns |
| Interaction: Age 26W x Group MP | 1.7391 | 1.5534 | 1.12 | 0.2629 | -1.3055 | 4.7837 | ns |
| Random Effect: Mouse Intercept Variance | 0.0613 | 0.0559 | 1.098 | 0.2724 | -0.0482 | 0.1708 | |
| Random Effect: Session Variance Component (nested within Mouse) | 0.0277 | 0.0574 | 0.483 | 0.6292 | -0.0849 | 0.1403 | |

**LME — RS | Evoked | Layer II/III [N obs = 228, N groups = 37, Log-Likelihood = -1121.137, Scale(sigma^2) = 1217.17766, Method = REML, Converged = Yes]**

| Term | Beta (Coef.) | SE | z | p | 95% CI Low | 95% CI High | sig |
|---|---|---|---|---|---|---|---|
| Intercept (CT, 12W, reference sex) | 39.1982 | 10.2918 | 3.809 | 0.0001 | 19.0266 | 59.3698 | |
| Age: 15W (vs 12W, reference) | -3.6303 | 15.6925 | -0.231 | 0.8171 | -34.3871 | 27.1265 | ns |
| Age: 26W (vs 12W, reference) | -2.8022 | 11.6461 | -0.241 | 0.8099 | -25.6282 | 20.0238 | ns |
| Group: MP (vs CT, reference) | -19.9663 | 14.8154 | -1.348 | 0.1778 | -49.0039 | 9.0714 | ns |
| Sex: Male (vs Female, reference) | 0.5483 | 12.989 | 0.042 | 0.9663 | -24.9097 | 26.0064 | ns |
| Interaction: Age 15W x Group MP | 30.187 | 19.1745 | 1.574 | 0.1154 | -7.3943 | 67.7683 | ns |
| Interaction: Age 26W x Group MP | 22.7884 | 17.3486 | 1.314 | 0.189 | -11.2143 | 56.791 | ns |
| Random Effect: Mouse Intercept Variance | 0 | 0.105 | 0 | 1 | -0.2058 | 0.2058 | |
| Random Effect: Session Variance Component (nested within Mouse) | 0.1594 | 0.1176 | 1.356 | 0.1751 | -0.071 | 0.3898 | |

**LME — RS | Evoked | Layer IV [N obs = 267, N groups = 39, Log-Likelihood = -1274.348, Scale(sigma^2) = 773.30214, Method = REML, Converged = Yes]**

| Term | Beta (Coef.) | SE | z | p | 95% CI Low | 95% CI High | sig |
|---|---|---|---|---|---|---|---|
| Intercept (CT, 12W, reference sex) | 49.521 | 16.3844 | 3.022 | 0.0025 | 17.4082 | 81.6338 | |

| | | | | | | | |
|---|---|---|---|---|---|---|---|
| Age: 15W (vs 12W, reference) | -23.4611 | 22.3597 | -1.049 | 0.2941 | -67.2854 | 20.3631 | ns |
| Age: 26W (vs 12W, reference) | -23.2725 | 17.3695 | -1.34 | 0.1803 | -57.316 | 10.7711 | ns |
| Group: MP (vs CT, reference) | -28.0913 | 20.8101 | -1.35 | 0.1771 | -68.8784 | 12.6958 | ns |
| Sex: Male (vs Female, reference) | -2.3277 | 13.4623 | -0.173 | 0.8627 | -28.7133 | 24.0579 | ns |
| Interaction: Age 15W x Group MP | 49.4093 | 25.2257 | 1.959 | 0.0501 | -0.0322 | 98.8509 | ns |
| Interaction: Age 26W x Group MP | 50.7681 | 28.0391 | 1.811 | 0.0702 | -4.1876 | 105.7238 | ns |
| Random Effect: Mouse Intercept Variance | 0 | 1.433 | 0 | 1 | -2.8086 | 2.8086 | |
| Random Effect: Session Variance Component (nested within Mouse) | 0.5998 | 1.1322 | 0.53 | 0.5963 | -1.6193 | 2.819 | |

**LME — RS | Evoked | Layer V [N obs = 1746, N groups = 45, Log-Likelihood = -8806.89, Scale(sigma^2) = 1359.33210, Method = REML, Converged = Yes]**

| Term | Beta (Coef.) | SE | z | p | 95% CI Low | 95% CI High | sig |
|---|---|---|---|---|---|---|---|
| Intercept (CT, 12W, reference sex) | 36.0862 | 6.8226 | 5.289 | 0 | 22.7142 | 49.4582 | |
| Age: 15W (vs 12W, reference) | 8.1306 | 8.3425 | 0.975 | 0.3298 | -8.2203 | 24.4816 | ns |
| Age: 26W (vs 12W, reference) | -2.6109 | 7.9076 | -0.33 | 0.7413 | -18.1096 | 12.8877 | ns |
| Group: MP (vs CT, reference) | -0.4844 | 8.3252 | -0.058 | 0.9536 | -16.8015 | 15.8327 | ns |
| Sex: Male (vs Female, reference) | -5.4126 | 6.0343 | -0.897 | 0.3697 | -17.2397 | 6.4144 | ns |
| Interaction: Age 15W x Group MP | 18.6431 | 11.9145 | 1.565 | 0.1176 | -4.709 | 41.9952 | ns |
| Interaction: Age 26W x Group MP | 23.7589 | 10.5022 | 2.262 | **0.0237** | 3.1749 | 44.3429 | * |
| Random Effect: Mouse Intercept Variance | 0.0615 | 0.0383 | 1.605 | 0.1085 | -0.0136 | 0.1367 | |
| Random Effect: Session Variance Component (nested within Mouse) | 0.0877 | 0.0326 | 2.688 | 0.0072 | 0.0237 | 0.1516 | |

**LME — RS | Evoked | Layer VI [N obs = 456, N groups = 40, Log-Likelihood = -2028.795, Scale(sigma^2) = 423.52862, Method = REML, Converged = Yes]**

| Term | Beta (Coef.) | SE | z | p | 95% CI Low | 95% CI High | sig |
|---|---|---|---|---|---|---|---|
| Intercept (CT, 12W, reference sex) | 17.4017 | 4.5729 | 3.805 | 0.0001 | 8.439 | 26.3643 | |
| Age: 15W (vs 12W, reference) | -0.151 | 5.8398 | -0.026 | 0.9794 | -11.5969 | 11.2948 | ns |
| Age: 26W (vs 12W, reference) | -0.3662 | 5.3458 | -0.069 | 0.9454 | -10.8437 | 10.1113 | ns |
| Group: MP (vs CT, reference) | 1.3094 | 5.6434 | 0.232 | 0.8165 | -9.7515 | 12.3702 | ns |
| Sex: Male (vs Female, reference) | 1.5249 | 5.0257 | 0.303 | 0.7616 | -8.3253 | 11.3751 | ns |
| Interaction: Age 15W x Group MP | 10.5762 | 7.7194 | 1.37 | 0.1707 | -4.5536 | 25.706 | ns |
| Interaction: Age 26W x Group MP | 4.322 | 7.0844 | 0.61 | 0.5418 | -9.5632 | 18.2071 | ns |
| Random Effect: Mouse Intercept Variance | 0 | 0.0597 | 0 | 1 | -0.117 | 0.117 | |
| Random Effect: Session Variance Component (nested within Mouse) | 0.1409 | 0.0675 | 2.087 | 0.0369 | 0.0086 | 0.2731 | |

**LME — FS | Baseline | Layer II/III [N obs = 181, N groups = 38, Log-Likelihood = -598.313, Scale(sigma^2) = 45.90966, Method = REML, Converged = No]**

| Term | Beta (Coef.) | SE | z | p | 95% CI Low | 95% CI High | sig |
|---|---|---|---|---|---|---|---|
| Intercept (CT, 12W, reference sex) | 4.8327 | 2.7108 | 1.783 | 0.0746 | -0.4804 | 10.1458 | |
| Age: 15W (vs 12W, reference) | 5.5991 | 3.481 | 1.608 | 0.1077 | -1.2236 | 12.4217 | ns |
| Age: 26W (vs 12W, reference) | 2.9835 | 2.9161 | 1.023 | 0.3062 | -2.7319 | 8.6989 | ns |
| Group: MP (vs CT, reference) | 0.4408 | 3.3101 | 0.133 | 0.8941 | -6.0469 | 6.9285 | ns |
| Sex: Male (vs Female, reference) | -0.8671 | 2.0248 | -0.428 | 0.6685 | -4.8356 | 3.1015 | ns |
| Interaction: Age 15W x Group MP | -3.8554 | 4.4294 | -0.87 | 0.3841 | -12.5368 | 4.826 | ns |
| Interaction: Age 26W x Group MP | -2.2085 | 3.8719 | -0.57 | 0.5684 | -9.7974 | 5.3803 | ns |
| Random Effect: Mouse Intercept Variance | 0.1168 | | | | | | |
| Random Effect: Session Variance Component (nested within Mouse) | 0.0245 | 0.0789 | 0.311 | 0.7556 | -0.13 | 0.1791 | |

**LME — FS | Baseline | Layer: Layer IV [N obs = 129, N groups = 37, Log-Likelihood = -431.826, Scale(sigma^2) = 47.55350, Method = REML, Converged = Yes]**

| Term | Beta (Coef.) | SE | z | p | 95% CI Low | 95% CI High | sig |
|---|---|---|---|---|---|---|---|
| Intercept (CT, 12W, reference sex) | 7.4269 | 2.8725 | 2.585 | 0.0097 | 1.7969 | 13.0569 | |
| Age: 15W (vs 12W, reference) | -1.7799 | 3.7245 | -0.478 | 0.6327 | -9.0798 | 5.5201 | ns |
| Age: 26W (vs 12W, reference) | -0.4824 | 3.3044 | -0.146 | 0.8839 | -6.959 | 5.9942 | ns |
| Group: MP (vs CT, reference) | -3.4764 | 3.5334 | -0.984 | 0.3252 | -10.4017 | 3.449 | ns |
| Sex: Male (vs Female, reference) | 3.0965 | 2.7551 | 1.124 | 0.2611 | -2.3034 | 8.4964 | ns |
| Interaction: Age 15W x Group MP | 5.4579 | 5.5552 | 0.982 | 0.3259 | -5.4301 | 16.3459 | ns |
| Interaction: Age 26W x Group MP | 6.9358 | 4.4994 | 1.541 | 0.1232 | -1.8828 | 15.7543 | ns |
| Random Effect: Mouse Intercept Variance | 0.2077 | 0.2643 | 0.786 | 0.4319 | -0.3103 | 0.7257 | |
| Random Effect: Session Variance Component (nested within Mouse) | 0.1546 | 0.2215 | 0.698 | 0.4851 | -0.2795 | 0.5887 | |

**LME — FS | Baseline | Layer V [N obs = 563, N groups = 45, Log-Likelihood = -2130.261, Scale(sigma^2) = 106.53711, Method = REML, Converged = Yes]**

| Term | Beta (Coef.) | SE | z | p | 95% CI Low | 95% CI High | sig |
|---|---|---|---|---|---|---|---|
| Intercept (CT, 12W, reference sex) | 8.6095 | 2.333 | 3.69 | 0.0002 | 4.0369 | 13.1822 | |
| Age: 15W (vs 12W, reference) | -0.255 | 2.8691 | -0.089 | 0.9292 | -5.8783 | 5.3683 | ns |
| Age: 26W (vs 12W, reference) | 0.0993 | 2.7217 | 0.036 | 0.9709 | -5.2352 | 5.4338 | ns |
| Group: MP (vs CT, reference) | -1.4074 | 2.8972 | -0.486 | 0.6271 | -7.0859 | 4.2711 | ns |
| Sex: Male (vs Female, reference) | -0.0643 | 2.1042 | -0.031 | 0.9756 | -4.1884 | 4.0599 | ns |
| Interaction: Age 15W x Group MP | 3.0205 | 4.1095 | 0.735 | 0.4623 | -5.0339 | 11.0749 | ns |

| | | | | | | | |
|---|---|---|---|---|---|---|---|
| Interaction: Age 26W x Group MP | -0.9136 | 3.6438 | -0.251 | 0.802 | -8.0553 | 6.228 | ns |
| Random Effect: Mouse Intercept Variance | 0.0596 | | | | | | |
| Random Effect: Session Variance Component (nested within Mouse) | 0.1295 | 0.1012 | 1.279 | 0.201 | -0.069 | 0.3279 | |

**LME — FS | Baseline | Layer VI [N obs = 336, N groups = 44, Log-Likelihood = -1087.214, Scale(sigma^2) = 33.78300, Method = REML, Converged = Yes]**

| Term | Beta (Coef.) | SE | z | p | 95% CI Low | 95% CI High | sig |
|---|---|---|---|---|---|---|---|
| Intercept (CT, 12W, reference sex) | 4.5788 | 1.6241 | 2.819 | 0.0048 | 1.3955 | 7.762 | |
| Age: 15W (vs 12W, reference) | -1.5249 | 1.985 | -0.768 | 0.4424 | -5.4154 | 2.3657 | ns |
| Age: 26W (vs 12W, reference) | 1.9283 | 1.8937 | 1.018 | 0.3086 | -1.7834 | 5.64 | ns |
| Group: MP (vs CT, reference) | 0.1728 | 1.985 | 0.087 | 0.9306 | -3.7177 | 4.0633 | ns |
| Sex: Male (vs Female, reference) | 2.4286 | 1.464 | 1.659 | 0.0971 | -0.4408 | 5.2981 | ns |
| Interaction: Age 15W x Group MP | 1.529 | 2.7739 | 0.551 | 0.5815 | -3.9078 | 6.9658 | ns |
| Interaction: Age 26W x Group MP | -2.6335 | 2.5222 | -1.044 | 0.2964 | -7.5769 | 2.3099 | ns |
| Random Effect: Mouse Intercept Variance | 0 | 0.116 | 0 | 1 | -0.2274 | 0.2274 | |
| Random Effect: Session Variance Component (nested within Mouse) | 0.3053 | 0.1583 | 1.929 | 0.0537 | -0.0049 | 0.6156 | |

**LME — FS | Evoked | Layer II/III [N obs = 181, N groups = 38, Log-Likelihood = -942.254, Scale(sigma^2) = 2030.18746, Method = REML, Converged = Yes]**

| Term | Beta (Coef.) | SE | z | p | 95% CI Low | 95% CI High | sig |
|---|---|---|---|---|---|---|---|
| Intercept (CT, 12W, reference sex) | 79.5998 | 28.6232 | 2.781 | 0.0054 | 23.4994 | 135.7002 | |
| Age: 15W (vs 12W, reference) | -16.2729 | 35.294 | -0.461 | 0.6447 | -85.4479 | 52.902 | ns |
| Age: 26W (vs 12W, reference) | -15.7521 | 31.2683 | -0.504 | 0.6144 | -77.0368 | 45.5325 | ns |
| Group: MP (vs CT, reference) | -40.7727 | 34.2534 | -1.19 | 0.2339 | -107.9082 | 26.3627 | ns |
| Sex: Male (vs Female, reference) | 14.4176 | 20.3569 | 0.708 | 0.4788 | -25.4811 | 54.3164 | ns |
| Interaction: Age 15W x Group MP | 58.2841 | 45.3112 | 1.286 | 0.1983 | -30.5243 | 147.0924 | ns |
| Interaction: Age 26W x Group MP | 54.6021 | 39.9214 | 1.368 | 0.1714 | -23.6423 | 132.8465 | ns |
| Random Effect: Mouse Intercept Variance | 0.5428 | 0.2842 | 1.91 | 0.0562 | -0.0143 | 1.0999 | |
| Random Effect: Session Variance Component (nested within Mouse) | 0.1158 | 0.1331 | 0.87 | 0.3841 | -0.145 | 0.3767 | |

**LME — FS | Evoked | Layer IV [N obs = 129, N groups = 37, Log-Likelihood = -650.489, Scale(sigma^2) = 1787.49902, Method = REML, Converged = Yes]**

| Term | Beta (Coef.) | SE | z | p | 95% CI Low | 95% CI High | sig |
|---|---|---|---|---|---|---|---|
| Intercept (CT, 12W, reference sex) | 66.4396 | 15.8044 | 4.204 | 0 | 35.4635 | 97.4158 | |
| Age: 15W (vs 12W, reference) | -19.2303 | 20.1859 | -0.953 | 0.3408 | -58.7939 | 20.3333 | ns |
| Age: 26W (vs 12W, reference) | -0.2796 | 18.0477 | -0.015 | 0.9876 | -35.6524 | 35.0932 | ns |
| Group: MP (vs CT, reference) | -23.0222 | 19.345 | -1.19 | 0.234 | -60.9378 | 14.8934 | ns |
| Sex: Male (vs Female, reference) | -6.0746 | 14.4761 | -0.42 | 0.6748 | -34.4472 | 22.2979 | ns |
| Interaction: Age 15W x Group MP | 38.6592 | 28.4291 | 1.36 | 0.1739 | -17.0607 | 94.3792 | ns |
| Interaction: Age 26W x Group MP | 41.4433 | 24.3958 | 1.699 | 0.0894 | -6.3716 | 89.2582 | ns |
| Random Effect: Mouse Intercept Variance | 0.0366 | 0.1699 | 0.216 | 0.8293 | -0.2964 | 0.3697 | |
| Random Effect: Session Variance Component (nested within Mouse) | 0.2101 | 0.2462 | 0.853 | 0.3935 | -0.2725 | 0.6927 | |

**LME — FS | Evoked | Layer V [N obs = 563, N groups = 45, Log-Likelihood = -2999.394, Scale(sigma^2) = 2438.33908, Method = REML, Converged = Yes]**

| Term | Beta (Coef.) | SE | z | p | 95% CI Low | 95% CI High | sig |
|---|---|---|---|---|---|---|---|
| Intercept (CT, 12W, reference sex) | 54.0101 | 11.4895 | 4.701 | 0 | 31.4911 | 76.5291 | |
| Age: 15W (vs 12W, reference) | 8.7512 | 14.1484 | 0.619 | 0.5362 | -18.9791 | 36.4816 | ns |
| Age: 26W (vs 12W, reference) | 5.6493 | 13.4178 | 0.421 | 0.6737 | -20.6491 | 31.9476 | ns |
| Group: MP (vs CT, reference) | -2.2186 | 14.2927 | -0.155 | 0.8766 | -30.2317 | 25.7946 | ns |
| Sex: Male (vs Female, reference) | 4.4475 | 10.5845 | 0.42 | 0.6743 | -16.2978 | 25.1927 | ns |
| Interaction: Age 15W x Group MP | 19.3842 | 20.3189 | 0.954 | 0.3401 | -20.4402 | 59.2085 | ns |
| Interaction: Age 26W x Group MP | -8.3111 | 18.165 | -0.458 | 0.6473 | -43.9138 | 27.2917 | ns |
| Random Effect: Mouse Intercept Variance | 0.0887 | 0.0815 | 1.088 | 0.2767 | -0.0711 | 0.2485 | |
| Random Effect: Session Variance Component (nested within Mouse) | 0.0984 | 0.1047 | 0.939 | 0.3475 | -0.1069 | 0.3036 | |

**LME — FS | Evoked | Layer VI [N obs = 336, N groups = 44, Log-Likelihood = -1704.807, Scale(sigma^2) = 1513.96560, Method = REML, Converged = Yes]**

| Term | Beta (Coef.) | SE | z | p | 95% CI Low | 95% CI High | sig |
|---|---|---|---|---|---|---|---|
| Intercept (CT, 12W, reference sex) | 31.5314 | 10.5269 | 2.995 | 0.0027 | 10.899 | 52.1637 | |
| Age: 15W (vs 12W, reference) | -10.7788 | 13.0224 | -0.828 | 0.4078 | -36.3023 | 14.7447 | ns |
| Age: 26W (vs 12W, reference) | 9.5645 | 12.3124 | 0.777 | 0.4373 | -14.5673 | 33.6963 | ns |
| Group: MP (vs CT, reference) | -11.2504 | 12.9053 | -0.872 | 0.3833 | -36.5443 | 14.0435 | ns |
| Sex: Male (vs Female, reference) | 19.78 | 9.4267 | 2.098 | **0.0359** | 1.304 | 38.2561 | * |
| Interaction: Age 15W x Group MP | 25.8653 | 18.2401 | 1.418 | 0.1562 | -9.8847 | 61.6152 | ns |
| Interaction: Age 26W x Group MP | -7.2581 | 16.3801 | -0.443 | 0.6577 | -39.3625 | 24.8462 | ns |
| Random Effect: Mouse Intercept Variance | 0.0832 | 0.0724 | 1.15 | 0.2501 | -0.0586 | 0.2251 | |
| Random Effect: Session Variance Component (nested within Mouse) | 0.1214 | 0.0835 | 1.453 | 0.1463 | -0.0424 | 0.2851 | |

**Figure 2 (Baseline and Evoked Firing Rates):** model-based simple-effects contrasts of genotype at each age, with Bonferroni correction applied across the 3 ages.

**Post-hoc simple-effects contrasts (CT vs MP) — Regular-Spiking (RS), Baseline, Layer II/III (Bonferroni correction across 3 ages)**

| Cell Type | Condition | Layer | Age (W) | Comparison | Estimate (Hz) | t_stat | p_raw | p_corrected | sig |
|---|---|---|---|---|---|---|---|---|---|
| Regular-Spiking (RS) | Baseline | Layer II/III | 12 | CT vs MP | -2.0052 | -0.952 | 0.340908 | 1 | ns |
| Regular-Spiking (RS) | Baseline | Layer II/III | 15 | CT vs MP | 1.1563 | 0.623 | 0.533061 | 1 | ns |
| Regular-Spiking (RS) | Baseline | Layer II/III | 26 | CT vs MP | 1.3863 | 1.068 | 0.285643 | 0.85693 | ns |

**Post-hoc simple-effects contrasts (CT vs MP) — Regular-Spiking (RS), Baseline, Layer IV (Bonferroni correction across 3 ages)**

| Cell Type | Condition | Layer | Age (W) | Comparison | Estimate (Hz) | t_stat | p_raw | p_corrected | sig |
|---|---|---|---|---|---|---|---|---|---|
| Regular-Spiking (RS) | Baseline | Layer IV | 12 | CT vs MP | -5.8465 | -1.498 | 0.134087 | 0.402262 | ns |
| Regular-Spiking (RS) | Baseline | Layer IV | 15 | CT vs MP | 4.2765 | 1.429 | 0.153142 | 0.459425 | ns |
| Regular-Spiking (RS) | Baseline | Layer IV | 26 | CT vs MP | 3.8119 | 1.768 | 0.076989 | 0.230966 | ns |

**Post-hoc simple-effects contrasts (CT vs MP) — Regular-Spiking (RS), Baseline, Layer V (Bonferroni correction across 3 ages)**

| Cell Type | Condition | Layer | Age (W) | Comparison | Estimate (Hz) | t_stat | p_raw | p_corrected | sig |
|---|---|---|---|---|---|---|---|---|---|
| Regular-Spiking (RS) | Baseline | Layer V | 12 | CT vs MP | 0.0612 | 0.048 | 0.962077 | 1 | ns |
| Regular-Spiking (RS) | Baseline | Layer V | 15 | CT vs MP | 3.998 | 3.127 | 0.001763 | **0.00529** | * |
| Regular-Spiking (RS) | Baseline | Layer V | 26 | CT vs MP | 3.1582 | 3.226 | 0.001254 | **0.003762** | * |

**Post-hoc simple-effects contrasts (CT vs MP) — Regular-Spiking (RS), Baseline, Layer VI (Bonferroni correction across 3 ages)**

| Cell Type | Condition | Layer | Age (W) | Comparison | Estimate (Hz) | t_stat | p_raw | p_corrected | sig |
|---|---|---|---|---|---|---|---|---|---|
| Regular-Spiking (RS) | Baseline | Layer VI | 12 | CT vs MP | -0.2936 | -0.237 | 0.812484 | 1 | ns |
| Regular-Spiking (RS) | Baseline | Layer VI | 15 | CT vs MP | -0.0751 | -0.062 | 0.950285 | 1 | ns |
| Regular-Spiking (RS) | Baseline | Layer VI | 26 | CT vs MP | 1.4455 | 1.538 | 0.123994 | 0.371983 | ns |

**Post-hoc simple-effects contrasts (CT vs MP) — Regular-Spiking (RS), Evoked, Layer II/III (Bonferroni correction across 3 ages)**

| Cell Type | Condition | Layer | Age (W) | Comparison | Estimate (Hz) | t_stat | p_raw | p_corrected | sig |
|---|---|---|---|---|---|---|---|---|---|
| Regular-Spiking (RS) | Evoked | Layer II/III | 12 | CT vs MP | -19.9663 | -1.348 | 0.177765 | 0.533296 | ns |
| Regular-Spiking (RS) | Evoked | Layer II/III | 15 | CT vs MP | 10.2207 | 0.79 | 0.429656 | 1 | ns |
| Regular-Spiking (RS) | Evoked | Layer II/III | 26 | CT vs MP | 2.8221 | 0.313 | 0.754116 | 1 | ns |

**Post-hoc simple-effects contrasts (CT vs MP) — Regular-Spiking (RS), Evoked, Layer IV (Bonferroni correction across 3 ages)**

| Cell Type | Condition | Layer | Age (W) | Comparison | Estimate (Hz) | t_stat | p_raw | p_corrected | sig |
|---|---|---|---|---|---|---|---|---|---|
| Regular-Spiking (RS) | Evoked | Layer IV | 12 | CT vs MP | -28.0913 | -1.35 | 0.177052 | 0.531157 | ns |
| Regular-Spiking (RS) | Evoked | Layer IV | 15 | CT vs MP | 21.318 | 1.45 | 0.147038 | 0.441113 | ns |
| Regular-Spiking (RS) | Evoked | Layer IV | 26 | CT vs MP | 22.6768 | 1.602 | 0.1091 | 0.327301 | ns |

**Post-hoc simple-effects contrasts (CT vs MP) — Regular-Spiking (RS), Evoked, Layer V (Bonferroni correction across 3 ages)**

| Cell Type | Condition | Layer | Age (W) | Comparison | Estimate (Hz) | t_stat | p_raw | p_corrected | sig |
|---|---|---|---|---|---|---|---|---|---|
| Regular-Spiking (RS) | Evoked | Layer V | 12 | CT vs MP | -0.4844 | -0.058 | 0.953604 | 1 | ns |
| Regular-Spiking (RS) | Evoked | Layer V | 15 | CT vs MP | 18.1587 | 2.158 | 0.030959 | 0.092877 | ns |
| Regular-Spiking (RS) | Evoked | Layer V | 26 | CT vs MP | 23.2745 | 3.645 | 0.000268 | **0.000803** | * |

**Post-hoc simple-effects contrasts (CT vs MP) — Regular-Spiking (RS), Evoked, Layer VI (Bonferroni correction across 3 ages)**

| Cell Type | Condition | Layer | Age (W) | Comparison | Estimate (Hz) | t_stat | p_raw | p_corrected | sig |
|---|---|---|---|---|---|---|---|---|---|
| Regular-Spiking (RS) | Evoked | Layer VI | 12 | CT vs MP | 1.3094 | 0.232 | 0.816524 | 1 | ns |
| Regular-Spiking (RS) | Evoked | Layer VI | 15 | CT vs MP | 11.8856 | 2.266 | 0.023471 | 0.070412 | ns |
| Regular-Spiking (RS) | Evoked | Layer VI | 26 | CT vs MP | 5.6313 | 1.321 | 0.186589 | 0.559766 | ns |

**Post-hoc simple-effects contrasts (CT vs MP) — Fast-Spiking (FS), Baseline, Layer II/III (Bonferroni correction across 3 ages)**

| Cell Type | Condition | Layer | Age (W) | Comparison | Estimate (Hz) | t_stat | p_raw | p_corrected | sig |
|---|---|---|---|---|---|---|---|---|---|
| Fast-Spiking (FS) | Baseline | Layer II/III | 12 | CT vs MP | 0.4408 | 0.133 | 0.894061 | 1 | ns |
| Fast-Spiking (FS) | Baseline | Layer II/III | 15 | CT vs MP | -3.4146 | -1.133 | 0.257231 | 0.771693 | ns |
| Fast-Spiking (FS) | Baseline | Layer II/III | 26 | CT vs MP | -1.7677 | -0.847 | 0.397083 | 1 | ns |

**Post-hoc simple-effects contrasts (CT vs MP) — Fast-Spiking (FS), Baseline, Layer IV (Bonferroni correction across 3 ages)**

| Cell Type | Condition | Layer | Age (W) | Comparison | Estimate (Hz) | t_stat | p_raw | p_corrected | sig |
|---|---|---|---|---|---|---|---|---|---|
| Fast-Spiking (FS) | Baseline | Layer IV | 12 | CT vs MP | -3.4764 | -0.984 | 0.32519 | 0.975569 | ns |
| Fast-Spiking (FS) | Baseline | Layer IV | 15 | CT vs MP | 1.9815 | 0.472 | 0.637079 | 1 | ns |

| | | | | | | | | | |
|---|---|---|---|---|---|---|---|---|---|
| Fast-Spiking (FS) | Baseline | Layer IV | 26 | CT vs MP | 3.4594 | 1.231 | 0.218266 | 0.654799 | ns |

**Post-hoc simple-effects contrasts (CT vs MP) — Fast-Spiking (FS), Baseline, Layer V (Bonferroni correction across 3 ages)**

| Cell Type | Condition | Layer | Age (W) | Comparison | Estimate (Hz) | t_stat | p_raw | p_corrected | sig |
|---|---|---|---|---|---|---|---|---|---|
| Fast-Spiking (FS) | Baseline | Layer V | 12 | CT vs MP | -1.4074 | -0.486 | 0.627123 | 1 | ns |
| Fast-Spiking (FS) | Baseline | Layer V | 15 | CT vs MP | 1.6131 | 0.558 | 0.577169 | 1 | ns |
| Fast-Spiking (FS) | Baseline | Layer V | 26 | CT vs MP | -2.3211 | -1.029 | 0.303479 | 0.910437 | ns |

**Post-hoc simple-effects contrasts (CT vs MP) — Fast-Spiking (FS), Baseline, Layer VI (Bonferroni correction across 3 ages)**

| Cell Type | Condition | Layer | Age (W) | Comparison | Estimate (Hz) | t_stat | p_raw | p_corrected | sig |
|---|---|---|---|---|---|---|---|---|---|
| Fast-Spiking (FS) | Baseline | Layer VI | 12 | CT vs MP | 0.1728 | 0.087 | 0.930619 | 1 | ns |
| Fast-Spiking (FS) | Baseline | Layer VI | 15 | CT vs MP | 1.7018 | 0.882 | 0.377768 | 1 | ns |
| Fast-Spiking (FS) | Baseline | Layer VI | 26 | CT vs MP | -2.4607 | -1.581 | 0.113813 | 0.341438 | ns |

**Post-hoc simple-effects contrasts (CT vs MP) — Fast-Spiking (FS), Evoked, Layer II/III (Bonferroni correction across 3 ages)**

| Cell Type | Condition | Layer | Age (W) | Comparison | Estimate (Hz) | t_stat | p_raw | p_corrected | sig |
|---|---|---|---|---|---|---|---|---|---|
| Fast-Spiking (FS) | Evoked | Layer II/III | 12 | CT vs MP | -40.7727 | -1.19 | 0.233918 | 0.701755 | ns |
| Fast-Spiking (FS) | Evoked | Layer II/III | 15 | CT vs MP | 17.5113 | 0.555 | 0.578567 | 1 | ns |
| Fast-Spiking (FS) | Evoked | Layer II/III | 26 | CT vs MP | 13.8294 | 0.662 | 0.50811 | 1 | ns |

**Post-hoc simple-effects contrasts (CT vs MP) — Fast-Spiking (FS), Evoked, Layer IV (Bonferroni correction across 3 ages)**

| Cell Type | Condition | Layer | Age (W) | Comparison | Estimate (Hz) | t_stat | p_raw | p_corrected | sig |
|---|---|---|---|---|---|---|---|---|---|
| Fast-Spiking (FS) | Evoked | Layer IV | 12 | CT vs MP | -23.0222 | -1.19 | 0.234014 | 0.702042 | ns |
| Fast-Spiking (FS) | Evoked | Layer IV | 15 | CT vs MP | 15.637 | 0.738 | 0.460604 | 1 | ns |
| Fast-Spiking (FS) | Evoked | Layer IV | 26 | CT vs MP | 18.4211 | 1.203 | 0.228932 | 0.686797 | ns |

**Post-hoc simple-effects contrasts (CT vs MP) — Fast-Spiking (FS), Evoked, Layer V (Bonferroni correction across 3 ages)**

| Cell Type | Condition | Layer | Age (W) | Comparison | Estimate (Hz) | t_stat | p_raw | p_corrected | sig |
|---|---|---|---|---|---|---|---|---|---|
| Fast-Spiking (FS) | Evoked | Layer V | 12 | CT vs MP | -2.2186 | -0.155 | 0.876645 | 1 | ns |
| Fast-Spiking (FS) | Evoked | Layer V | 15 | CT vs MP | 17.1656 | 1.186 | 0.235766 | 0.707298 | ns |
| Fast-Spiking (FS) | Evoked | Layer V | 26 | CT vs MP | -10.5297 | -0.939 | 0.347601 | 1 | ns |

**Post-hoc simple-effects contrasts (CT vs MP) — Fast-Spiking (FS), Evoked, Layer VI (Bonferroni correction across 3 ages)**

| Cell Type | Condition | Layer | Age (W) | Comparison | Estimate (Hz) | t_stat | p_raw | p_corrected | sig |
|---|---|---|---|---|---|---|---|---|---|
| Fast-Spiking (FS) | Evoked | Layer VI | 12 | CT vs MP | -11.2504 | -0.872 | 0.383335 | 1 | ns |
| Fast-Spiking (FS) | Evoked | Layer VI | 15 | CT vs MP | 14.6149 | 1.145 | 0.252155 | 0.756464 | ns |
| Fast-Spiking (FS) | Evoked | Layer VI | 26 | CT vs MP | -18.5086 | -1.84 | 0.065752 | 0.197257 | ns |

**Figure S2 (Units/session, Fraction of FS, Sparse RS):** LME Models for predicting each session-level metric (genotype × age interaction).

**LME — Units per Session [N sessions = 84, N mice = 45, Log-Likelihood = -365.944, Scale(sigma^2) = 524.42984, Method = REML, Converged = Yes]**

| Term | Beta (Coef.) | SE | z | p | 95% CI Low | 95% CI High | sig |
|---|---|---|---|---|---|---|---|
| Intercept (CT, 12W, reference sex) | 41.867 | 9.583 | 4.369 | 0 | 23.0846 | 60.6494 | |
| Group: MP (vs CT, reference) | 2.0615 | 11.8811 | 0.174 | 0.8623 | -21.225 | 25.3479 | ns |
| Age: 15W (vs 12W, reference) | 3.2681 | 11.7533 | 0.278 | 0.781 | -19.768 | 26.3041 | ns |
| Age: 26W (vs 12W, reference) | 20.2417 | 11.3164 | 1.789 | 0.0737 | -1.938 | 42.4214 | ns |
| Sex: Male (vs Female, reference) | 0.3025 | 8.5044 | 0.036 | 0.9716 | -16.3657 | 16.9708 | ns |
| C(group, Treatment(reference='CT'))[T.MP]:C(age, Treatment(reference='12'))[T.15] | -11.0806 | 16.5834 | -0.668 | 0.504 | -43.5835 | 21.4223 | ns |
| C(group, Treatment(reference='CT'))[T.MP]:C(age, Treatment(reference='12'))[T.26] | -25.971 | 15.0703 | -1.723 | 0.0848 | -55.5083 | 3.5663 | ns |
| Random Effect: Mouse Intercept Variance | 119.456 | 118.5138 | 1.008 | 0.3135 | -112.8267 | 351.7388 | |

**LME — Fraction of FS Units [N sessions = 84, N mice = 45, Log-Likelihood = 63.811, Scale(sigma^2) = 0.00584, Method = REML, Converged = Yes]**

| Term | Beta (Coef.) | SE | z | p | 95% CI Low | 95% CI High | sig |
|---|---|---|---|---|---|---|---|
| Intercept (CT, 12W, reference sex) | 0.3365 | 0.0412 | 8.178 | 0 | 0.2559 | 0.4172 | |
| Group: MP (vs CT, reference) | 0.0139 | 0.0509 | 0.272 | 0.7856 | -0.086 | 0.1137 | ns |
| Age: 15W (vs 12W, reference) | 0.0032 | 0.0503 | 0.063 | 0.9498 | -0.0954 | 0.1018 | ns |
| Age: 26W (vs 12W, reference) | -0.0319 | 0.0484 | -0.66 | 0.5092 | -0.1268 | 0.0629 | ns |
| Sex: Male (vs Female, reference) | -0.018 | 0.0359 | -0.503 | 0.6149 | -0.0883 | 0.0522 | ns |
| C(group, Treatment(reference='CT'))[T.MP]:C(age, Treatment(reference='12'))[T.15] | 0.037 | 0.0719 | 0.515 | 0.6067 | -0.1039 | 0.1779 | ns |
| C(group, Treatment(reference='CT'))[T.MP]:C(age, Treatment(reference='12'))[T.26] | -0.0123 | 0.0645 | -0.191 | 0.8483 | -0.1387 | 0.1141 | ns |
| Random Effect: Mouse Intercept Variance | 0.0042 | 0.0024 | 1.782 | 0.0748 | -0.0004 | 0.0089 | |

**LME — Fraction of Sparse RS Units [N sessions = 84, N mice = 45, Log-Likelihood = 39.481, Scale(sigma^2) = 0.01404, Method = REML, Converged = Yes]**

| Term | Beta (Coef.) | SE | z | p | 95% CI Low | 95% CI High | sig |
|---|---|---|---|---|---|---|---|
| Intercept (CT, 12W, reference sex) | 0.2977 | 0.0495 | 6.011 | 0 | 0.2006 | 0.3947 | |
| Group: MP (vs CT, reference) | 0.0389 | 0.0613 | 0.634 | 0.5261 | -0.0813 | 0.159 | ns |
| Age: 15W (vs 12W, reference) | 0.0178 | 0.0607 | 0.293 | 0.7699 | -0.1013 | 0.1368 | ns |
| Age: 26W (vs 12W, reference) | 0.0394 | 0.0585 | 0.673 | 0.5012 | -0.0754 | 0.1541 | ns |
| Sex: Male (vs Female, reference) | -0.0006 | 0.0439 | -0.014 | 0.9891 | -0.0867 | 0.0855 | ns |
| C(group, Treatment(reference='CT'))[T.MP]:C(age, Treatment(reference='12'))[T.15] | -0.1144 | 0.0856 | -1.336 | 0.1816 | -0.2821 | 0.0534 | ns |
| C(group, Treatment(reference='CT'))[T.MP]:C(age, Treatment(reference='12'))[T.26] | -0.1608 | 0.0777 | -2.069 | **0.0386** | -0.3132 | -0.0085 | * |
| Random Effect: Mouse Intercept Variance | 0.0031 | 0.0035 | 0.897 | 0.3697 | -0.0037 | 0.01 | |

**Figure S2 (Units/session, Fraction of FS, Sparse RS):** model-based simple-effects contrasts of genotype at each age, with Bonferroni correction.

**Post-hoc simple-effects contrasts (CT vs MP) — Units per Session (Bonferroni correction across 3 ages) [omnibus not significant — exploratory]**

| Metric | Age (W) | Comparison | Estimate | t_stat | p_raw | p_corrected | sig |
|---|---|---|---|---|---|---|---|
| Units per Session | 12 | CT vs MP | 2.0615 | 0.174 | 0.862251 | 1 | ns |
| Units per Session | 15 | CT vs MP | -9.0191 | -0.785 | 0.432239 | 1 | ns |
| Units per Session | 26 | CT vs MP | -23.9095 | -2.582 | 0.009833 | **0.0295** | * |

**Post-hoc simple-effects contrasts (CT vs MP) — Fraction of FS Units (Bonferroni correction across 3 ages) [omnibus not significant — exploratory]**

| Metric | Age (W) | Comparison | Estimate | t_stat | p_raw | p_corrected | sig |
|---|---|---|---|---|---|---|---|
| Fraction of FS Units | 12 | CT vs MP | 0.0139 | 0.272 | 0.785623 | 1 | ns |
| Fraction of FS Units | 15 | CT vs MP | 0.0509 | 1.014 | 0.310577 | 0.931732 | ns |
| Fraction of FS Units | 26 | CT vs MP | 0.0015 | 0.039 | 0.96928 | 1 | ns |

**Post-hoc simple-effects contrasts (CT vs MP) — Fraction of Sparse RS Units (Baseline FR < 1 Hz) (Bonferroni correction across 3 ages)**

| Metric | Age (W) | Comparison | Estimate | t_stat | p_raw | p_corrected | sig |
|---|---|---|---|---|---|---|---|
| Fraction of Sparse RS Units (Baseline FR < 1 Hz) | 12 | CT vs MP | 0.0389 | 0.634 | 0.526081 | 1 | ns |
| Fraction of Sparse RS Units (Baseline FR < 1 Hz) | 15 | CT vs MP | -0.0755 | -1.273 | 0.203068 | 0.609203 | ns |
| Fraction of Sparse RS Units (Baseline FR < 1 Hz) | 26 | CT vs MP | -0.1219 | -2.553 | 0.010692 | **0.032076** | * |

**Figure 3 (Fraction of Responsvie Units):** 2-proportion z-test on responsive-unit counts for one cell type, at each age x response window x stimulus intensity.

Bonferroni correction is applied jointly across all 3 ages x 2 periods x 8 intensities (48 comparisons) within each cell type.

**Post-hoc CT vs MP -- Regular-Spiking (RS) (Bonferroni correction across 48 age x period x intensity comparisons)**

| Cell Type | Period | Age (W) | Intensity | Comparison | Estimate (MP-CT) | z_stat | 95% CI Low | 95% CI High | p_raw | p_corrected | sig |
|---|---|---|---|---|---|---|---|---|---|---|---|
| Regular-Spiking (RS) | ONSET | 12 | 1 | CT vs MP | 0.0026 | 0.812 | -0.0025 | 0.0076 | 0.416870938 | 1 | ns |
| Regular-Spiking (RS) | ONSET | 12 | 2 | CT vs MP | -0.0001 | -0.014 | -0.014 | 0.0138 | 0.988648964 | 1 | ns |
| Regular-Spiking (RS) | ONSET | 12 | 5 | CT vs MP | 0.0184 | 0.856 | -0.0228 | 0.0596 | 0.391884509 | 1 | ns |
| Regular-Spiking (RS) | ONSET | 12 | 10 | CT vs MP | -0.0821 | -2.403 | -0.1503 | -0.0138 | 0.016269173 | 0.780920321 | ns |
| Regular-Spiking (RS) | ONSET | 12 | 20 | CT vs MP | -0.1543 | -3.897 | -0.2317 | -0.0769 | 9.74909E-05 | **0.004679562** | * |
| Regular-Spiking (RS) | ONSET | 12 | 40 | CT vs MP | -0.1488 | -3.699 | -0.2268 | -0.0708 | 0.000216184 | **0.010376842** | * |
| Regular-Spiking (RS) | ONSET | 12 | 50 | CT vs MP | -0.1274 | -3.175 | -0.205 | -0.0498 | 0.00149855 | 0.071930403 | ns |
| Regular-Spiking (RS) | ONSET | 12 | 75 | CT vs MP | -0.0884 | -2.271 | -0.1636 | -0.0132 | 0.023131971 | 1 | ns |
| Regular-Spiking (RS) | ONSET | 15 | 1 | CT vs MP | -0.0015 | -0.274 | -0.0117 | 0.0087 | 0.784361773 | 1 | ns |
| Regular-Spiking (RS) | ONSET | 15 | 2 | CT vs MP | 0.0004 | 0.075 | -0.0091 | 0.0099 | 0.940250367 | 1 | ns |
| Regular-Spiking (RS) | ONSET | 15 | 5 | CT vs MP | 0.0008 | 0.042 | -0.0385 | 0.0402 | 0.966421481 | 1 | ns |
| Regular-Spiking (RS) | ONSET | 15 | 10 | CT vs MP | 0.0413 | 1.203 | -0.0271 | 0.1098 | 0.22916316 | 1 | ns |
| Regular-Spiking (RS) | ONSET | 15 | 20 | CT vs MP | 0.0708 | 1.85 | -0.0044 | 0.146 | 0.064378614 | 1 | ns |
| Regular-Spiking (RS) | ONSET | 15 | 40 | CT vs MP | 0.0986 | 2.571 | 0.0245 | 0.1728 | 0.010142367 | 0.4868336 | ns |
| Regular-Spiking (RS) | ONSET | 15 | 50 | CT vs MP | 0.1052 | 2.762 | 0.0321 | 0.1783 | 0.005740656 | 0.275551499 | ns |
| Regular-Spiking (RS) | ONSET | 15 | 75 | CT vs MP | 0.0744 | 2.055 | 0.0053 | 0.1435 | 0.03991856 | 1 | ns |
| Regular-Spiking (RS) | ONSET | 26 | 1 | CT vs MP | 0.0034 | 1.184 | -0.0034 | 0.0102 | 0.236218485 | 1 | ns |
| Regular-Spiking (RS) | ONSET | 26 | 2 | CT vs MP | 0.0309 | 4.419 | 0.0133 | 0.0485 | 9.89514E-06 | **0.000474967** | * |
| Regular-Spiking (RS) | ONSET | 26 | 5 | CT vs MP | 0.1157 | 5.717 | 0.0724 | 0.1589 | 1.08115E-08 | **5.18954E-07** | * |
| Regular-Spiking (RS) | ONSET | 26 | 10 | CT vs MP | 0.1242 | 4.389 | 0.068 | 0.1804 | 1.13666E-05 | **0.000545597** | * |
| Regular-Spiking (RS) | ONSET | 26 | 20 | CT vs MP | 0.1939 | 6.557 | 0.1374 | 0.2503 | 5.479E-11 | **2.63001E-09** | * |
| Regular-Spiking (RS) | ONSET | 26 | 40 | CT vs MP | 0.1959 | 6.691 | 0.141 | 0.2507 | 2.223E-11 | **1.06715E-09** | * |
| Regular-Spiking (RS) | ONSET | 26 | 50 | CT vs MP | 0.19 | 6.557 | 0.1361 | 0.2439 | 5.489E-11 | **2.63496E-09** | * |
| Regular-Spiking (RS) | ONSET | 26 | 75 | CT vs MP | 0.1825 | 6.557 | 0.1318 | 0.2332 | 5.493E-11 | **2.63666E-09** | * |
| Regular-Spiking (RS) | OFFSET | 12 | 1 | CT vs MP | 0 | | 0 | 0 | | | |
| Regular-Spiking (RS) | OFFSET | 12 | 2 | CT vs MP | -0.0013 | -0.298 | -0.0105 | 0.0078 | 0.765336317 | 1 | ns |
| Regular-Spiking (RS) | OFFSET | 12 | 5 | CT vs MP | 0.0115 | 1.381 | -0.0029 | 0.026 | 0.167185643 | 1 | ns |
| Regular-Spiking (RS) | OFFSET | 12 | 10 | CT vs MP | 0.0328 | 1.635 | -0.0045 | 0.0702 | 0.102114648 | 1 | ns |
| Regular-Spiking (RS) | OFFSET | 12 | 20 | CT vs MP | -0.0031 | -0.091 | -0.0706 | 0.0643 | 0.927867652 | 1 | ns |
| Regular-Spiking (RS) | OFFSET | 12 | 40 | CT vs MP | -0.0419 | -1.102 | -0.1169 | 0.0331 | 0.270543559 | 1 | ns |
| Regular-Spiking (RS) | OFFSET | 12 | 50 | CT vs MP | -0.042 | -1.098 | -0.1174 | 0.0333 | 0.27200905 | 1 | ns |
| Regular-Spiking (RS) | OFFSET | 12 | 75 | CT vs MP | -0.0327 | -0.822 | -0.1107 | 0.0453 | 0.411121653 | 1 | ns |
| Regular-Spiking (RS) | OFFSET | 15 | 1 | CT vs MP | 0 | | 0 | 0 | | | |
| Regular-Spiking (RS) | OFFSET | 15 | 2 | CT vs MP | 0.0022 | 0.57 | -0.0066 | 0.011 | 0.568696491 | 1 | ns |
| Regular-Spiking (RS) | OFFSET | 15 | 5 | CT vs MP | 0.0252 | 2.66 | 0.0018 | 0.0486 | 0.007824004 | 0.375552191 | ns |
| Regular-Spiking (RS) | OFFSET | 15 | 10 | CT vs MP | 0.0205 | 0.918 | -0.0248 | 0.0657 | 0.358651906 | 1 | ns |
| Regular-Spiking (RS) | OFFSET | 15 | 20 | CT vs MP | 0.1147 | 3.648 | 0.0495 | 0.18 | 0.000264424 | **0.012692347** | * |
| Regular-Spiking (RS) | OFFSET | 15 | 40 | CT vs MP | 0.1347 | 3.688 | 0.0616 | 0.2078 | 0.000226104 | **0.010852994** | * |
| Regular-Spiking (RS) | OFFSET | 15 | 50 | CT vs MP | 0.1361 | 3.643 | 0.0619 | 0.2102 | 0.000269097 | **0.012916662** | * |
| Regular-Spiking (RS) | OFFSET | 15 | 75 | CT vs MP | 0.1238 | 3.214 | 0.0488 | 0.1987 | 0.001307787 | 0.062773782 | ns |
| Regular-Spiking (RS) | OFFSET | 26 | 1 | CT vs MP | 0.0034 | 1.184 | -0.0034 | 0.0102 | 0.236218485 | 1 | ns |
| Regular-Spiking (RS) | OFFSET | 26 | 2 | CT vs MP | -0.0013 | -0.391 | -0.0074 | 0.0048 | 0.696137628 | 1 | ns |
| Regular-Spiking (RS) | OFFSET | 26 | 5 | CT vs MP | 0.0505 | 4.027 | 0.0225 | 0.0786 | 5.64928E-05 | **0.002711652** | * |
| Regular-Spiking (RS) | OFFSET | 26 | 10 | CT vs MP | 0.1137 | 5.312 | 0.0686 | 0.1588 | 1.08243E-07 | **5.19564E-06** | * |
| Regular-Spiking (RS) | OFFSET | 26 | 20 | CT vs MP | 0.1232 | 4.575 | 0.069 | 0.1774 | 4.75889E-06 | **0.000228427** | * |
| Regular-Spiking (RS) | OFFSET | 26 | 40 | CT vs MP | 0.1373 | 4.794 | 0.0806 | 0.194 | 1.63796E-06 | **7.8622E-05** | * |
| Regular-Spiking (RS) | OFFSET | 26 | 50 | CT vs MP | 0.1423 | 4.914 | 0.0852 | 0.1993 | 8.92606E-07 | **4.28451E-05** | * |
| Regular-Spiking (RS) | OFFSET | 26 | 75 | CT vs MP | 0.1779 | 6.016 | 0.1211 | 0.2347 | 1.7867E-09 | **8.57615E-08** | * |

**Post-hoc CT vs MP -- Fast-Spiking (FS) (Bonferroni correction across 48 age x period x intensity comparisons)**

| Cell Type | Period | Age (W) | Intensity | Comparison | Estimate (MP-CT) | z_stat | 95% CI Low | 95% CI High | p_raw | p_corrected | sig |
|---|---|---|---|---|---|---|---|---|---|---|---|
| Fast-Spiking (FS) | ONSET | 12 | 1 | CT vs MP | 0 | | 0 | 0 | | | |
| Fast-Spiking (FS) | ONSET | 12 | 2 | CT vs MP | -0.0248 | -2.117 | -0.0525 | 0.0029 | 0.034236685 | 1 | ns |
| Fast-Spiking (FS) | ONSET | 12 | 5 | CT vs MP | 0.0128 | 0.33 | -0.0624 | 0.088 | 0.741038867 | 1 | ns |
| Fast-Spiking (FS) | ONSET | 12 | 10 | CT vs MP | -0.0845 | -1.573 | -0.1911 | 0.022 | 0.115832218 | 1 | ns |
| Fast-Spiking (FS) | ONSET | 12 | 20 | CT vs MP | -0.1479 | -2.516 | -0.2614 | -0.0344 | 0.011875395 | 0.570018971 | ns |
| Fast-Spiking (FS) | ONSET | 12 | 40 | CT vs MP | -0.1134 | -1.976 | -0.2239 | -0.0029 | 0.048133566 | 1 | ns |
| Fast-Spiking (FS) | ONSET | 12 | 50 | CT vs MP | -0.113 | -2.004 | -0.2211 | -0.0049 | 0.045066533 | 1 | ns |
| Fast-Spiking (FS) | ONSET | 12 | 75 | CT vs MP | -0.0837 | -1.664 | -0.1793 | 0.012 | 0.096026213 | 1 | ns |
| Fast-Spiking (FS) | ONSET | 15 | 1 | CT vs MP | -0.004 | -0.726 | -0.0118 | 0.0038 | 0.467755564 | 1 | ns |
| Fast-Spiking (FS) | ONSET | 15 | 2 | CT vs MP | -0.008 | -1.028 | -0.019 | 0.003 | 0.303828588 | 1 | ns |
| Fast-Spiking (FS) | ONSET | 15 | 5 | CT vs MP | -0.1163 | -2.899 | -0.1864 | -0.0461 | 0.003749267 | 0.179964829 | ns |
| Fast-Spiking (FS) | ONSET | 15 | 10 | CT vs MP | -0.0857 | -1.596 | -0.1903 | 0.0188 | 0.110549875 | 1 | ns |
| Fast-Spiking (FS) | ONSET | 15 | 20 | CT vs MP | -0.0242 | -0.468 | -0.1259 | 0.0775 | 0.639629079 | 1 | ns |
| Fast-Spiking (FS) | ONSET | 15 | 40 | CT vs MP | -0.0181 | -0.388 | -0.1104 | 0.0741 | 0.69804947 | 1 | ns |

| | | | | | | | | | | | |
|---|---|---|---|---|---|---|---|---|---|---|---|
| Fast-Spiking (FS) | ONSET | 15 | 50 | CT vs MP | 0.003 | 0.068 | -0.0833 | 0.0893 | 0.945438469 | 1 | ns |
| Fast-Spiking (FS) | ONSET | 15 | 75 | CT vs MP | -0.0153 | -0.391 | -0.0928 | 0.0622 | 0.695948643 | 1 | ns |
| Fast-Spiking (FS) | ONSET | 26 | 1 | CT vs MP | -0.0029 | -0.728 | -0.0086 | 0.0028 | 0.466574271 | 1 | ns |
| Fast-Spiking (FS) | ONSET | 26 | 2 | CT vs MP | 0.0178 | 1.098 | -0.0164 | 0.052 | 0.272339781 | 1 | ns |
| Fast-Spiking (FS) | ONSET | 26 | 5 | CT vs MP | 0.0318 | 0.822 | -0.045 | 0.1087 | 0.410834636 | 1 | ns |
| Fast-Spiking (FS) | ONSET | 26 | 10 | CT vs MP | -0.0679 | -1.485 | -0.1569 | 0.0211 | 0.137413346 | 1 | ns |
| Fast-Spiking (FS) | ONSET | 26 | 20 | CT vs MP | -0.0552 | -1.25 | -0.1423 | 0.032 | 0.211229706 | 1 | ns |
| Fast-Spiking (FS) | ONSET | 26 | 40 | CT vs MP | -0.0506 | -1.209 | -0.1337 | 0.0325 | 0.226797967 | 1 | ns |
| Fast-Spiking (FS) | ONSET | 26 | 50 | CT vs MP | -0.0551 | -1.36 | -0.1359 | 0.0257 | 0.173730368 | 1 | ns |
| Fast-Spiking (FS) | ONSET | 26 | 75 | CT vs MP | -0.047 | -1.256 | -0.122 | 0.028 | 0.208963476 | 1 | ns |
| Fast-Spiking (FS) | OFFSET | 12 | 1 | CT vs MP | 0 | | 0 | 0 | | | |
| Fast-Spiking (FS) | OFFSET | 12 | 2 | CT vs MP | 0 | | 0 | 0 | | | |
| Fast-Spiking (FS) | OFFSET | 12 | 5 | CT vs MP | -0.0083 | -1.218 | -0.0244 | 0.0079 | 0.223105068 | 1 | ns |
| Fast-Spiking (FS) | OFFSET | 12 | 10 | CT vs MP | 0.0177 | 0.565 | -0.0424 | 0.0777 | 0.572261372 | 1 | ns |
| Fast-Spiking (FS) | OFFSET | 12 | 20 | CT vs MP | 0.0012 | 0.023 | -0.1034 | 0.1059 | 0.981376711 | 1 | ns |
| Fast-Spiking (FS) | OFFSET | 12 | 40 | CT vs MP | -0.0411 | -0.706 | -0.1555 | 0.0732 | 0.480166382 | 1 | ns |
| Fast-Spiking (FS) | OFFSET | 12 | 50 | CT vs MP | 0.0065 | 0.11 | -0.1086 | 0.1215 | 0.912338485 | 1 | ns |
| Fast-Spiking (FS) | OFFSET | 12 | 75 | CT vs MP | -0.0084 | -0.146 | -0.1218 | 0.1049 | 0.883947799 | 1 | ns |
| Fast-Spiking (FS) | OFFSET | 15 | 1 | CT vs MP | 0 | | 0 | 0 | | | |
| Fast-Spiking (FS) | OFFSET | 15 | 2 | CT vs MP | 0.0076 | 1.381 | -0.0072 | 0.0224 | 0.167353349 | 1 | ns |
| Fast-Spiking (FS) | OFFSET | 15 | 5 | CT vs MP | -0.0171 | -0.879 | -0.0522 | 0.018 | 0.379404342 | 1 | ns |
| Fast-Spiking (FS) | OFFSET | 15 | 10 | CT vs MP | -0.035 | -0.888 | -0.1098 | 0.0399 | 0.374344636 | 1 | ns |
| Fast-Spiking (FS) | OFFSET | 15 | 20 | CT vs MP | 0.0445 | 0.873 | -0.0563 | 0.1453 | 0.382753556 | 1 | ns |
| Fast-Spiking (FS) | OFFSET | 15 | 40 | CT vs MP | 0.1172 | 2.18 | 0.0129 | 0.2215 | 0.029237117 | 1 | ns |
| Fast-Spiking (FS) | OFFSET | 15 | 50 | CT vs MP | 0.0898 | 1.699 | -0.0121 | 0.1916 | 0.089283866 | 1 | ns |
| Fast-Spiking (FS) | OFFSET | 15 | 75 | CT vs MP | 0.0775 | 1.55 | -0.018 | 0.173 | 0.121118902 | 1 | ns |
| Fast-Spiking (FS) | OFFSET | 26 | 1 | CT vs MP | -0.0032 | -0.405 | -0.0178 | 0.0113 | 0.685334179 | 1 | ns |
| Fast-Spiking (FS) | OFFSET | 26 | 2 | CT vs MP | -0.0058 | -1.031 | -0.0138 | 0.0022 | 0.302718905 | 1 | ns |
| Fast-Spiking (FS) | OFFSET | 26 | 5 | CT vs MP | 0.0369 | 1.398 | -0.0175 | 0.0913 | 0.162163144 | 1 | ns |
| Fast-Spiking (FS) | OFFSET | 26 | 10 | CT vs MP | 0.026 | 0.668 | -0.0511 | 0.1031 | 0.503905749 | 1 | ns |
| Fast-Spiking (FS) | OFFSET | 26 | 20 | CT vs MP | -0.0621 | -1.378 | -0.1495 | 0.0253 | 0.168148852 | 1 | ns |
| Fast-Spiking (FS) | OFFSET | 26 | 40 | CT vs MP | -0.1095 | -2.39 | -0.1987 | -0.0203 | 0.016835742 | 0.808115626 | ns |
| Fast-Spiking (FS) | OFFSET | 26 | 50 | CT vs MP | -0.1095 | -2.404 | -0.1988 | -0.0203 | 0.016213103 | 0.778228946 | ns |
| Fast-Spiking (FS) | OFFSET | 26 | 75 | CT vs MP | -0.0629 | -1.463 | -0.1482 | 0.0224 | 0.143519474 | 1 | ns |

**Figure 3 (Fraction of units with well-fitted sensitivity parameter):** 2-proportion z-test on sensitive-unit counts (n_sens / n_total).

Bonferroni correction is applied jointly across all 3 ages x 2 periods (6 comparisons) within each cell type

**Post-hoc CT vs MP -- Regular-Spiking (RS) (Bonferroni correction across 6 age x period comparisons)**

| Cell Type | Period | Age (W) | Compariso | Estimate (MP-CT) | z_stat | 95% CI Low | 95% CI High | p_raw | p_corrected | sig |
|---|---|---|---|---|---|---|---|---|---|---|
| Regular-Spiking (RS) | ONSET | 12 | CT vs MP | -0.093 | -2.38 | -0.17 | -0.0161 | 0.017325 | 0.103948975 | ns |
| Regular-Spiking (RS) | ONSET | 15 | CT vs MP | 0.1115 | 2.897 | 0.0364 | 0.1866 | 0.003763 | **0.022578591** | * |
| Regular-Spiking (RS) | ONSET | 26 | CT vs MP | 0.1735 | 5.868 | 0.1165 | 0.2304 | 4.41E-09 | **2.64591E-08** | * |
| Regular-Spiking (RS) | OFFSET | 12 | CT vs MP | -0.0162 | -0.465 | -0.0846 | 0.0523 | 0.641834 | 1 | ns |
| Regular-Spiking (RS) | OFFSET | 15 | CT vs MP | 0.1175 | 3.466 | 0.0484 | 0.1867 | 0.000528 | **0.003166424** | * |
| Regular-Spiking (RS) | OFFSET | 26 | CT vs MP | 0.1522 | 5.795 | 0.0988 | 0.2055 | 6.82E-09 | **4.09152E-08** | * |

**Post-hoc CT vs MP -- Fast-Spiking (FS) (Bonferroni correction across 6 age x period comparisons)**

| Cell Type | Period | Age (W) | Compariso | Estimate (MP-CT) | z_stat | 95% CI Low | 95% CI High | p_raw | p_corrected | sig |
|---|---|---|---|---|---|---|---|---|---|---|
| Fast-Spiking (FS) | ONSET | 12 | CT vs MP | -0.1204 | -2.046 | -0.2347 | -0.0061 | 0.040714 | 0.244285976 | ns |
| Fast-Spiking (FS) | ONSET | 15 | CT vs MP | 0.0209 | 0.41 | -0.0784 | 0.1201 | 0.682057 | 1 | ns |
| Fast-Spiking (FS) | ONSET | 26 | CT vs MP | -0.0568 | -1.273 | -0.1448 | 0.0312 | 0.203058 | 1 | ns |
| Fast-Spiking (FS) | OFFSET | 12 | CT vs MP | -0.0007 | -0.013 | -0.1105 | 0.109 | 0.989477 | 1 | ns |
| Fast-Spiking (FS) | OFFSET | 15 | CT vs MP | 0.1255 | 2.367 | 0.0213 | 0.2297 | 0.017946 | 0.107677851 | ns |
| Fast-Spiking (FS) | OFFSET | 26 | CT vs MP | -0.0524 | -1.153 | -0.1409 | 0.0361 | 0.24899 | 1 | ns |

**Figure 3 (I0 values):** LME Models to test Age x Group interaction for sensitivity parameters (I0).

**LME -- RS | Period: ONSET [N obs = 1240, N groups = 45, Log-Likelihood = -4912.218, Scale(sigma^2) = 147.60075, Method = REML, Converged = Yes]**

| Term | Beta (Coef.) | SE | z | p | 95% CI Low | 95% CI High | sig |
|---|---|---|---|---|---|---|---|
| Intercept (CT, 12W, reference sex) | 31.5341 | 3.228 | 9.769 | 0 | 25.2074 | 37.8609 | |
| Age: 15W (vs 12W, reference) | 2.2704 | 3.8814 | 0.585 | 0.5586 | -5.337 | 9.8779 | ns |
| Age: 26W (vs 12W, reference) | -1.4591 | 3.7343 | -0.391 | 0.696 | -8.7781 | 5.86 | ns |
| Group: MP (vs CT, reference) | 2.7214 | 3.945 | 0.69 | 0.4903 | -5.0108 | 10.4535 | ns |
| Sex: Male (vs Female, reference) | 0.4634 | 2.7915 | 0.166 | 0.8681 | -5.0078 | 5.9346 | ns |
| Interaction: Age 15W x Group MP | -5.9209 | 5.4843 | -1.08 | 0.2803 | -16.6698 | 4.8281 | ns |
| Interaction: Age 26W x Group MP | -5.896 | 4.9055 | -1.202 | 0.2294 | -15.5105 | 3.7186 | ns |
| Random Effect: Mouse Intercept Variance | 0.0558 | 0.0697 | 0.8 | 0.4234 | -0.0808 | 0.1923 | |
| Random Effect: Session Variance Component (nested within Mouse) | 0.3187 | 0.086 | 3.707 | 0.0002 | 0.1502 | 0.4872 | |

**LME -- RS | Period: OFFSET [N obs = 708, N groups = 44, Log-Likelihood = -2800.377, Scale(sigma^2) = 144.02070, Method = REML, Converged = Yes]**

| Term | Beta (Coef.) | SE | z | p | 95% CI Low | 95% CI High | sig |
|---|---|---|---|---|---|---|---|
| Intercept (CT, 12W, reference sex) | 35.5661 | 3.6723 | 9.685 | 0 | 28.3685 | 42.7637 | |
| Age: 15W (vs 12W, reference) | 1.8585 | 4.3323 | 0.429 | 0.6679 | -6.6326 | 10.3496 | ns |
| Age: 26W (vs 12W, reference) | -1.3631 | 4.2148 | -0.323 | 0.7464 | -9.6239 | 6.8978 | ns |
| Group: MP (vs CT, reference) | 1.0697 | 4.4085 | 0.243 | 0.8083 | -7.5707 | 9.7101 | ns |
| Sex: Male (vs Female, reference) | 2.3482 | 3.051 | 0.77 | 0.4415 | -3.6317 | 8.328 | ns |
| Interaction: Age 15W x Group MP | -4.0571 | 5.9821 | -0.678 | 0.4976 | -15.7817 | 7.6676 | ns |
| Interaction: Age 26W x Group MP | -3.037 | 5.424 | -0.56 | 0.5755 | -13.6679 | 7.5939 | ns |
| Random Effect: Mouse Intercept Variance | 0.0609 | 0.0852 | 0.715 | 0.4746 | -0.1061 | 0.2279 | |
| Random Effect: Session Variance Component (nested within Mouse) | 0.3211 | 0.1058 | 3.035 | 0.0024 | 0.1137 | 0.5285 | |

**LME -- FS | Period: ONSET [N obs = 728, N groups = 45, Log-Likelihood = -2759.934, Scale(sigma^2) = 95.65995, Method = REML, Converged = Yes]**

| Term | Beta (Coef.) | SE | z | p | 95% CI Low | 95% CI High | sig |
|---|---|---|---|---|---|---|---|
| Intercept (CT, 12W, reference sex) | 32.1413 | 3.6564 | 8.79 | 0 | 24.9748 | 39.3077 | |
| Age: 15W (vs 12W, reference) | -3.6834 | 4.3626 | -0.844 | 0.3985 | -12.2339 | 4.8671 | ns |
| Age: 26W (vs 12W, reference) | -4.7463 | 4.2452 | -1.118 | 0.2635 | -13.0667 | 3.5741 | ns |
| Group: MP (vs CT, reference) | 3.0693 | 4.4709 | 0.686 | 0.4924 | -5.6935 | 11.8321 | ns |
| Sex: Male (vs Female, reference) | 0.7425 | 3.1534 | 0.235 | 0.8139 | -5.4382 | 6.9231 | ns |
| Interaction: Age 15W x Group MP | -1.4486 | 6.0927 | -0.238 | 0.8121 | -13.3901 | 10.4929 | ns |
| Interaction: Age 26W x Group MP | -3.5961 | 5.5961 | -0.643 | 0.5205 | -14.5642 | 7.372 | ns |
| Random Effect: Mouse Intercept Variance | 0 | 0.1547 | 0 | 1 | -0.3032 | 0.3032 | |
| Random Effect: Session Variance Component (nested within Mouse) | 0.8783 | 0.2202 | 3.989 | 0.0001 | 0.4467 | 1.3098 | |

**LME -- FS | Period: OFFSET [N obs = 493, N groups = 44, Log-Likelihood = -1920.891, Scale(sigma^2) = 116.21166, Method = REML, Converged = Yes]**

| Term | Beta (Coef.) | SE | z | p | 95% CI Low | 95% CI High | sig |
|---|---|---|---|---|---|---|---|
| Intercept (CT, 12W, reference sex) | 43.0393 | 4.1912 | 10.269 | 0 | 34.8246 | 51.254 | |
| Age: 15W (vs 12W, reference) | -6.7028 | 5.0084 | -1.338 | 0.1808 | -16.519 | 3.1134 | ns |
| Age: 26W (vs 12W, reference) | -9.3245 | 4.853 | -1.921 | 0.0547 | -18.8363 | 0.1872 | ns |
| Group: MP (vs CT, reference) | -4.4472 | 5.0969 | -0.873 | 0.3829 | -14.4369 | 5.5425 | ns |
| Sex: Male (vs Female, reference) | 3.6029 | 3.5491 | 1.015 | 0.31 | -3.3531 | 10.559 | ns |
| Interaction: Age 15W x Group MP | 4.1859 | 6.9114 | 0.606 | 0.5448 | -9.3603 | 17.732 | ns |
| Interaction: Age 26W x Group MP | 2.927 | 6.379 | 0.459 | 0.6463 | -9.5757 | 15.4297 | ns |
| Random Effect: Mouse Intercept Variance | 0 | 0.1817 | 0 | 1 | -0.3562 | 0.3562 | |
| Random Effect: Session Variance Component (nested within Mouse) | 0.8395 | 0.2341 | 3.586 | 0.0003 | 0.3806 | 1.2983 | |

**Figure 3 (I0 Values):** model-based simple effects of genotype (MP - CT) at each age (12/15/26W) with Bonferroni correction.

**Post-hoc simple-effects contrasts (CT vs MP) -- Regular-Spiking (RS), ONSET (Bonferroni correction across 3 ages)**

| Cell Type | Period | Age (W) | Comparison | Estimate (mN) | t_stat | 95% CI Low | 95% CI High | p_raw | p_corrected | sig |
|---|---|---|---|---|---|---|---|---|---|---|
| Regular-Spiking (RS) | ONSET | 12 | CT vs MP | 2.7214 | 0.69 | -5.0108 | 10.4535 | 0.49030617 | 1 | ns |
| Regular-Spiking (RS) | ONSET | 15 | CT vs MP | -3.1995 | -0.857 | -10.5181 | 4.1192 | 0.39153883 | 1 | ns |
| Regular-Spiking (RS) | ONSET | 26 | CT vs MP | -3.1746 | -1.091 | -8.8753 | 2.5261 | 0.27507289 | 0.82521866 | ns |

**Post-hoc simple-effects contrasts (CT vs MP) -- Regular-Spiking (RS), OFFSET (Bonferroni correction across 3 ages)**

| Cell Type | Period | Age (W) | Comparison | Estimate (mN) | t_stat | 95% CI Low | 95% CI High | p_raw | p_corrected | sig |
|---|---|---|---|---|---|---|---|---|---|---|
| Regular-Spiking (RS) | OFFSET | 12 | CT vs MP | 1.0697 | 0.243 | -7.5707 | 9.7101 | 0.80828179 | 1 | ns |
| Regular-Spiking (RS) | OFFSET | 15 | CT vs MP | -2.9874 | -0.747 | -10.8238 | 4.8491 | 0.45495984 | 1 | ns |
| Regular-Spiking (RS) | OFFSET | 26 | CT vs MP | -1.9673 | -0.624 | -8.1453 | 4.2106 | 0.53253075 | 1 | ns |

**Post-hoc simple-effects contrasts (CT vs MP) -- Fast-Spiking (FS), ONSET (Bonferroni correction across 3 ages)**

| Cell Type | Period | Age (W) | Comparison | Estimate (mN) | t_stat | 95% CI Low | 95% CI High | p_raw | p_corrected | sig |
|---|---|---|---|---|---|---|---|---|---|---|
| Fast-Spiking (FS) | ONSET | 12 | CT vs MP | 3.0693 | 0.686 | -5.6935 | 11.8321 | 0.49239963 | 1 | ns |
| Fast-Spiking (FS) | ONSET | 15 | CT vs MP | 1.6207 | 0.396 | -6.4048 | 9.6462 | 0.69225491 | 1 | ns |
| Fast-Spiking (FS) | ONSET | 26 | CT vs MP | -0.5268 | -0.157 | -7.1119 | 6.0582 | 0.87539478 | 1 | ns |

**Post-hoc simple-effects contrasts (CT vs MP) -- Fast-Spiking (FS), OFFSET (Bonferroni correction across 3 ages)**

| Cell Type | Period | Age (W) | Comparison | Estimate (mN) | t_stat | 95% CI Low | 95% CI High | p_raw | p_corrected | sig |
|---|---|---|---|---|---|---|---|---|---|---|
| Fast-Spiking (FS) | OFFSET | 12 | CT vs MP | -4.4472 | -0.873 | -14.4369 | 5.5425 | 0.38291577 | 1 | ns |
| Fast-Spiking (FS) | OFFSET | 15 | CT vs MP | -0.2614 | -0.056 | -9.3461 | 8.8234 | 0.95503474 | 1 | ns |
| Fast-Spiking (FS) | OFFSET | 26 | CT vs MP | -1.5202 | -0.396 | -9.049 | 6.0086 | 0.69228138 | 1 | ns |

**Figure 4 (SNR):** LME models for RS and FS units during ON and OFF periods.

**LME -- RS | Period: ON-period | Layer: Layer II/III [N obs = 225, N groups = 37, Log-Likelihood = -685.159, Scale(sigma^2) = 24.32883, Method = REML, Converged = Yes]**

| Term | Beta (Coef.) | SE | z | p | 95% CI Low | 95% CI High | sig |
|---|---|---|---|---|---|---|---|
| Intercept (CT, 12W, reference sex) | -7.38 | 1.6925 | -4.36 | 0 | -10.6972 | -4.0627 | |
| Age: 15W (vs 12W, reference) | -2.1048 | 2.4837 | -0.847 | 0.3967 | -6.9727 | 2.7631 | ns |
| Age: 26W (vs 12W, reference) | -1.2122 | 1.9319 | -0.627 | 0.5304 | -4.9988 | 2.5743 | ns |
| Group: MP (vs CT, reference) | -2.7737 | 2.3995 | -1.156 | 0.2477 | -7.4767 | 1.9292 | ns |
| Sex: Male (vs Female, reference) | -0.1876 | 2.0675 | -0.091 | 0.9277 | -4.2398 | 3.8646 | ns |
| Interaction: Age 15W x Group MP | 5.3788 | 3.1709 | 1.696 | 0.0898 | -0.836 | 11.5936 | ns |
| Interaction: Age 26W x Group MP | 3.6558 | 2.852 | 1.282 | 0.1999 | -1.9341 | 9.2457 | ns |
| Random Effect: Mouse Intercept Variance | 0.0674 | 0.1395 | 0.483 | 0.629 | -0.206 | 0.3407 | |
| Random Effect: Session Variance Component (nested within Mouse) | 0.2208 | 0.1703 | 1.296 | 0.1948 | -0.113 | 0.5546 | |

**LME -- RS | Period: ON-period | Layer: Layer IV [N obs = 262, N groups = 39, Log-Likelihood = -796.231, Scale(sigma^2) = 23.64347, Method = REML, Converged = Yes]**

| Term | Beta (Coef.) | SE | z | p | 95% CI Low | 95% CI High | sig |
|---|---|---|---|---|---|---|---|
| Intercept (CT, 12W, reference sex) | -7.3309 | 2.3509 | -3.118 | 0.0018 | -11.9387 | -2.7232 | |
| Age: 15W (vs 12W, reference) | -5.7904 | 2.9554 | -1.959 | 0.0501 | -11.5829 | 0.0021 | ns |
| Age: 26W (vs 12W, reference) | -3.666 | 2.502 | -1.465 | 0.1429 | -8.5697 | 1.2377 | ns |
| Group: MP (vs CT, reference) | -4.8134 | 2.7989 | -1.72 | 0.0855 | -10.2992 | 0.6724 | ns |
| Sex: Male (vs Female, reference) | 1.3116 | 1.7748 | 0.739 | 0.4599 | -2.167 | 4.7902 | ns |
| Interaction: Age 15W x Group MP | 7.6631 | 3.5474 | 2.16 | **0.0308** | 0.7102 | 14.616 | * |
| Interaction: Age 26W x Group MP | 8.5408 | 3.165 | 2.699 | **0.007** | 2.3376 | 14.744 | * |
| Random Effect: Mouse Intercept Variance | 0.0229 | 0.2339 | 0.098 | 0.9219 | -0.4356 | 0.4814 | |
| Random Effect: Session Variance Component (nested within Mouse) | 0.2868 | 0.2466 | 1.163 | 0.2448 | -0.1965 | 0.7701 | |

**LME -- RS | Period: ON-period | Layer: Layer V [N obs = 1728, N groups = 45, Log-Likelihood = -5335.037, Scale(sigma^2) = 25.98143, Method = REML, Converged = No]**

| Term | Beta (Coef.) | SE | z | p | 95% CI Low | 95% CI High | sig |
|---|---|---|---|---|---|---|---|
| Intercept (CT, 12W, reference sex) | -9.6564 | 1.1768 | -8.205 | 0 | -11.9629 | -7.3498 | |
| Age: 15W (vs 12W, reference) | 1.6358 | 1.4358 | 1.139 | 0.2546 | -1.1784 | 4.45 | ns |
| Age: 26W (vs 12W, reference) | 0.4907 | 1.3745 | 0.357 | 0.7211 | -2.2033 | 3.1847 | ns |
| Group: MP (vs CT, reference) | 0.6443 | 1.4558 | 0.443 | 0.658 | -2.2089 | 3.4976 | ns |
| Sex: Male (vs Female, reference) | -0.0625 | 1.04 | -0.06 | 0.9521 | -2.1009 | 1.976 | ns |
| Interaction: Age 15W x Group MP | 0.6708 | 2.0227 | 0.332 | 0.7402 | -3.2936 | 4.6352 | ns |
| Interaction: Age 26W x Group MP | 1.751 | 1.8371 | 0.953 | 0.3405 | -1.8497 | 5.3517 | ns |
| Random Effect: Mouse Intercept Variance | 0.0304 | 0.0925 | 0.328 | 0.7429 | -0.151 | 0.2117 | |
| Random Effect: Session Variance Component (nested within Mouse) | 0.2874 | 0.1563 | 1.839 | 0.0659 | -0.0189 | 0.5937 | |

**LME -- RS | Period: ON-period | Layer: Layer VI [N obs = 452, N groups = 40, Log-Likelihood = -1362.518, Scale(sigma^2) = 23.42658, Method = REML, Converged = No]**

| Term | Beta (Coef.) | SE | z | p | 95% CI Low | 95% CI High | sig |
|---|---|---|---|---|---|---|---|
| Intercept (CT, 12W, reference sex) | -11.6416 | 1.0694 | -10.886 | 0 | -13.7376 | -9.5456 | |
| Age: 15W (vs 12W, reference) | -0.8431 | 1.381 | -0.611 | 0.5415 | -3.5498 | 1.8635 | ns |
| Age: 26W (vs 12W, reference) | -0.3727 | 1.2447 | -0.299 | 0.7646 | -2.8122 | 2.0668 | ns |
| Group: MP (vs CT, reference) | -0.3149 | 1.3212 | -0.238 | 0.8116 | -2.9043 | 2.2746 | ns |
| Sex: Male (vs Female, reference) | 0.8542 | 1.1976 | 0.713 | 0.4757 | -1.4931 | 3.2015 | ns |
| Interaction: Age 15W x Group MP | 2.1897 | 1.8522 | 1.182 | 0.2371 | -1.4406 | 5.82 | ns |
| Interaction: Age 26W x Group MP | 1.0362 | 1.6563 | 0.626 | 0.5316 | -2.21 | 4.2825 | ns |
| Random Effect: Mouse Intercept Variance | 0.0481 | 0.0571 | 0.842 | 0.3996 | -0.0638 | 0.1599 | |
| Random Effect: Session Variance Component (nested within Mouse) | 0.0608 | 0.0493 | 1.233 | 0.2176 | -0.0359 | 0.1576 | |

**LME -- RS | Period: OFF-period | Layer: Layer II/III [N obs = 226, N groups = 37, Log-Likelihood = -685.824, Scale(sigma^2) = 25.30992, Method = REML, Converged = No]**

| Term | Beta (Coef.) | SE | z | p | 95% CI Low | 95% CI High | sig |
|---|---|---|---|---|---|---|---|
| Intercept (CT, 12W, reference sex) | -10.016 | 1.4892 | -6.726 | 0 | -12.9347 | -7.0973 | |
| Age: 15W (vs 12W, reference) | -1.0206 | 2.2734 | -0.449 | 0.6535 | -5.4764 | 3.4351 | ns |
| Age: 26W (vs 12W, reference) | -0.0617 | 1.6892 | -0.037 | 0.9708 | -3.3725 | 3.249 | ns |
| Group: MP (vs CT, reference) | -1.2976 | 2.1259 | -0.61 | 0.5416 | -5.4642 | 2.869 | ns |
| Sex: Male (vs Female, reference) | -2.1112 | 1.8736 | -1.127 | 0.2598 | -5.7833 | 1.561 | ns |
| Interaction: Age 15W x Group MP | 3.7948 | 2.817 | 1.347 | 0.178 | -1.7265 | 9.316 | ns |
| Interaction: Age 26W x Group MP | -0.328 | 2.5296 | -0.13 | 0.8968 | -5.2859 | 4.6299 | ns |
| Random Effect: Mouse Intercept Variance | 0.0514 | 0.1985 | 0.259 | 0.7958 | -0.3377 | 0.4405 | |
| Random Effect: Session Variance Component (nested within Mouse) | 0.0928 | 0.1143 | 0.812 | 0.4171 | -0.1313 | 0.3168 | |

**LME -- RS | Period: OFF-period | Layer: Layer IV [N obs = 264, N groups = 39, Log-Likelihood = -773.589, Scale(sigma^2) = 20.75390, Method = REML, Converged = Yes]**

| Term | Beta (Coef.) | SE | z | p | 95% CI Low | 95% CI High | sig |
|---|---|---|---|---|---|---|---|
| Intercept (CT, 12W, reference sex) | -8.8242 | 1.7371 | -5.08 | 0 | -12.2289 | -5.4196 | |
| Age: 15W (vs 12W, reference) | -6.2167 | 2.1371 | -2.909 | **0.0036** | -10.4053 | -2.0282 | * |
| Age: 26W (vs 12W, reference) | -3.4742 | 1.8253 | -1.903 | 0.057 | -7.0517 | 0.1034 | ns |
| Group: MP (vs CT, reference) | -5.3143 | 2.0546 | -2.587 | **0.0097** | -9.3411 | -1.2874 | * |
| Sex: Male (vs Female, reference) | 1.2144 | 1.2717 | 0.955 | 0.3396 | -1.2782 | 3.7069 | ns |
| Interaction: Age 15W x Group MP | 7.4537 | 2.4608 | 3.029 | **0.0025** | 2.6306 | 12.2768 | * |
| Interaction: Age 26W x Group MP | 7.7946 | 2.2973 | 3.393 | **0.0007** | 3.2921 | 12.2972 | * |
| Random Effect: Mouse Intercept Variance | 0.0001 | 0.0742 | 0.002 | 0.9987 | -0.1454 | 0.1456 | |
| Random Effect: Session Variance Component (nested within Mouse) | 0.0819 | 0.0769 | 1.065 | 0.287 | -0.0688 | 0.2326 | |

**LME -- RS | Period: OFF-period | Layer: Layer V [N obs = 1722, N groups = 45, Log-Likelihood = -5261.036, Scale(sigma^2) = 25.00379, Method = REML, Converged = Yes]**

| Term | Beta (Coef.) | SE | z | p | 95% CI Low | 95% CI High | sig |
|---|---|---|---|---|---|---|---|
| Intercept (CT, 12W, reference sex) | -12.3287 | 0.8281 | -14.888 | 0 | -13.9517 | -10.7057 | |
| Age: 15W (vs 12W, reference) | 1.2675 | 1.0027 | 1.264 | 0.2062 | -0.6978 | 3.2329 | ns |
| Age: 26W (vs 12W, reference) | 0.5977 | 0.9573 | 0.624 | 0.5324 | -1.2785 | 2.474 | ns |
| Group: MP (vs CT, reference) | 0.9957 | 1.0088 | 0.987 | 0.3236 | -0.9815 | 2.9728 | ns |
| Sex: Male (vs Female, reference) | -0.1608 | 0.7342 | -0.219 | 0.8266 | -1.5999 | 1.2783 | ns |
| Interaction: Age 15W x Group MP | 0.9589 | 1.41 | 0.68 | 0.4965 | -1.8047 | 3.7225 | ns |
| Interaction: Age 26W x Group MP | 1.3483 | 1.2658 | 1.065 | 0.2868 | -1.1326 | 3.8293 | ns |
| Random Effect: Mouse Intercept Variance | 0 | 0.0341 | 0 | 1 | -0.0668 | 0.0668 | |
| Random Effect: Session Variance Component (nested within Mouse) | 0.143 | 0.0458 | 3.121 | 0.0018 | 0.0532 | 0.2328 | |

**LME -- RS | Period: OFF-period | Layer: Layer VI [N obs = 446, N groups = 40, Log-Likelihood = -1262.999, Scale(sigma^2) = 16.36954, Method = REML, Converged = No]**

| Term | Beta (Coef.) | SE | z | p | 95% CI Low | 95% CI High | sig |
|---|---|---|---|---|---|---|---|
| Intercept (CT, 12W, reference sex) | -14.3611 | 0.8574 | -16.75 | 0 | -16.0416 | -12.6807 | |
| Age: 15W (vs 12W, reference) | -0.6139 | 1.0995 | -0.558 | 0.5766 | -2.769 | 1.5411 | ns |
| Age: 26W (vs 12W, reference) | 0.0019 | 1.0032 | 0.002 | 0.9985 | -1.9643 | 1.9682 | ns |
| Group: MP (vs CT, reference) | -0.1008 | 1.0548 | -0.096 | 0.9239 | -2.1682 | 1.9666 | ns |
| Sex: Male (vs Female, reference) | 0.6414 | 0.9437 | 0.68 | 0.4967 | -1.2082 | 2.491 | ns |
| Interaction: Age 15W x Group MP | 1.7784 | 1.4802 | 1.201 | 0.2296 | -1.1228 | 4.6795 | ns |
| Interaction: Age 26W x Group MP | 0.9143 | 1.3284 | 0.688 | 0.4913 | -1.6894 | 3.518 | ns |
| Random Effect: Mouse Intercept Variance | 0.0433 | 0.0567 | 0.764 | 0.4449 | -0.0678 | 0.1545 | |
| Random Effect: Session Variance Component (nested within Mouse) | 0.0422 | 0.0434 | 0.971 | 0.3316 | -0.043 | 0.1273 | |

**LME -- FS | Period: ON-period | Layer: Layer II/III [N obs = 181, N groups = 38, Log-Likelihood = -531.623, Scale(sigma^2) = 18.80372, Method = REML, Converged = Yes]**

| Term | Beta (Coef.) | SE | z | p | 95% CI Low | 95% CI High | sig |
|---|---|---|---|---|---|---|---|
| Intercept (CT, 12W, reference sex) | -3.9668 | 2.6122 | -1.519 | 0.1289 | -9.0865 | 1.153 | |
| Age: 15W (vs 12W, reference) | -1.6829 | 3.2359 | -0.52 | 0.603 | -8.0251 | 4.6593 | ns |
| Age: 26W (vs 12W, reference) | -1.6455 | 2.8529 | -0.577 | 0.5641 | -7.2371 | 3.9461 | ns |
| Group: MP (vs CT, reference) | -4.4819 | 3.135 | -1.43 | 0.1528 | -10.6263 | 1.6625 | ns |
| Sex: Male (vs Female, reference) | 1.4008 | 1.8691 | 0.749 | 0.4536 | -2.2626 | 5.0642 | ns |
| Interaction: Age 15W x Group MP | 4.9246 | 4.1411 | 1.189 | 0.2344 | -3.1919 | 13.041 | ns |
| Interaction: Age 26W x Group MP | 5.4012 | 3.6595 | 1.476 | 0.14 | -1.7712 | 12.5737 | ns |
| Random Effect: Mouse Intercept Variance | 0.5178 | 0.2341 | 2.212 | 0.027 | 0.059 | 0.9767 | |
| Random Effect: Session Variance Component (nested within Mouse) | 0.0368 | 0.1332 | 0.277 | 0.7822 | -0.2243 | 0.298 | |

**LME -- FS | Period: ON-period | Layer: Layer IV [N obs = 129, N groups = 37, Log-Likelihood = -391.248, Scale(sigma^2) = 24.68488, Method = REML, Converged = Yes]**

| Term | Beta (Coef.) | SE | z | p | 95% CI Low | 95% CI High | sig |
|---|---|---|---|---|---|---|---|
| Intercept (CT, 12W, reference sex) | -3.9111 | 2.0099 | -1.946 | 0.0517 | -7.8504 | 0.0281 | |
| Age: 15W (vs 12W, reference) | -2.6569 | 2.6069 | -1.019 | 0.3081 | -7.7663 | 2.4525 | ns |
| Age: 26W (vs 12W, reference) | -1.4904 | 2.309 | -0.645 | 0.5186 | -6.016 | 3.0351 | ns |
| Group: MP (vs CT, reference) | -3.9934 | 2.475 | -1.613 | 0.1066 | -8.8444 | 0.8576 | ns |
| Sex: Male (vs Female, reference) | -1.6225 | 1.8653 | -0.87 | 0.3844 | -5.2785 | 2.0334 | ns |
| Interaction: Age 15W x Group MP | 5.0453 | 3.7385 | 1.35 | 0.1772 | -2.282 | 12.3726 | ns |
| Interaction: Age 26W x Group MP | 4.3104 | 3.1392 | 1.373 | 0.1697 | -1.8423 | 10.4632 | ns |
| Random Effect: Mouse Intercept Variance | 0.1588 | 0.1936 | 0.82 | 0.412 | -0.2206 | 0.5382 | |
| Random Effect: Session Variance Component (nested within Mouse) | 0.1721 | 0.186 | 0.926 | 0.3546 | -0.1923 | 0.5366 | |

**LME -- FS | Period: ON-period | Layer: Layer V [N obs = 561, N groups = 45, Log-Likelihood = -1757.214, Scale(sigma^2) = 28.83793, Method = REML, Converged = Yes]**

| Term | Beta (Coef.) | SE | z | p | 95% CI Low | 95% CI High | sig |
|---|---|---|---|---|---|---|---|
| Intercept (CT, 12W, reference sex) | -6.2757 | 1.2182 | -5.151 | 0 | -8.6635 | -3.888 | |
| Age: 15W (vs 12W, reference) | -0.4211 | 1.4972 | -0.281 | 0.7785 | -3.3555 | 2.5133 | ns |
| Age: 26W (vs 12W, reference) | 0.212 | 1.4212 | 0.149 | 0.8814 | -2.5735 | 2.9974 | ns |
| Group: MP (vs CT, reference) | -1.0473 | 1.5115 | -0.693 | 0.4884 | -4.0098 | 1.9153 | ns |
| Sex: Male (vs Female, reference) | 0.9607 | 1.1248 | 0.854 | 0.393 | -1.2438 | 3.1653 | ns |
| Interaction: Age 15W x Group MP | 3.1507 | 2.1543 | 1.463 | 0.1436 | -1.0717 | 7.3731 | ns |
| Interaction: Age 26W x Group MP | 0.7978 | 1.9215 | 0.415 | 0.678 | -2.9682 | 4.5638 | ns |
| Random Effect: Mouse Intercept Variance | 0.087 | 0.079 | 1.102 | 0.2704 | -0.0677 | 0.2418 | |
| Random Effect: Session Variance Component (nested within Mouse) | 0.0801 | 0.0727 | 1.102 | 0.2705 | -0.0624 | 0.2226 | |

**LME -- FS | Period: ON-period | Layer: Layer VI [N obs = 335, N groups = 44, Log-Likelihood = -1059.052, Scale(sigma^2) = 29.31518, Method = REML, Converged = Yes]**

| Term | Beta (Coef.) | SE | z | p | 95% CI Low | 95% CI High | sig |
|---|---|---|---|---|---|---|---|
| Intercept (CT, 12W, reference sex) | -11.1601 | 1.7313 | -6.446 | 0 | -14.5533 | -7.7668 | |
| Age: 15W (vs 12W, reference) | 0.2783 | 2.1512 | 0.129 | 0.8971 | -3.938 | 4.4947 | ns |
| Age: 26W (vs 12W, reference) | 2.6948 | 2.0243 | 1.331 | 0.1831 | -1.2728 | 6.6623 | ns |
| Group: MP (vs CT, reference) | -0.4023 | 2.1261 | -0.189 | 0.8499 | -4.5693 | 3.7647 | ns |
| Sex: Male (vs Female, reference) | 2.697 | 1.539 | 1.752 | 0.0797 | -0.3194 | 5.7133 | ns |
| Interaction: Age 15W x Group MP | 0.732 | 3.0313 | 0.241 | 0.8092 | -5.2092 | 6.6732 | ns |
| Interaction: Age 26W x Group MP | -2.0351 | 2.6957 | -0.755 | 0.4503 | -7.3185 | 3.2483 | ns |
| Random Effect: Mouse Intercept Variance | 0.1935 | 0.1049 | 1.844 | 0.0652 | -0.0122 | 0.3992 | |
| Random Effect: Session Variance Component (nested within Mouse) | 0.1339 | 0.0844 | 1.587 | 0.1124 | -0.0314 | 0.2992 | |

**LME -- FS | Period: OFF-period | Layer: Layer II/III [N obs = 181, N groups = 38, Log-Likelihood = -538.508, Scale(sigma^2) = 19.92243, Method = REML, Converged = Yes]**

| Term | Beta (Coef.) | SE | z | p | 95% CI Low | 95% CI High | sig |
|---|---|---|---|---|---|---|---|
| Intercept (CT, 12W, reference sex) | -6.7715 | 2.1953 | -3.085 | 0.002 | -11.0741 | -2.4689 | |
| Age: 15W (vs 12W, reference) | -2.2619 | 2.7678 | -0.817 | 0.4138 | -7.6867 | 3.1629 | ns |
| Age: 26W (vs 12W, reference) | 0.199 | 2.4047 | 0.083 | 0.934 | -4.5141 | 4.9121 | ns |
| Group: MP (vs CT, reference) | -4.6904 | 2.7087 | -1.732 | 0.0833 | -9.9993 | 0.6185 | ns |
| Sex: Male (vs Female, reference) | 2.2226 | 1.6845 | 1.319 | 0.187 | -1.0789 | 5.5241 | ns |
| Interaction: Age 15W x Group MP | 5.4947 | 3.5789 | 1.535 | 0.1247 | -1.5197 | 12.5092 | ns |
| Interaction: Age 26W x Group MP | 3.682 | 3.1995 | 1.151 | 0.2498 | -2.5889 | 9.9529 | ns |
| Random Effect: Mouse Intercept Variance | 0.0671 | 0.1755 | 0.383 | 0.7021 | -0.2768 | 0.4111 | |
| Random Effect: Session Variance Component (nested within Mouse) | 0.3794 | 0.2257 | 1.681 | 0.0927 | -0.0629 | 0.8217 | |

**LME -- FS | Period: OFF-period | Layer: Layer IV [N obs = 129, N groups = 37, Log-Likelihood = -384.715, Scale(sigma^2) = 22.17361, Method = REML, Converged = No]**

| Term | Beta (Coef.) | SE | z | p | 95% CI Low | 95% CI High | sig |
|---|---|---|---|---|---|---|---|
| Intercept (CT, 12W, reference sex) | -6.8243 | 1.8199 | -3.75 | 0.0002 | -10.3913 | -3.2573 | |
| Age: 15W (vs 12W, reference) | -2.9007 | 2.3175 | -1.252 | 0.2107 | -7.443 | 1.6416 | ns |
| Age: 26W (vs 12W, reference) | -0.6201 | 2.076 | -0.299 | 0.7652 | -4.689 | 3.4487 | ns |
| Group: MP (vs CT, reference) | -2.3471 | 2.2611 | -1.038 | 0.2992 | -6.7787 | 2.0845 | ns |
| Sex: Male (vs Female, reference) | -0.0529 | 1.6596 | -0.032 | 0.9745 | -3.3056 | 3.1997 | ns |
| Interaction: Age 15W x Group MP | 3.7014 | 3.2613 | 1.135 | 0.2564 | -2.6906 | 10.0934 | ns |
| Interaction: Age 26W x Group MP | 2.0111 | 2.802 | 0.718 | 0.4729 | -3.4807 | 7.5029 | ns |
| Random Effect: Mouse Intercept Variance | 0.0015 | 0.1781 | 0.009 | 0.9932 | -0.3475 | 0.3505 | |
| Random Effect: Session Variance Component (nested within Mouse) | 0.3015 | 0.2472 | 1.22 | 0.2225 | -0.183 | 0.786 | |

**LME -- FS | Period: OFF-period | Layer: Layer V [N obs = 561, N groups = 45, Log-Likelihood = -1758.147, Scale(sigma^2) = 28.65878, Method = REML, Converged = Yes]**

| Term | Beta (Coef.) | SE | z | p | 95% CI Low | 95% CI High | sig |
|---|---|---|---|---|---|---|---|
| Intercept (CT, 12W, reference sex) | -10.8759 | 1.0997 | -9.89 | 0 | -13.0313 | -8.7205 | |
| Age: 15W (vs 12W, reference) | 0.6146 | 1.3526 | 0.454 | 0.6496 | -2.0364 | 3.2656 | ns |
| Age: 26W (vs 12W, reference) | 1.9957 | 1.2853 | 1.553 | 0.1205 | -0.5233 | 4.5148 | ns |
| Group: MP (vs CT, reference) | 1.8303 | 1.3756 | 1.33 | 0.1834 | -0.8659 | 4.5265 | ns |
| Sex: Male (vs Female, reference) | 1.3904 | 1.0281 | 1.352 | 0.1763 | -0.6246 | 3.4054 | ns |
| Interaction: Age 15W x Group MP | -0.2677 | 1.9147 | -0.14 | 0.8888 | -4.0205 | 3.4851 | ns |
| Interaction: Age 26W x Group MP | -2.5335 | 1.75 | -1.448 | 0.1477 | -5.9634 | 0.8965 | ns |
| Random Effect: Mouse Intercept Variance | 0 | 0.0639 | 0 | 1 | -0.1253 | 0.1253 | |
| Random Effect: Session Variance Component (nested within Mouse) | 0.1712 | 0.0795 | 2.153 | 0.0313 | 0.0154 | 0.327 | |

**LME -- FS | Period: OFF-period | Layer: Layer VI [N obs = 333, N groups = 44, Log-Likelihood = -996.714, Scale(sigma^2) = 20.66076, Method = REML, Converged = Yes]**

| Term | Beta (Coef.) | SE | z | p | 95% CI Low | 95% CI High | sig |
|---|---|---|---|---|---|---|---|
| Intercept (CT, 12W, reference sex) | -13.4775 | 1.3737 | -9.811 | 0 | -16.1699 | -10.7852 | |
| Age: 15W (vs 12W, reference) | -1.1021 | 1.7032 | -0.647 | 0.5176 | -4.4403 | 2.2361 | ns |
| Age: 26W (vs 12W, reference) | 1.7026 | 1.6065 | 1.06 | 0.2892 | -1.446 | 4.8511 | ns |
| Group: MP (vs CT, reference) | -0.538 | 1.6857 | -0.319 | 0.7496 | -3.842 | 2.766 | ns |
| Sex: Male (vs Female, reference) | 3.0265 | 1.2296 | 2.461 | **0.0138** | 0.6166 | 5.4364 | * |
| Interaction: Age 15W x Group MP | 0.9458 | 2.3835 | 0.397 | 0.6915 | -3.7258 | 5.6175 | ns |
| Interaction: Age 26W x Group MP | -1.4147 | 2.1399 | -0.661 | 0.5085 | -5.6087 | 2.7794 | ns |
| Random Effect: Mouse Intercept Variance | 0.1062 | 0.0952 | 1.115 | 0.2648 | -0.0805 | 0.2928 | |
| Random Effect: Session Variance Component (nested within Mouse) | 0.2086 | 0.1141 | 1.828 | 0.0675 | -0.015 | 0.4322 | |

**Figure 4 (SNR):** Post-hoc: model-based simple effects of genotype (MP - CT) at each age (12/15/26W), with Bonferroni correction.

**Post-hoc simple-effects contrasts (CT vs MP) -- Regular-Spiking (RS), ON-period, Layer II/III (Bonferroni correction across 3 ages)**

| Cell Type | Period | Layer | Age (W) | Comparison | Estimate (dB) | t_stat | p_raw | p_corrected | sig |
|---|---|---|---|---|---|---|---|---|---|
| Regular-Spiking (RS) | ON-period | Layer II/III | 12 | CT vs MP | -2.7737 | -1.156 | 0.247695 | 0.743085 | ns |
| Regular-Spiking (RS) | ON-period | Layer II/III | 15 | CT vs MP | 2.6051 | 1.189 | 0.234534 | 0.703602 | ns |
| Regular-Spiking (RS) | ON-period | Layer II/III | 26 | CT vs MP | 0.8821 | 0.577 | 0.563988 | 1 | ns |

**Post-hoc simple-effects contrasts (CT vs MP) -- Regular-Spiking (RS), ON-period, Layer IV (Bonferroni correction across 3 ages)**

| Cell Type | Period | Layer | Age (W) | Comparison | Estimate (dB) | t_stat | p_raw | p_corrected | sig |
|---|---|---|---|---|---|---|---|---|---|
| Regular-Spiking (RS) | ON-period | Layer IV | 12 | CT vs MP | -4.8134 | -1.72 | 0.085484 | 0.256452 | ns |
| Regular-Spiking (RS) | ON-period | Layer IV | 15 | CT vs MP | 2.8497 | 1.231 | 0.218424 | 0.655273 | ns |
| Regular-Spiking (RS) | ON-period | Layer IV | 26 | CT vs MP | 3.7275 | 2.525 | 0.011583 | **0.03475** | * |

**Post-hoc simple-effects contrasts (CT vs MP) -- Regular-Spiking (RS), ON-period, Layer V (Bonferroni correction across 3 ages)**

| Cell Type | Period | Layer | Age (W) | Comparison | Estimate (dB) | t_stat | p_raw | p_corrected | sig |
|---|---|---|---|---|---|---|---|---|---|
| Regular-Spiking (RS) | ON-period | Layer V | 12 | CT vs MP | 0.6443 | 0.443 | 0.658046 | 1 | ns |
| Regular-Spiking (RS) | ON-period | Layer V | 15 | CT vs MP | 1.3151 | 0.943 | 0.34587 | 1 | ns |
| Regular-Spiking (RS) | ON-period | Layer V | 26 | CT vs MP | 2.3953 | 2.168 | 0.030185 | 0.090554 | ns |

**Post-hoc simple-effects contrasts (CT vs MP) -- Regular-Spiking (RS), ON-period, Layer VI (Bonferroni correction across 3 ages)**

| Cell Type | Period | Layer | Age (W) | Comparison | Estimate (dB) | t_stat | p_raw | p_corrected | sig |
|---|---|---|---|---|---|---|---|---|---|
| Regular-Spiking (RS) | ON-period | Layer VI | 12 | CT vs MP | -0.3149 | -0.238 | 0.811633 | 1 | ns |
| Regular-Spiking (RS) | ON-period | Layer VI | 15 | CT vs MP | 1.8748 | 1.462 | 0.14361 | 0.430829 | ns |
| Regular-Spiking (RS) | ON-period | Layer VI | 26 | CT vs MP | 0.7214 | 0.722 | 0.470439 | 1 | ns |

**Post-hoc simple-effects contrasts (CT vs MP) -- Regular-Spiking (RS), OFF-period, Layer II/III (Bonferroni correction across 3 ages)**

| Cell Type | Period | Layer | Age (W) | Comparison | Estimate (dB) | t_stat | p_raw | p_corrected | sig |
|---|---|---|---|---|---|---|---|---|---|
| Regular-Spiking (RS) | OFF-period | Layer II/III | 12 | CT vs MP | -1.2976 | -0.61 | 0.541616 | 1 | ns |
| Regular-Spiking (RS) | OFF-period | Layer II/III | 15 | CT vs MP | 2.4972 | 1.281 | 0.200222 | 0.600666 | ns |
| Regular-Spiking (RS) | OFF-period | Layer II/III | 26 | CT vs MP | -1.6255 | -1.2 | 0.230309 | 0.690927 | ns |

**Post-hoc simple-effects contrasts (CT vs MP) -- Regular-Spiking (RS), OFF-period, Layer IV (Bonferroni correction across 3 ages)**

| Cell Type | Period | Layer | Age (W) | Comparison | Estimate (dB) | t_stat | p_raw | p_corrected | sig |
|---|---|---|---|---|---|---|---|---|---|
| Regular-Spiking (RS) | OFF-period | Layer IV | 12 | CT vs MP | -5.3143 | -2.587 | 0.009693 | **0.029079** | * |
| Regular-Spiking (RS) | OFF-period | Layer IV | 15 | CT vs MP | 2.1394 | 1.466 | 0.142574 | 0.427721 | ns |
| Regular-Spiking (RS) | OFF-period | Layer IV | 26 | CT vs MP | 2.4804 | 2.41 | 0.015972 | **0.047917** | * |

**Post-hoc simple-effects contrasts (CT vs MP) -- Regular-Spiking (RS), OFF-period, Layer V (Bonferroni correction across 3 ages)**

| Cell Type | Period | Layer | Age (W) | Comparison | Estimate (dB) | t_stat | p_raw | p_corrected | sig |
|---|---|---|---|---|---|---|---|---|---|
| Regular-Spiking (RS) | OFF-period | Layer V | 12 | CT vs MP | 0.9957 | 0.987 | 0.323633 | 0.9709 | ns |
| Regular-Spiking (RS) | OFF-period | Layer V | 15 | CT vs MP | 1.9546 | 2.01 | 0.044417 | 0.13325 | ns |
| Regular-Spiking (RS) | OFF-period | Layer V | 26 | CT vs MP | 2.344 | 3.068 | 0.002154 | **0.006463** | * |

**Post-hoc simple-effects contrasts (CT vs MP) -- Regular-Spiking (RS), OFF-period, Layer VI (Bonferroni correction across 3 ages)**

| Cell Type | Period | Layer | Age (W) | Comparison | Estimate (dB) | t_stat | p_raw | p_corrected | sig |
|---|---|---|---|---|---|---|---|---|---|
| Regular-Spiking (RS) | OFF-period | Layer VI | 12 | CT vs MP | -0.1008 | -0.096 | 0.92389 | 1 | ns |
| Regular-Spiking (RS) | OFF-period | Layer VI | 15 | CT vs MP | 1.6776 | 1.639 | 0.101256 | 0.303768 | ns |
| Regular-Spiking (RS) | OFF-period | Layer VI | 26 | CT vs MP | 0.8135 | 1.016 | 0.309602 | 0.928806 | ns |

**Post-hoc simple-effects contrasts (CT vs MP) -- Fast-Spiking (FS), ON-period, Layer II/III (Bonferroni correction across 3 ages)**

| Cell Type | Period | Layer | Age (W) | Comparison | Estimate (dB) | t_stat | p_raw | p_corrected | sig |
|---|---|---|---|---|---|---|---|---|---|
| Fast-Spiking (FS) | ON-period | Layer II/III | 12 | CT vs MP | -4.4819 | -1.43 | 0.152821 | 0.458462 | ns |
| Fast-Spiking (FS) | ON-period | Layer II/III | 15 | CT vs MP | 0.4427 | 0.155 | 0.876578 | 1 | ns |
| Fast-Spiking (FS) | ON-period | Layer II/III | 26 | CT vs MP | 0.9194 | 0.479 | 0.632026 | 1 | ns |

**Post-hoc simple-effects contrasts (CT vs MP) -- Fast-Spiking (FS), ON-period, Layer IV (Bonferroni correction across 3 ages)**

| Cell Type | Period | Layer | Age (W) | Comparison | Estimate (dB) | t_stat | p_raw | p_corrected | sig |
|---|---|---|---|---|---|---|---|---|---|
| Fast-Spiking (FS) | ON-period | Layer IV | 12 | CT vs MP | -3.9934 | -1.613 | 0.106643 | 0.31993 | ns |
| Fast-Spiking (FS) | ON-period | Layer IV | 15 | CT vs MP | 1.0519 | 0.374 | 0.708688 | 1 | ns |
| Fast-Spiking (FS) | ON-period | Layer IV | 26 | CT vs MP | 0.3171 | 0.163 | 0.870855 | 1 | ns |

**Post-hoc simple-effects contrasts (CT vs MP) -- Fast-Spiking (FS), ON-period, Layer V (Bonferroni correction across 3 ages)**

| Cell Type | Period | Layer | Age (W) | Comparison | Estimate (dB) | t_stat | p_raw | p_corrected | sig |
|---|---|---|---|---|---|---|---|---|---|
| Fast-Spiking (FS) | ON-period | Layer V | 12 | CT vs MP | -1.0473 | -0.693 | 0.488414 | 1 | ns |
| Fast-Spiking (FS) | ON-period | Layer V | 15 | CT vs MP | 2.1035 | 1.372 | 0.170078 | 0.510235 | ns |
| Fast-Spiking (FS) | ON-period | Layer V | 26 | CT vs MP | -0.2494 | -0.21 | 0.833405 | 1 | ns |

**Post-hoc simple-effects contrasts (CT vs MP) -- Fast-Spiking (FS), ON-period, Layer VI (Bonferroni correction across 3 ages)**

| Cell Type | Period | Layer | Age (W) | Comparison | Estimate (dB) | t_stat | p_raw | p_corrected | sig |
|---|---|---|---|---|---|---|---|---|---|
| Fast-Spiking (FS) | ON-period | Layer VI | 12 | CT vs MP | -0.4023 | -0.189 | 0.849934 | 1 | ns |
| Fast-Spiking (FS) | ON-period | Layer VI | 15 | CT vs MP | 0.3298 | 0.154 | 0.877501 | 1 | ns |
| Fast-Spiking (FS) | ON-period | Layer VI | 26 | CT vs MP | -2.4374 | -1.47 | 0.141521 | 0.424564 | ns |

**Post-hoc simple-effects contrasts (CT vs MP) -- Fast-Spiking (FS), OFF-period, Layer II/III (Bonferroni correction across 3 ages)**

| Cell Type | Period | Layer | Age (W) | Comparison | Estimate (dB) | t_stat | p_raw | p_corrected | sig |
|---|---|---|---|---|---|---|---|---|---|
| Fast-Spiking (FS) | OFF-period | Layer II/III | 12 | CT vs MP | -4.6904 | -1.732 | 0.083342 | 0.250026 | ns |
| Fast-Spiking (FS) | OFF-period | Layer II/III | 15 | CT vs MP | 0.8043 | 0.337 | 0.736135 | 1 | ns |
| Fast-Spiking (FS) | OFF-period | Layer II/III | 26 | CT vs MP | -1.0084 | -0.593 | 0.553037 | 1 | ns |

**Post-hoc simple-effects contrasts (CT vs MP) -- Fast-Spiking (FS), OFF-period, Layer IV (Bonferroni correction across 3 ages)**

| Cell Type | Period | Layer | Age (W) | Comparison | Estimate (dB) | t_stat | p_raw | p_corrected | sig |
|---|---|---|---|---|---|---|---|---|---|
| Fast-Spiking (FS) | OFF-period | Layer IV | 12 | CT vs MP | -2.3471 | -1.038 | 0.299248 | 0.897745 | ns |
| Fast-Spiking (FS) | OFF-period | Layer IV | 15 | CT vs MP | 1.3543 | 0.564 | 0.572736 | 1 | ns |
| Fast-Spiking (FS) | OFF-period | Layer IV | 26 | CT vs MP | -0.336 | -0.193 | 0.846603 | 1 | ns |

**Post-hoc simple-effects contrasts (CT vs MP) -- Fast-Spiking (FS), OFF-period, Layer V (Bonferroni correction across 3 ages)**

| Cell Type | Period | Layer | Age (W) | Comparison | Estimate (dB) | t_stat | p_raw | p_corrected | sig |
|---|---|---|---|---|---|---|---|---|---|
| Fast-Spiking (FS) | OFF-period | Layer V | 12 | CT vs MP | 1.8303 | 1.33 | 0.183358 | 0.550073 | ns |
| Fast-Spiking (FS) | OFF-period | Layer V | 15 | CT vs MP | 1.5626 | 1.181 | 0.237472 | 0.712416 | ns |
| Fast-Spiking (FS) | OFF-period | Layer V | 26 | CT vs MP | -0.7032 | -0.65 | 0.515768 | 1 | ns |

**Post-hoc simple-effects contrasts (CT vs MP) -- Fast-Spiking (FS), OFF-period, Layer VI (Bonferroni correction across 3 ages)**

| Cell Type | Period | Layer | Age (W) | Comparison | Estimate (dB) | t_stat | p_raw | p_corrected | sig |
|---|---|---|---|---|---|---|---|---|---|
| Fast-Spiking (FS) | OFF-period | Layer VI | 12 | CT vs MP | -0.538 | -0.319 | 0.749597 | 1 | ns |
| Fast-Spiking (FS) | OFF-period | Layer VI | 15 | CT vs MP | 0.4078 | 0.244 | 0.807238 | 1 | ns |
| Fast-Spiking (FS) | OFF-period | Layer VI | 26 | CT vs MP | -1.9527 | -1.482 | 0.138368 | 0.415103 | ns |

**Figure 5 (Signal Corr.):** LME models for all units, RS units only, RS-FS units, and FS units only.

**LME — All [N sessions = 84, N mice = 45, Log-Likelihood = 51.091, Scale(sigma^2) = 0.00451, Method = REML, Converged = Yes]**

| Term | Beta (Coef.) | SE | z | p | 95% CI Low | 95% CI High | sig |
|---|---|---|---|---|---|---|---|
| Intercept (CT, 12W, reference sex) | 0.2391 | 0.0443 | 5.394 | 0 | 0.1522 | 0.326 | |
| Group: MP (vs CT, reference) | -0.0232 | 0.055 | -0.422 | 0.6732 | -0.1309 | 0.0845 | ns |
| Age: 15W (vs 12W, reference) | 0.0453 | 0.0543 | 0.835 | 0.4037 | -0.0611 | 0.1518 | ns |
| Age: 26W (vs 12W, reference) | 0.0071 | 0.0523 | 0.135 | 0.8924 | -0.0954 | 0.1096 | ns |
| Sex: Male (vs Female, reference) | 0.0139 | 0.039 | 0.356 | 0.7219 | -0.0626 | 0.0903 | ns |
| Interaction: Age 15W x Group MP | 0.0766 | 0.077 | 0.995 | 0.3196 | -0.0743 | 0.2276 | ns |
| Interaction: Age 26W x Group MP | 0.1112 | 0.0696 | 1.596 | 0.1104 | -0.0253 | 0.2476 | ns |
| Random Effect: Mouse Intercept Variance | 0.0034 | | | | | | |
| Random Effect: Session Variance Component (nested within Mouse) | 0.0051 | | | | | | |

**LME — RS [N sessions = 84, N mice = 45, Log-Likelihood = 37.781, Scale(sigma^2) = 0.00626, Method = REML, Converged = Yes]**

| Term | Beta (Coef.) | SE | z | p | 95% CI Low | 95% CI High | sig |
|---|---|---|---|---|---|---|---|
| Intercept (CT, 12W, reference sex) | 0.2099 | 0.0472 | 4.448 | 0 | 0.1174 | 0.3024 | |
| Group: MP (vs CT, reference) | 0.01 | 0.0586 | 0.17 | 0.8649 | -0.1048 | 0.1248 | ns |
| Age: 15W (vs 12W, reference) | 0.0444 | 0.0579 | 0.768 | 0.4426 | -0.069 | 0.1579 | ns |
| Age: 26W (vs 12W, reference) | 0.0052 | 0.0559 | 0.093 | 0.926 | -0.1044 | 0.1147 | ns |
| Sex: Male (vs Female, reference) | -0.0005 | 0.0422 | -0.012 | 0.99 | -0.0832 | 0.0822 | ns |
| Interaction: Age 15W x Group MP | 0.0642 | 0.081 | 0.792 | 0.4281 | -0.0946 | 0.223 | ns |
| Interaction: Age 26W x Group MP | 0.1191 | 0.0743 | 1.603 | 0.109 | -0.0266 | 0.2649 | ns |
| Random Effect: Mouse Intercept Variance | 0.0008 | | | | | | |
| Random Effect: Session Variance Component (nested within Mouse) | 0.0104 | | | | | | |

**LME — FS [N sessions = 83, N mice = 45, Log-Likelihood = 33.642, Scale(sigma^2) = 0.00673, Method = REML, Converged = Yes]**

| Term | Beta (Coef.) | SE | z | p | 95% CI Low | 95% CI High | sig |
|---|---|---|---|---|---|---|---|
| Intercept (CT, 12W, reference sex) | 0.319 | 0.0632 | 5.045 | 0 | 0.195 | 0.4429 | |
| Group: MP (vs CT, reference) | -0.0728 | 0.0783 | -0.929 | 0.3528 | -0.2262 | 0.0807 | ns |
| Age: 15W (vs 12W, reference) | 0.0421 | 0.0776 | 0.542 | 0.5878 | -0.11 | 0.1941 | ns |
| Age: 26W (vs 12W, reference) | 0.0475 | 0.0743 | 0.64 | 0.522 | -0.098 | 0.1931 | ns |
| Sex: Male (vs Female, reference) | 0.0463 | 0.0549 | 0.844 | 0.3984 | -0.0612 | 0.1539 | ns |
| Interaction: Age 15W x Group MP | 0.0968 | 0.1112 | 0.871 | 0.384 | -0.1211 | 0.3147 | ns |
| Interaction: Age 26W x Group MP | 0.048 | 0.099 | 0.485 | 0.6279 | -0.1461 | 0.2421 | ns |
| Random Effect: Mouse Intercept Variance | 0.0113 | | | | | | |
| Random Effect: Session Variance Component (nested within Mouse) | 0.0048 | | | | | | |

**LME — RS-FS [N sessions = 84, N mice = 45, Log-Likelihood = 53.039, Scale(sigma^2) = 0.00428, Method = REML, Converged = Yes]**

| Term | Beta (Coef.) | SE | z | p | 95% CI Low | 95% CI High | sig |
|---|---|---|---|---|---|---|---|
| Intercept (CT, 12W, reference sex) | 0.2588 | 0.0436 | 5.933 | 0 | 0.1733 | 0.3443 | |
| Group: MP (vs CT, reference) | -0.0455 | 0.0541 | -0.841 | 0.4003 | -0.1515 | 0.0605 | ns |
| Age: 15W (vs 12W, reference) | 0.0326 | 0.0534 | 0.609 | 0.5424 | -0.0722 | 0.1373 | ns |
| Age: 26W (vs 12W, reference) | 0.0054 | 0.0514 | 0.105 | 0.9165 | -0.0954 | 0.1062 | ns |
| Sex: Male (vs Female, reference) | 0.0264 | 0.0383 | 0.688 | 0.4915 | -0.0488 | 0.1015 | ns |
| Interaction: Age 15W x Group MP | 0.0972 | 0.0759 | 1.282 | 0.2 | -0.0515 | 0.2459 | ns |
| Interaction: Age 26W x Group MP | 0.1174 | 0.0685 | 1.714 | 0.0866 | -0.0169 | 0.2518 | ns |
| Random Effect: Mouse Intercept Variance | 0.0035 | | | | | | |
| Random Effect: Session Variance Component (nested within Mouse) | 0.0047 | | | | | | |

**Figure 5 (Signal Corr):** Model-based simple effect contrasts with Bonferroni correction.

**Post-hoc simple-effects contrasts (CT vs MP) — All (Bonferroni correction across 3 ages)**

| Populatio | Age (W) | Comparison | Estimate | t_stat | p_raw | p_corrected | sig |
|---|---|---|---|---|---|---|---|
| All | 12 | CT vs MP | -0.0232 | -0.422 | 0.673202 | 1 | ns |
| All | 15 | CT vs MP | 0.0535 | 1 | 0.317525 | 0.952575 | ns |
| All | 26 | CT vs MP | 0.088 | 2.058 | 0.039566 | 0.118697 | ns |

**Post-hoc simple-effects contrasts (CT vs MP) — RS (Bonferroni correction across 3 ages)**

| Populatio | Age (W) | Comparison | Estimate | t_stat | p_raw | p_corrected | sig |
|---|---|---|---|---|---|---|---|
| RS | 12 | CT vs MP | 0.01 | 0.17 | 0.864864 | 1 | ns |
| RS | 15 | CT vs MP | 0.0742 | 1.332 | 0.18287 | 0.548609 | ns |
| RS | 26 | CT vs MP | 0.1291 | 2.82 | 0.004796 | **0.014388** | * |

**Post-hoc simple-effects contrasts (CT vs MP) — FS (Bonferroni correction across 3 ages)**

| Populatio | Age (W) | Comparison | Estimate | t_stat | p_raw | p_corrected | sig |
|---|---|---|---|---|---|---|---|
| FS | 12 | CT vs MP | -0.0728 | -0.929 | 0.352813 | 1 | ns |
| FS | 15 | CT vs MP | 0.024 | 0.308 | 0.757835 | 1 | ns |
| FS | 26 | CT vs MP | -0.0247 | -0.408 | 0.682974 | 1 | ns |

**Post-hoc simple-effects contrasts (CT vs MP) — RS-FS (Bonferroni correction across 3 ages)**

| Populatio | Age (W) | Comparison | Estimate | t_stat | p_raw | p_corrected | sig |
|---|---|---|---|---|---|---|---|
| RS-FS | 12 | CT vs MP | -0.0455 | -0.841 | 0.400277 | 1 | ns |
| RS-FS | 15 | CT vs MP | 0.0517 | 0.981 | 0.326511 | 0.979533 | ns |
| RS-FS | 26 | CT vs MP | 0.072 | 1.711 | 0.087086 | 0.261259 | ns |

**Figure 5 (Noise Corr):** LME Models for all units, RS-RS units only, RS-FS units, and FS-FS units only.

**LME — All [N sessions = 84, N mice = 45, Log-Likelihood = 56.753, Scale(sigma^2) = 0.00390, Method = REML, Converged = Yes]**

| Term | Beta (Coef.) | SE | z | p | 95% CI Low | 95% CI High | sig |
|---|---|---|---|---|---|---|---|
| Intercept (CT, 12W, reference sex) | 0.1969 | 0.0377 | 5.23 | 0 | 0.1231 | 0.2707 | |
| Group: MP (vs CT, reference) | 0.0107 | 0.0467 | 0.23 | 0.8184 | -0.0808 | 0.1023 | ns |
| Age: 15W (vs 12W, reference) | 0.011 | 0.0462 | 0.238 | 0.8118 | -0.0795 | 0.1015 | ns |
| Age: 26W (vs 12W, reference) | -0.0334 | 0.0446 | -0.75 | 0.4533 | -0.1207 | 0.0539 | ns |
| Sex: Male (vs Female, reference) | 0.0305 | 0.0336 | 0.909 | 0.3632 | -0.0353 | 0.0963 | ns |
| Interaction: Age 15W x Group MP | -0.017 | 0.0648 | -0.262 | 0.7931 | -0.1441 | 0.1101 | ns |
| Interaction: Age 26W x Group MP | 0.0086 | 0.0593 | 0.145 | 0.8845 | -0.1076 | 0.1248 | ns |
| Random Effect: Mouse Intercept Variance | 0.001 | | | | | | |
| Random Effect: Session Variance Component (nested within Mouse) | 0.0059 | | | | | | |

**LME — RS [N sessions = 84, N mice = 45, Log-Likelihood = 50.206, Scale(sigma^2) = 0.00458, Method = REML, Converged = Yes]**

| Term | Beta (Coef.) | SE | z | p | 95% CI Low | 95% CI High | sig |
|---|---|---|---|---|---|---|---|
| Intercept (CT, 12W, reference sex) | 0.1836 | 0.0404 | 4.546 | 0 | 0.1044 | 0.2628 | |
| Group: MP (vs CT, reference) | 0.0227 | 0.0501 | 0.454 | 0.65 | -0.0755 | 0.121 | ns |
| Age: 15W (vs 12W, reference) | -0.0097 | 0.0495 | -0.195 | 0.845 | -0.1068 | 0.0874 | ns |
| Age: 26W (vs 12W, reference) | -0.0496 | 0.0478 | -1.038 | 0.2993 | -0.1434 | 0.0441 | ns |
| Sex: Male (vs Female, reference) | 0.0153 | 0.0361 | 0.425 | 0.6707 | -0.0554 | 0.086 | ns |
| Interaction: Age 15W x Group MP | -0.0105 | 0.0694 | -0.152 | 0.8793 | -0.1466 | 0.1255 | ns |
| Interaction: Age 26W x Group MP | 0.0226 | 0.0636 | 0.356 | 0.7222 | -0.1021 | 0.1473 | ns |
| Random Effect: Mouse Intercept Variance | 0.0008 | | | | | | |
| Random Effect: Session Variance Component (nested within Mouse) | 0.0073 | | | | | | |

**LME — FS [N sessions = 83, N mice = 45, Log-Likelihood = 40.958, Scale(sigma^2) = 0.00578, Method = REML, Converged = Yes]**

| Term | Beta (Coef.) | SE | z | p | 95% CI Low | 95% CI High | sig |
|---|---|---|---|---|---|---|---|
| Intercept (CT, 12W, reference sex) | 0.2645 | 0.0498 | 5.312 | 0 | 0.1669 | 0.3621 | |
| Group: MP (vs CT, reference) | -0.0326 | 0.0617 | -0.528 | 0.5976 | -0.1536 | 0.0884 | ns |
| Age: 15W (vs 12W, reference) | 0.0132 | 0.0614 | 0.216 | 0.8292 | -0.107 | 0.1335 | ns |
| Age: 26W (vs 12W, reference) | 0.0071 | 0.0587 | 0.121 | 0.9036 | -0.108 | 0.1223 | ns |
| Sex: Male (vs Female, reference) | 0.0567 | 0.044 | 1.288 | 0.1976 | -0.0296 | 0.143 | ns |
| Interaction: Age 15W x Group MP | 0.0308 | 0.0869 | 0.354 | 0.7233 | -0.1396 | 0.2012 | ns |
| Interaction: Age 26W x Group MP | -0.0123 | 0.0782 | -0.157 | 0.875 | -0.1656 | 0.141 | ns |
| Random Effect: Mouse Intercept Variance | 0.0041 | | | | | | |
| Random Effect: Session Variance Component (nested within Mouse) | 0.0067 | | | | | | |

**LME — RS-FS [N sessions = 84, N mice = 45, Log-Likelihood = 58.363, Scale(sigma^2) = 0.00353, Method = REML, Converged = Yes]**

| Term | Beta (Coef.) | SE | z | p | 95% CI Low | 95% CI High | sig |
|---|---|---|---|---|---|---|---|
| Intercept (CT, 12W, reference sex) | 0.2101 | 0.0354 | 5.935 | 0 | 0.1407 | 0.2795 | |
| Group: MP (vs CT, reference) | -0.0064 | 0.044 | -0.145 | 0.8851 | -0.0925 | 0.0798 | ns |
| Age: 15W (vs 12W, reference) | 0.0054 | 0.0434 | 0.125 | 0.9005 | -0.0797 | 0.0905 | ns |
| Age: 26W (vs 12W, reference) | -0.0328 | 0.042 | -0.782 | 0.434 | -0.1151 | 0.0494 | ns |
| Sex: Male (vs Female, reference) | 0.0376 | 0.0317 | 1.183 | 0.2367 | -0.0247 | 0.0998 | ns |
| Interaction: Age 15W x Group MP | -0.0015 | 0.0606 | -0.024 | 0.9806 | -0.1202 | 0.1173 | ns |
| Interaction: Age 26W x Group MP | 0.0165 | 0.0558 | 0.295 | 0.768 | -0.0929 | 0.1259 | ns |
| Random Effect: Mouse Intercept Variance | 0.0001 | | | | | | |
| Random Effect: Session Variance Component (nested within Mouse) | 0.0066 | | | | | | |

**Figure 5 (Noise Corr):** Model-based simple effect contrasts, with Bonferroni correction.

**Post-hoc simple-effects contrasts (CT vs MP) — All (Bonferroni correction across 3 ages)**

| Population | Age (W) | Comparison | Estimate | t_stat | p_raw | p_corrected | sig |
|---|---|---|---|---|---|---|---|
| All | 12 | CT vs MP | 0.0107 | 0.23 | 0.818399 | 1 | ns |
| All | 15 | CT vs MP | -0.0063 | -0.14 | 0.888337 | 1 | ns |
| All | 26 | CT vs MP | 0.0193 | 0.53 | 0.595966 | 1 | ns |

**Post-hoc simple-effects contrasts (CT vs MP) — RS (Bonferroni correction across 3 ages)**

| Population | Age (W) | Comparison | Estimate | t_stat | p_raw | p_corrected | sig |
|---|---|---|---|---|---|---|---|
| RS | 12 | CT vs MP | 0.0227 | 0.454 | 0.649986 | 1 | ns |
| RS | 15 | CT vs MP | 0.0122 | 0.256 | 0.798151 | 1 | ns |
| RS | 26 | CT vs MP | 0.0454 | 1.158 | 0.246713 | 0.740139 | ns |

**Post-hoc simple-effects contrasts (CT vs MP) — FS (Bonferroni correction across 3 ages)**

| Population | Age (W) | Comparison | Estimate | t_stat | p_raw | p_corrected | sig |
|---|---|---|---|---|---|---|---|
| FS | 12 | CT vs MP | -0.0326 | -0.528 | 0.597587 | 1 | ns |
| FS | 15 | CT vs MP | -0.0018 | -0.03 | 0.976375 | 1 | ns |
| FS | 26 | CT vs MP | -0.0449 | -0.935 | 0.349893 | 1 | ns |

**Post-hoc simple-effects contrasts (CT vs MP) — RS-FS (Bonferroni correction across 3 ages)**

| Population | Age (W) | Comparison | Estimate | t_stat | p_raw | p_corrected | sig |
|---|---|---|---|---|---|---|---|
| RS-FS | 12 | CT vs MP | -0.0064 | -0.145 | 0.885083 | 1 | ns |
| RS-FS | 15 | CT vs MP | -0.0078 | -0.188 | 0.850511 | 1 | ns |
| RS-FS | 26 | CT vs MP | 0.0101 | 0.294 | 0.768687 | 1 | ns |

**Figure 6 (Population Coupling):** LME models for all units, RS units only, RS-FS units, and FS units only.

**LME — All [N units = 3906, N mice = 45, Log-Likelihood = -25691.395, Scale(sigma^2) = 28848.28769, Method = REML, Converged = Yes]**

| Term | Beta (Coef.) | SE | z | p | 95% CI Low | 95% CI High | sig |
|---|---|---|---|---|---|---|---|
| Intercept (CT, 12W, reference sex) | 219.2089 | 51.5068 | 4.256 | 0 | 118.2574 | 320.1603 | |
| Age: 15W (vs 12W, reference) | -5.4985 | 63.2457 | -0.087 | 0.9307 | -129.4578 | 118.4609 | ns |
| Age: 26W (vs 12W, reference) | 37.036 | 60.4783 | 0.612 | 0.5403 | -81.4993 | 155.5712 | ns |
| Group: MP (vs CT, reference) | -40.8191 | 63.6508 | -0.641 | 0.5213 | -165.5723 | 83.9341 | ns |
| Sex: Male (vs Female, reference) | 21.0058 | 45.2437 | 0.464 | 0.6424 | -67.6703 | 109.6818 | ns |
| Interaction: Age 15W x Group MP | 41.4988 | 89.9117 | 0.462 | 0.6444 | -134.7248 | 217.7225 | ns |
| Interaction: Age 26W x Group MP | -29.3088 | 80.4953 | -0.364 | 0.7158 | -187.0766 | 128.459 | ns |
| Random Effect: Mouse Intercept Variance | 5597.751 | 2623.769 | 2.133 | 0.0329 | 455.259 | 10740.2431 | |
| Random Effect: Session Variance Component (nested within Mouse) | 10082.9848 | 2295.147 | 4.393 | 0 | 5584.58 | 14581.3895 | |

**LME — RS [N units = 2697, N mice = 45, Log-Likelihood = -16003.207, Scale(sigma^2) = 7813.86159, Method = REML, Converged = Yes]**

| Term | Beta (Coef.) | SE | z | p | 95% CI Low | 95% CI High | sig |
|---|---|---|---|---|---|---|---|
| Intercept (CT, 12W, reference sex) | 115.5084 | 30.9811 | 3.728 | 0.0002 | 54.7866 | 176.2302 | |
| Age: 15W (vs 12W, reference) | -6.986 | 37.9954 | -0.184 | 0.8541 | -81.4556 | 67.4835 | ns |
| Age: 26W (vs 12W, reference) | 9.6459 | 36.3211 | 0.266 | 0.7906 | -61.5423 | 80.834 | ns |
| Group: MP (vs CT, reference) | -17.2371 | 38.2782 | -0.45 | 0.6525 | -92.2611 | 57.7868 | ns |
| Sex: Male (vs Female, reference) | 3.2709 | 27.1824 | 0.12 | 0.9042 | -50.0056 | 56.5474 | ns |
| Interaction: Age 15W x Group MP | 17.7089 | 54.2507 | 0.326 | 0.7441 | -88.6204 | 124.0383 | ns |
| Interaction: Age 26W x Group MP | 16.7656 | 48.3458 | 0.347 | 0.7288 | -77.9904 | 111.5217 | ns |
| Random Effect: Mouse Intercept Variance | 2214.2094 | 922.0299 | 2.401 | 0.0163 | 407.0639 | 4021.3548 | |
| Random Effect: Session Variance Component (nested within Mouse) | 3235.8723 | 743.0681 | 4.355 | 0 | 1779.4855 | 4692.2591 | |

**LME — FS [N units = 1209, N mice = 45, Log-Likelihood = -7204.176, Scale(sigma^2) = 7935.57541, Method = REML, Converged = Yes]**

| Term | Beta (Coef.) | SE | z | p | 95% CI Low | 95% CI High | sig |
|---|---|---|---|---|---|---|---|
| Intercept (CT, 12W, reference sex) | 110.5365 | 28.7967 | 3.839 | 0.0001 | 54.0961 | 166.9769 | |
| Age: 15W (vs 12W, reference) | 12.6162 | 35.4863 | 0.356 | 0.7222 | -56.9357 | 82.1681 | ns |
| Age: 26W (vs 12W, reference) | 33.8227 | 33.7718 | 1.002 | 0.3166 | -32.3688 | 100.0142 | ns |
| Group: MP (vs CT, reference) | -29.8608 | 35.4593 | -0.842 | 0.3997 | -99.3597 | 39.6382 | ns |
| Sex: Male (vs Female, reference) | 18.1401 | 25.5913 | 0.709 | 0.4784 | -32.0179 | 68.2981 | ns |
| Interaction: Age 15W x Group MP | 25.9087 | 49.6176 | 0.522 | 0.6016 | -71.34 | 123.1574 | ns |
| Interaction: Age 26W x Group MP | -53.5413 | 44.8475 | -1.194 | 0.2325 | -141.4407 | 34.3581 | ns |
| Random Effect: Mouse Intercept Variance | 780.6174 | 895.9747 | 0.871 | 0.3836 | -975.4607 | 2536.6954 | |
| Random Effect: Session Variance Component (nested within Mouse) | 4412.3731 | 1071.996 | 4.116 | 0 | 2311.2987 | 6513.4475 | |

**LME — Baseline [N units = 3904, N mice = 45, Log-Likelihood = -25110.081, Scale(sigma^2) = 21499.38037, Method = REML, Converged = Yes]**

| Term | Beta (Coef.) | SE | z | p | 95% CI Low | 95% CI High | sig |
|---|---|---|---|---|---|---|---|
| Intercept (CT, 12W, reference sex) | 188.7348 | 46.1746 | 4.087 | 0 | 98.2342 | 279.2353 | |
| Age: 15W (vs 12W, reference) | 9.8733 | 56.7491 | 0.174 | 0.8619 | -101.3529 | 121.0996 | ns |
| Age: 26W (vs 12W, reference) | 38.0414 | 54.2655 | 0.701 | 0.4833 | -68.3171 | 144.4 | ns |
| Group: MP (vs CT, reference) | -21.6212 | 57.084 | -0.379 | 0.7049 | -133.5038 | 90.2613 | ns |
| Sex: Male (vs Female, reference) | 21.2144 | 40.674 | 0.522 | 0.602 | -58.5051 | 100.9339 | ns |
| Interaction: Age 15W x Group MP | 6.9925 | 80.4435 | 0.087 | 0.9307 | -150.6739 | 164.6589 | ns |
| Interaction: Age 26W x Group MP | -53.4946 | 72.2173 | -0.741 | 0.4588 | -195.0378 | 88.0487 | ns |
| Random Effect: Mouse Intercept Variance | 4050.6443 | 2096.117 | 1.932 | 0.0533 | -57.6703 | 8158.959 | |
| Random Effect: Session Variance Component (nested within Mouse) | 8994.5188 | 2005.569 | 4.485 | 0 | 5063.6755 | 12925.362 | |

**LME — Evoked [N units = 3906, N mice = 45, Log-Likelihood = -26276.963, Scale(sigma^2) = 39037.43762, Method = REML, Converged = Yes]**

| Term | Beta (Coef.) | SE | z | p | 95% CI Low | 95% CI High | sig |
|---|---|---|---|---|---|---|---|
| Intercept (CT, 12W, reference sex) | 247.5999 | 58.4297 | 4.238 | 0 | 133.0798 | 362.12 | |
| Age: 15W (vs 12W, reference) | -16.7295 | 71.6847 | -0.233 | 0.8155 | -157.2289 | 123.77 | ns |
| Age: 26W (vs 12W, reference) | 30.4099 | 68.547 | 0.444 | 0.6573 | -103.9398 | 164.7596 | ns |
| Group: MP (vs CT, reference) | -58.6537 | 72.1826 | -0.813 | 0.4165 | -200.1291 | 82.8217 | ns |
| Sex: Male (vs Female, reference) | 21.0182 | 51.165 | 0.411 | 0.6812 | -79.2634 | 121.2997 | ns |
| Interaction: Age 15W x Group MP | 78.4838 | 102.2152 | 0.768 | 0.4426 | -121.8542 | 278.8219 | ns |
| Interaction: Age 26W x Group MP | -0.1239 | 91.2474 | -0.001 | 0.9989 | -178.9654 | 178.7176 | ns |
| Random Effect: Mouse Intercept Variance | 7987.7175 | 3408.184 | 2.344 | 0.0191 | 1307.7995 | 14667.6354 | |
| Random Effect: Session Variance Component (nested within Mouse) | 11462.2258 | 2673.398 | 4.288 | 0 | 6222.4615 | 16701.99 | |

**LME — RS-Baseline [N units = 2696, N mice = 45, Log-Likelihood = -15610.653, Scale(sigma^2) = 5841.58916, Method = REML, Converged = Yes]**

| Term | Beta (Coef.) | SE | z | p | 95% CI Low | 95% CI High | sig |
|---|---|---|---|---|---|---|---|
| Intercept (CT, 12W, reference sex) | 103.3096 | 27.7753 | 3.719 | 0.0002 | 48.871 | 157.7482 | |
| Age: 15W (vs 12W, reference) | -2.5232 | 34.0828 | -0.074 | 0.941 | -69.3243 | 64.2779 | ns |
| Age: 26W (vs 12W, reference) | 7.1779 | 32.5926 | 0.22 | 0.8257 | -56.7024 | 71.0582 | ns |
| Group: MP (vs CT, reference) | -10.3574 | 34.3326 | -0.302 | 0.7629 | -77.648 | 56.9332 | ns |
| Sex: Male (vs Female, reference) | 4.5611 | 24.4238 | 0.187 | 0.8519 | -43.3087 | 52.4308 | ns |
| Interaction: Age 15W x Group MP | 6.6903 | 48.5562 | 0.138 | 0.8904 | -88.478 | 101.8587 | ns |
| Interaction: Age 26W x Group MP | -0.5612 | 43.3788 | -0.013 | 0.9897 | -85.5821 | 84.4597 | ns |
| Random Effect: Mouse Intercept Variance | 1644.1876 | 728.1067 | 2.258 | 0.0239 | 217.1247 | 3071.2506 | |
| Random Effect: Session Variance Component (nested within Mouse) | 2875.9899 | 643.5952 | 4.469 | 0 | 1614.5665 | 4137.4133 | |

**LME — RS-Evoked [N units = 2697, N mice = 45, Log-Likelihood = -16448.061, Scale(sigma^2) = 10919.43690, Method = REML, Converged = Yes]**

| Term | Beta (Coef.) | SE | z | p | 95% CI Low | 95% CI High | sig |
|---|---|---|---|---|---|---|---|
| Intercept (CT, 12W, reference sex) | 127.9521 | 35.2005 | 3.635 | 0.0003 | 58.9603 | 196.9438 | |
| Age: 15W (vs 12W, reference) | -10.6567 | 43.1427 | -0.247 | 0.8049 | -95.2147 | 73.9014 | ns |
| Age: 26W (vs 12W, reference) | 8.5246 | 41.2257 | 0.207 | 0.8362 | -72.2763 | 89.3256 | ns |
| Group: MP (vs CT, reference) | -24.6925 | 43.4698 | -0.568 | 0.57 | -109.8918 | 60.5068 | ns |
| Sex: Male (vs Female, reference) | 3.7269 | 30.8031 | 0.121 | 0.9037 | -56.646 | 64.0998 | ns |
| Interaction: Age 15W x Group MP | 31.6343 | 61.7679 | 0.512 | 0.6085 | -89.4286 | 152.6972 | ns |
| Interaction: Age 26W x Group MP | 35.9986 | 54.879 | 0.656 | 0.5118 | -71.5623 | 143.5595 | ns |
| Random Effect: Mouse Intercept Variance | 3119.0877 | 1205.955 | 2.586 | 0.0097 | 755.4586 | 5482.7167 | |
| Random Effect: Session Variance Component (nested within Mouse) | 3653.0389 | 865.0996 | 4.223 | 0 | 1957.4749 | 5348.603 | |

**LME — FS-Baseline [N units = 1208, N mice = 45, Log-Likelihood = -7018.244, Scale(sigma^2) = 5844.17312, Method = REML, Converged = Yes]**

| Term | Beta (Coef.) | SE | z | p | 95% CI Low | 95% CI High | sig |
|---|---|---|---|---|---|---|---|
| Intercept (CT, 12W, reference sex) | 87.4459 | 25.6479 | 3.409 | 0.0007 | 37.177 | 137.7148 | |
| Age: 15W (vs 12W, reference) | 26.7437 | 31.6586 | 0.845 | 0.3982 | -35.3061 | 88.7935 | ns |
| Age: 26W (vs 12W, reference) | 40.5293 | 30.095 | 1.347 | 0.1781 | -18.4557 | 99.5143 | ns |
| Group: MP (vs CT, reference) | -13.137 | 31.5826 | -0.416 | 0.6774 | -75.0376 | 48.7637 | ns |
| Sex: Male (vs Female, reference) | 16.7319 | 22.8371 | 0.733 | 0.4638 | -28.028 | 61.4917 | ns |
| Interaction: Age 15W x Group MP | -0.0314 | 44.1065 | -0.001 | 0.9994 | -86.4785 | 86.4157 | ns |
| Interaction: Age 26W x Group MP | -61.826 | 39.9616 | -1.547 | 0.1218 | -140.1492 | 16.4972 | ns |
| Random Effect: Mouse Intercept Variance | 522.838 | 714.906 | 0.731 | 0.4646 | -878.3521 | 1924.0281 | |
| Random Effect: Session Variance Component (nested within Mouse) | 3722.4559 | 887.2432 | 4.196 | 0 | 1983.4911 | 5461.4207 | |

**LME — FS-Evoked [N units = 1209, N mice = 45, Log-Likelihood = -7344.599, Scale(sigma^2) = 10048.15473, Method = REML, Converged = Yes]**

| Term | Beta (Coef.) | SE | z | p | 95% CI Low | 95% CI High | sig |
|---|---|---|---|---|---|---|---|
| Intercept (CT, 12W, reference sex) | 128.5598 | 32.2225 | 3.99 | 0.0001 | 65.4048 | 191.7148 | |
| Age: 15W (vs 12W, reference) | 4.9887 | 39.6635 | 0.126 | 0.8999 | -72.7503 | 82.7276 | ns |
| Age: 26W (vs 12W, reference) | 26.7417 | 37.7743 | 0.708 | 0.479 | -47.2946 | 100.778 | ns |
| Group: MP (vs CT, reference) | -41.0575 | 39.6781 | -1.035 | 0.3008 | -118.8252 | 36.7102 | ns |
| Sex: Male (vs Female, reference) | 16.2089 | 28.5747 | 0.567 | 0.5705 | -39.7964 | 72.2142 | ns |
| Interaction: Age 15W x Group MP | 49.3606 | 55.6274 | 0.887 | 0.3749 | -59.6672 | 158.3884 | ns |
| Interaction: Age 26W x Group MP | -44.0628 | 50.17 | -0.878 | 0.3798 | -142.3942 | 54.2685 | ns |
| Random Effect: Mouse Intercept Variance | 1147.2209 | 1122.737 | 1.022 | 0.3069 | -1053.3028 | 3347.7445 | |
| Random Effect: Session Variance Component (nested within Mouse) | 5197.0313 | 1283.157 | 4.05 | 0.0001 | 2682.0896 | 7711.9729 | |

**Figure 6 (Population Coupling):** Model-based simple effect contrasts with Bonferroni correction.

**Post-hoc simple-effects contrasts (CT vs MP) — All (Bonferroni correction across 3 ages)**

| Set | Age (W) | Comparison | Estimate | t_stat | p_raw | p_corrected | sig |
|---|---|---|---|---|---|---|---|
| All | 12 | CT vs MP | -40.8191 | -0.641 | 0.521329 | 1 | ns |
| All | 15 | CT vs MP | 0.6797 | 0.011 | 0.991372 | 1 | ns |
| All | 26 | CT vs MP | -70.128 | -1.425 | 0.154255 | 0.462766 | ns |

**Post-hoc simple-effects contrasts (CT vs MP) — RS (Bonferroni correction across 3 ages)**

| Set | Age (W) | Comparison | Estimate | t_stat | p_raw | p_corrected | sig |
|---|---|---|---|---|---|---|---|
| RS | 12 | CT vs MP | -17.2371 | -0.45 | 0.652485 | 1 | ns |
| RS | 15 | CT vs MP | 0.4718 | 0.012 | 0.990099 | 1 | ns |
| RS | 26 | CT vs MP | -0.4715 | -0.016 | 0.987255 | 1 | ns |

**Post-hoc simple-effects contrasts (CT vs MP) — FS (Bonferroni correction across 3 ages)**

| Set | Age (W) | Comparison | Estimate | t_stat | p_raw | p_corrected | sig |
|---|---|---|---|---|---|---|---|
| FS | 12 | CT vs MP | -29.8608 | -0.842 | 0.399724 | 1 | ns |
| FS | 15 | CT vs MP | -3.9521 | -0.115 | 0.908773 | 1 | ns |
| FS | 26 | CT vs MP | -83.4021 | -3.041 | 0.00236 | **0.007081** | * |

**Post-hoc simple-effects contrasts (CT vs MP) — Baseline (Bonferroni correction across 3 ages)**

| Set | Age (W) | Comparison | Estimate | t_stat | p_raw | p_corrected | sig |
|---|---|---|---|---|---|---|---|
| Baseline | 12 | CT vs MP | -21.6212 | -0.379 | 0.704865 | 1 | ns |
| Baseline | 15 | CT vs MP | -14.6287 | -0.261 | 0.794435 | 1 | ns |
| Baseline | 26 | CT vs MP | -75.1158 | -1.7 | 0.089162 | 0.267487 | ns |

**Post-hoc simple-effects contrasts (CT vs MP) — Evoked (Bonferroni correction across 3 ages)**

| Set | Age (W) | Comparison | Estimate | t_stat | p_raw | p_corrected | sig |
|---|---|---|---|---|---|---|---|
| Evoked | 12 | CT vs MP | -58.6537 | -0.813 | 0.416463 | 1 | ns |
| Evoked | 15 | CT vs MP | 19.8302 | 0.277 | 0.78174 | 1 | ns |
| Evoked | 26 | CT vs MP | -58.7776 | -1.054 | 0.291819 | 0.875457 | ns |

**Post-hoc simple-effects contrasts (CT vs MP) — RS-Baseline (Bonferroni correction across 3 ages)**

| Set | Age (W) | Comparison | Estimate | t_stat | p_raw | p_corrected | sig |
|---|---|---|---|---|---|---|---|
| RS-Baseli | 12 | CT vs MP | -10.3574 | -0.302 | 0.762896 | 1 | ns |
| RS-Baseli | 15 | CT vs MP | -3.6671 | -0.108 | 0.91405 | 1 | ns |
| RS-Baseli | 26 | CT vs MP | -10.9186 | -0.412 | 0.680307 | 1 | ns |

**Post-hoc simple-effects contrasts (CT vs MP) — RS-Evoked (Bonferroni correction across 3 ages)**

| Set | Age (W) | Comparison | Estimate | t_stat | p_raw | p_corrected | sig |
|---|---|---|---|---|---|---|---|
| RS-Evoke | 12 | CT vs MP | -24.6925 | -0.568 | 0.570009 | 1 | ns |
| RS-Evoke | 15 | CT vs MP | 6.9418 | 0.16 | 0.872818 | 1 | ns |
| RS-Evoke | 26 | CT vs MP | 11.3061 | 0.338 | 0.73561 | 1 | ns |

**Post-hoc simple-effects contrasts (CT vs MP) — FS-Baseline (Bonferroni correction across 3 ages)**

| Set | Age (W) | Comparison | Estimate | t_stat | p_raw | p_corrected | sig |
|---|---|---|---|---|---|---|---|
| FS-Baselir | 12 | CT vs MP | -13.137 | -0.416 | 0.677442 | 1 | ns |
| FS-Baselir | 15 | CT vs MP | -13.1684 | -0.43 | 0.666955 | 1 | ns |
| FS-Baselir | 26 | CT vs MP | -74.963 | -3.066 | 0.002171 | **0.006513** | * |

**Post-hoc simple-effects contrasts (CT vs MP) — FS-Evoked (Bonferroni correction across 3 ages)**

| Set | Age (W) | Comparison | Estimate | t_stat | p_raw | p_corrected | sig |
|---|---|---|---|---|---|---|---|
| FS-Evoke | 12 | CT vs MP | -41.0575 | -1.035 | 0.30078 | 0.902339 | ns |
| FS-Evoke | 15 | CT vs MP | 8.3031 | 0.214 | 0.830246 | 1 | ns |
| FS-Evoke | 26 | CT vs MP | -85.1203 | -2.775 | 0.005523 | **0.016568** | * |

**Figure S7 (LFPs):** LME Models for predicting Layer IV LFP power, and normalized band powers.

**LME — Layer IV Baseline AUC [N obs = 84, N groups = 45, Log-Likelihood = -737.445, Scale(sigma^2) = 4267677.065343, Method = REML, Converged = Yes]**

| Term | Beta (Coef.) | SE | z | p | 95% CI Low | 95% CI High | sig |
|---|---|---|---|---|---|---|---|
| Intercept (CT, 12WK, reference sex) | 5648.6788 | 1678.156 | 3.366 | 0.0008 | 2359.5537 | 8937.804 | |
| Age: 15WK (vs 12WK, reference) | -617.6653 | 2046.238 | -0.302 | 0.7628 | -4628.2182 | 3392.8877 | ns |
| Age: 26WK (vs 12WK, reference; pools former 24WK+28WK animals) | -1708.0453 | 1964.646 | -0.869 | 0.3846 | -5558.6799 | 2142.5893 | ns |
| Group: MP (vs CT, reference) | 143.3547 | 2077.877 | 0.069 | 0.945 | -3929.2096 | 4215.9191 | ns |
| Sex: Male (vs Female, reference) | -246.0915 | 1433.104 | -0.172 | 0.8637 | -3054.9234 | 2562.7405 | ns |
| Interaction: Age 15WK x Group MP | 615.6959 | 2953.611 | 0.208 | 0.8349 | -5173.2752 | 6404.667 | ns |
| Interaction: Age 26WK x Group MP | -1210.293 | 2623.27 | -0.461 | 0.6445 | -6351.8086 | 3931.2225 | ns |
| Random Effect: Mouse Intercept Variance | 2.3601 | 0.9305 | 2.537 | 0.0112 | 0.5365 | 4.1838 | |

**LME — Beta Band - Baseline [N obs = 84, N groups = 45, Log-Likelihood = 165.374, Scale(sigma^2) = 0.000469, Method = REML, Converged = Yes]**

| Term | Beta (Coef.) | SE | z | p | 95% CI Low | 95% CI High | sig |
|---|---|---|---|---|---|---|---|
| Intercept (CT, 12WK, reference sex) | 0.2457 | 0.0103 | 23.755 | 0 | 0.2254 | 0.2659 | |
| Age: 15WK (vs 12WK, reference) | 0.0215 | 0.0127 | 1.701 | 0.089 | -0.0033 | 0.0463 | ns |
| Age: 26WK (vs 12WK, reference; pools former 24WK+28WK animals) | 0.0035 | 0.0122 | 0.29 | 0.7717 | -0.0204 | 0.0275 | ns |
| Group: MP (vs CT, reference) | 0.0191 | 0.0128 | 1.489 | 0.1366 | -0.006 | 0.0442 | ns |
| Sex: Male (vs Female, reference) | 0.0007 | 0.0091 | 0.072 | 0.9427 | -0.0171 | 0.0184 | ns |
| Interaction: Age 15WK x Group MP | -0.0351 | 0.018 | -1.946 | 0.0516 | -0.0704 | 0.0002 | ns |
| Interaction: Age 26WK x Group MP | 0.0027 | 0.0162 | 0.168 | 0.8666 | -0.0291 | 0.0346 | ns |
| Random Effect: Mouse Intercept Variance | 0.4568 | 0.3199 | 1.428 | 0.1533 | -0.1702 | 1.0838 | |

**LME — Gamma Band - Baseline [N obs = 84, N groups = 45, Log-Likelihood = 175.716, Scale(sigma^2) = 0.000334, Method = REML, Converged = Yes]**

| Term | Beta (Coef.) | SE | z | p | 95% CI Low | 95% CI High | sig |
|---|---|---|---|---|---|---|---|
| Intercept (CT, 12WK, reference sex) | 0.0859 | 0.0094 | 9.133 | 0 | 0.0674 | 0.1043 | |
| Age: 15WK (vs 12WK, reference) | 0.0028 | 0.0115 | 0.241 | 0.8098 | -0.0197 | 0.0253 | ns |
| Age: 26WK (vs 12WK, reference; pools former 24WK+28WK animals) | 0.006 | 0.0111 | 0.539 | 0.5896 | -0.0157 | 0.0277 | ns |
| Group: MP (vs CT, reference) | 0.0005 | 0.0116 | 0.042 | 0.9668 | -0.0223 | 0.0233 | ns |
| Sex: Male (vs Female, reference) | 0.0011 | 0.0082 | 0.134 | 0.8931 | -0.015 | 0.0172 | ns |
| Interaction: Age 15WK x Group MP | -0.0065 | 0.0164 | -0.395 | 0.6931 | -0.0386 | 0.0256 | ns |
| Interaction: Age 26WK x Group MP | 0.0087 | 0.0147 | 0.593 | 0.5529 | -0.0201 | 0.0376 | ns |
| Random Effect: Mouse Intercept Variance | 0.6128 | 0.3828 | 1.601 | 0.1094 | -0.1375 | 1.363 | |

**LME — Beta Band - Evoked (40mN) [N obs = 84, N groups = 45, Log-Likelihood = 16.761, Scale(sigma^2) = 0.025107, Method = REML, Converged = Yes]**

| Term | Beta (Coef.) | SE | z | p | 95% CI Low | 95% CI High | sig |
|---|---|---|---|---|---|---|---|
| Intercept (CT, 12WK, reference sex) | 0.391 | 0.0667 | 5.859 | 0 | 0.2602 | 0.5218 | |
| Age: 15WK (vs 12WK, reference) | 0.1153 | 0.0819 | 1.408 | 0.159 | -0.0451 | 0.2757 | ns |
| Age: 26WK (vs 12WK, reference; pools former 24WK+28WK animals) | -0.0215 | 0.0788 | -0.273 | 0.7848 | -0.176 | 0.133 | ns |
| Group: MP (vs CT, reference) | 0.0597 | 0.0828 | 0.722 | 0.4703 | -0.1025 | 0.2219 | ns |
| Sex: Male (vs Female, reference) | 0.0329 | 0.059 | 0.558 | 0.577 | -0.0828 | 0.1486 | ns |
| Interaction: Age 15WK x Group MP | -0.1031 | 0.1156 | -0.892 | 0.3722 | -0.3296 | 0.1234 | ns |
| Interaction: Age 26WK x Group MP | 0.0802 | 0.105 | 0.764 | 0.4451 | -0.1256 | 0.2859 | ns |
| Random Effect: Mouse Intercept Variance | 0.2382 | 0.2208 | 1.079 | 0.2807 | -0.1945 | 0.6709 | |

**LME — Gamma Band - Evoked (40mN) [N obs = 84, N groups = 45, Log-Likelihood = 79.114, Scale(sigma^2) = 0.003922, Method = REML, Converged = Yes]**

| Term | Beta (Coef.) | SE | z | p | 95% CI Low | 95% CI High | sig |
|---|---|---|---|---|---|---|---|
| Intercept (CT, 12WK, reference sex) | 0.1207 | 0.0337 | 3.581 | 0.0003 | 0.0546 | 0.1868 | |
| Age: 15WK (vs 12WK, reference) | 0.077 | 0.0412 | 1.867 | 0.0619 | -0.0038 | 0.1578 | ns |
| Age: 26WK (vs 12WK, reference; pools former 24WK+28WK animals) | 0.0347 | 0.0397 | 0.875 | 0.3817 | -0.043 | 0.1124 | ns |
| Group: MP (vs CT, reference) | 0.0194 | 0.0418 | 0.464 | 0.6426 | -0.0625 | 0.1013 | ns |
| Sex: Male (vs Female, reference) | 0.0025 | 0.0293 | 0.085 | 0.9322 | -0.055 | 0.06 | ns |
| Interaction: Age 15WK x Group MP | -0.0327 | 0.059 | -0.555 | 0.5792 | -0.1483 | 0.0829 | ns |
| Interaction: Age 26WK x Group MP | 0.0195 | 0.0529 | 0.37 | 0.7117 | -0.0841 | 0.1231 | ns |
| Random Effect: Mouse Intercept Variance | 0.7287 | 0.384 | 1.897 | 0.0578 | -0.0241 | 1.4814 | |

**Figure S7 (LFPs):** model-based simple effects of genotype (MP - CT) at each age, with Bonferroni corrections.

**Post-hoc simple-effects contrasts (CT vs MP) — Layer IV Baseline AUC (Bonferroni correction across 3 ages)**

| Measure | Age (WK) | Comparison | Estimate (μV²) | t_stat | 95% CI Low | 95% CI High | p_raw | p_corrected | sig |
|---|---|---|---|---|---|---|---|---|---|
| Layer IV Baseline AUC | 12 | CT vs MP | 143.35475 | 0.069 | -3929.20959 | 4215.91908 | 0.944997 | 1 | ns |
| Layer IV Baseline AUC | 15 | CT vs MP | 759.05062 | 0.367 | -3295.17514 | 4813.27637 | 0.713654 | 1 | ns |
| Layer IV Baseline AUC | 26 | CT vs MP | -1066.93829 | -0.667 | -4203.07167 | 2069.1951 | 0.504903 | 1 | ns |

**Post-hoc simple-effects contrasts (CT vs MP) — Beta Band - Baseline (Bonferroni correction across 3 ages)**

| Measure | Age (WK) | Comparison | Estimate (norm. power) | t_stat | 95% CI Low | 95% CI High | p_raw | p_corrected | sig |
|---|---|---|---|---|---|---|---|---|---|
| Beta Band - Baseline | 12 | CT vs MP | 0.01908 | 1.489 | -0.00604 | 0.0442 | 0.136578 | 0.409734 | ns |
| Beta Band - Baseline | 15 | CT vs MP | -0.01602 | -1.269 | -0.04075 | 0.00872 | 0.204412 | 0.613235 | ns |
| Beta Band - Baseline | 26 | CT vs MP | 0.02181 | 2.19 | 0.00229 | 0.04132 | 0.028516 | 0.085549 | ns |

**Post-hoc simple-effects contrasts (CT vs MP) — Gamma Band - Baseline (Bonferroni correction across 3 ages)**

| Measure | Age (WK) | Comparison | Estimate (norm. power) | t_stat | 95% CI Low | 95% CI High | p_raw | p_corrected | sig |
|---|---|---|---|---|---|---|---|---|---|
| Gamma Band - Baseline | 12 | CT vs MP | 0.00049 | 0.042 | -0.02233 | 0.0233 | 0.966769 | 1 | ns |
| Gamma Band - Baseline | 15 | CT vs MP | -0.00598 | -0.524 | -0.02835 | 0.01639 | 0.600406 | 1 | ns |
| Gamma Band - Baseline | 26 | CT vs MP | 0.00923 | 1.023 | -0.00845 | 0.0269 | 0.306293 | 0.91888 | ns |

**Post-hoc simple-effects contrasts (CT vs MP) — Beta Band - Evoked (40mN) (Bonferroni correction across 3 ages)**

| Measure | Age (WK) | Comparison | Estimate (norm. power) | t_stat | 95% CI Low | 95% CI High | p_raw | p_corrected | sig |
|---|---|---|---|---|---|---|---|---|---|
| Beta Band - Evoked (40mN) | 12 | CT vs MP | 0.05975 | 0.722 | -0.10246 | 0.22195 | 0.470335 | 1 | ns |
| Beta Band - Evoked (40mN) | 15 | CT vs MP | -0.04338 | -0.542 | -0.20031 | 0.11356 | 0.587997 | 1 | ns |
| Beta Band - Evoked (40mN) | 26 | CT vs MP | 0.1399 | 2.167 | 0.01335 | 0.26645 | 0.030257 | 0.090771 | ns |

**Post-hoc simple-effects contrasts (CT vs MP) — Gamma Band - Evoked (40mN) (Bonferroni correction across 3 ages)**

| Measure | Age (WK) | Comparison | Estimate (norm. power) | t_stat | 95% CI Low | 95% CI High | p_raw | p_corrected | sig |
|---|---|---|---|---|---|---|---|---|---|
| Gamma Band - Evoked (40mN) | 12 | CT vs MP | 0.01939 | 0.464 | -0.06248 | 0.10126 | 0.642574 | 1 | ns |
| Gamma Band - Evoked (40mN) | 15 | CT vs MP | -0.01333 | -0.323 | -0.09413 | 0.06747 | 0.746453 | 1 | ns |
| Gamma Band - Evoked (40mN) | 26 | CT vs MP | 0.03893 | 1.202 | -0.02453 | 0.10238 | 0.229255 | 0.687764 | ns |
